# A baseline for aquatic parasite diversity in West African mangroves: host-parasite records, DNA barcodes, and an experimental design for future use

**DOI:** 10.64898/2026.10.30.762766

**Authors:** Armando J. Cruz-Laufer, Lucas Van Passel, Oumar Sadio, Farid Dahdouh-Guebas, Ralph Appy, James P. Bernot, Chahinez Bouguerche, Arfang Diamanka, Jesús Hernández-Orts, Nikol Kmentová, Edgar F. Mendoza-Franco, Camila Pantoja, Miriam I. Shigoley, Njord Vangelder, Chelsea L. Wood, Maarten P. M. Vanhove

**Affiliations:** UHasselt – Hasselt University, Centre for Environmental Sciences, Research Group Zoology: Biodiversity & Toxicology, 3590 Diepenbeek, Belgium; Systems Ecology and Resource Management Research Unit (SERM), Université libre de Bruxelles-ULB, Brussels, Belgium; Hogeschool PXL – PXL BIO-Research, 3590 Diepenbeek, Belgium; IRD, Univ Brest, CNRS, Ifremer, LEMAR, BP 1386 Dakar, Senegal; bDIV: Ecology, Evolution & Genetics, Biology Department, Vrije Universiteit Brussel - VUB, Brussels, Belgium; Mangrove Specialist Group (MSG), Species Survival Commission (SSC), International Union for the Conservation of Nature (IUCN), Zoological Society of London, London, United Kingdom; Cabrillo Marine Aquarium, 3720 Stephen M. White Drive, San Pedro, CA, 90731, USA; Department of Biology, California State University at Long Beach, 1250 Bellflower Blvd., Long Beach, CA, 90840, USA; Department of Ecology and Evolutionary Biology, University of Connecticut, Storrs, CT, USA; Department of Invertebrate Zoology, Smithsonian National Museum of Natural History, Washington, DC, USA; Department of Zoology, Swedish Museum of Natural History, Stockholm, Sweden; Laboratory of Parasitology, Department of Animal Biology, Faculty of Sciences and Technologies, University Cheikh Anta Diop of Dakar, BP 5005, Dakar, Senegal; Department of Invertebrates, Natural History Museum of Geneva, Geneva, Switzerland; Institute of Parasitology, Biology Centre, Czech Academy of Sciences, České Budějovice, Czech Republic; Department of Life Sciences, Natural History Museum, London, UK; Freshwater Biology, Operational Directorate Natural Environment, Royal Belgian Institute of Natural Sciences, Brussels, Belgium; Instituto de Ecología, Pesquerías y Oceanografía del Golfo de México (EPOMEX), Universidad Autónoma de Campeche, Campeche, Mexico; Department of Veterinary Management of Animal Resources, Faculty of Veterinary Medicine, Liège University, Liège, Belgium; School of Aquatic and Fishery Sciences, University of Washington, Seattle, Washington, USA

**Keywords:** DNA reference library, fisheries impact, macroparasites, marine protected area, Senegal

## Abstract

Macroparasites represent a major fraction of biodiversity and biomass in marine ecosystems, yet remain substantially underrepresented in ecological surveys and food-web models, particularly in the Global South. This study addresses this geographic bias by providing the first comprehensive ecological survey of macroparasite communities of fishes in a West African Marine Protected Area (MPA) and adjacent Open-Access Area (OAA) characterised by mangrove forests. We collected 5,020 parasite specimens from 10 fish and one crab species in the Saloum Delta, Senegal, identifying 27 parasite species across all major taxa. Our findings include 15 new records for Senegal of which 11 are also new for West Africa. Additionally, 64% of host-parasite combinations constitute new host records. We provide 208 DNA barcodes, offering the first molecular data for 31% of these species globally and establishing a reference library for future DNA metabarcoding efforts in the region. Using sample size–corrected biodiversity indices and mixed-model analyses, we detected significant variations in parasite infracommunities and component communities between the MPA and OAA, demonstrating the feasibility of using comparative study designs to monitor fisheries impacts on parasite diversity in the Saloum Delta. Furthermore, the identification of trophically transmitted parasites provided evidence of unstudied food-web connections, e.g. predator-prey linkages between fishes and higher-trophic-level definitive hosts including birds and elasmobranchs. By integrating morphological identification, molecular data, and archiving of voucher specimens into publicly accessible, curated collections, this study establishes a new baseline for marine parasite biodiversity in West Africa and provides a methodological blueprint for surveying understudied regions.

## Introduction

Marine fishes host a diverse assemblage of different groups of macroparasites, including flatworms, roundworms, acanthocephalans, and arthropods (Rohde 2002). Macroparasites can account for 17–33% of total host-parasite biomass (Lambden and Johnson 2013; Preston et al. 2013) and 40% of species in an ecosystem (Dobson et al. 2008). By comprising a substantial portion of community biomass and biodiversity, parasitic organisms increase linkage in ecological communities (Lafferty et al. 2006; McLaughlin et al. 2020; Morton and Lafferty 2022) and thus potentially increase the stability of food webs against environmental disturbance (Landi et al. 2018). However, despite their ubiquity and relevance, macroparasites are rarely considered in studies assembling marine food webs (McLaughlin et al. 2020).

The scientific literature on the ecology of parasite communities remains particularly limited for Global South countries. While some food web studies feature South African and Mexican marine parasites (McLaughlin et al. 2020), the biodiversity of parasites in (sub-)tropical regions remains largely unknown (Carlson et al. 2020; Pappalardo et al. 2020). For instance, the African continent contains several of the world’s biodiversity hotspots (Myers et al. 2000), yet most parasite species in this region are probably undescribed (Jorge and Poulin 2018).

Even known wildlife parasites often lack baseline data about their population size (Wood et al. 2026), genetic diversity (Poulin et al. 2019), life cycles (Bennett et al. 2023), and host repertoires (Wells and Clark 2019). Limited funding opportunities in many Global South countries (Vanhove et al. 2017; Sánchez Herrera et al. 2024; Hayah et al. 2025) may further exacerbate this imbalance. Although progress has been made, e.g., with identification guides for the parasites of African freshwater fishes (Scholz et al. 2018b), ecological surveys targeting marine parasite communities, specifically along the coast of sub-Saharan Africa, remain scarce.

Macroparasite diversity is threatened by anthropogenic pressures including global warming, habitat degradation, and pollution (Marcogliese 2023). Another major threat to marine biodiversity is overfishing (Arthington et al. 2016; Mazaris et al. 2019), which in some cases has been linked to the extinction of parasite species (MacKenzie and Pert 2018). But how do parasite communities react to fisheries pressure, i.e., the removal of their host fishes from the environment? Several direct and indirect effects may influence parasite populations at each development life stage (Wood et al. 2010). Host population density, fragmentation of host populations, selective removal of infected or uninfected hosts, a reduction of food web complexity, absence of predators, competitors, or prey may alter how parasite species are impacted by human activity (see detailed review in Wood et al. 2010). Notably, while several studies have attempted to test the effect of fisheries on parasite populations (Wood et al. 2014; MacKenzie and Pert 2018; Henriksen et al. 2019), the majority of these focus on the Global North. Tropical and subtropical marine parasites are heavily underrepresented, even for commercially relevant fishes (Reed 2015).

We aim to address this geographic bias by surveying marine parasite communities in a mangrove ecosystem in sub-Saharan Africa. Mangrove forests are a widely distributed ecosystem across tropical and subtropical coastlines (Bunting et al. 2022). They provide ecosystem services to neighbouring communities such as timber and firewood (Satyanarayana et al. 2021; Dahdouh-Guebas et al. 2026), disease vector control (Griffin and Knight 2012; Cruz-Laufer et al. 2026a, 2026b), coastline stabilisation (Barbier et al. 2011; Strain et al. 2022), tourism (Spalding and Parrett 2019), and carbon storage (Wolswijk et al. 2026). Mangrove fisheries are frequently cited as an important ecosystem service and are estimated to support 4.1 million fishers worldwide (zu Ermgassen et al. 2021, 2025). At the same time, overfishing is one of the major threats to fish assemblages in mangrove forests (Sadio et al. 2015; Reis-Filho et al. 2019; Rasoanirina et al. 2026). This threat may spill over into other ecosystems as mangrove forests serve as nurseries for an abundance of juvenile marine fishes (zu Ermgassen et al. 2025).

Our study provides a first survey of macroparasites in a marine protected area (MPA) and adjacent open-access area (OAA) in West Africa. The Bamboung MPA in the Saloum Delta in Senegal has previously been used to study the effects of total fishery closure in mangrove forests (Sadio et al. 2015; Thiaw et al. 2021). We provide morphological and molecular data on the parasites infecting eight target fishes in this ecosystem, alongside photographic evidence. Furthermore, we tested the use of statistical tools for biodiversity analysis including biodiversity indices and mixed model analysis to detect variations in parasite communities at the level of parasite infracommunities (within-host communities), component communities (host population–specific communities), and host-parasite interaction networks, providing a first insight into the structure and function of parasite communities in the region.

## Material and methods

### Sample sites and sample collection

In September 2022, fishes were collected in the Saloum Delta, Senegal, during an exploratory expedition to survey their parasite communities. We chose this region because the impact of fishing bans on the assemblages of fishes has been studied extensively in the past. Notably, comparisons between the Bamboung Marine Protected Area (MPA), where fishing is banned, and the nearby Bolong de Sangako – an open-access area (OAA) – show that fishing bans have led to an increase in biomass of piscivorous fishes (Sadio et al. 2015; Thiaw et al. 2021). We adopted this comparative approach (MPA *vs.* OAA) for our parasitological study. Fishes were collected by local fishers using beach seine nets in five locations: three in the MPA and two in the OAA (Fig. 1A–C: Table 1). Additionally, mudskippers, *Periophthalmus barbarus* (L., 1766), were caught by hand with the help of volunteers in the village of Néma Ba, a third OAA site. Geographical coordinates and sampling dates are listed in Table 1.

**Figure 1.**
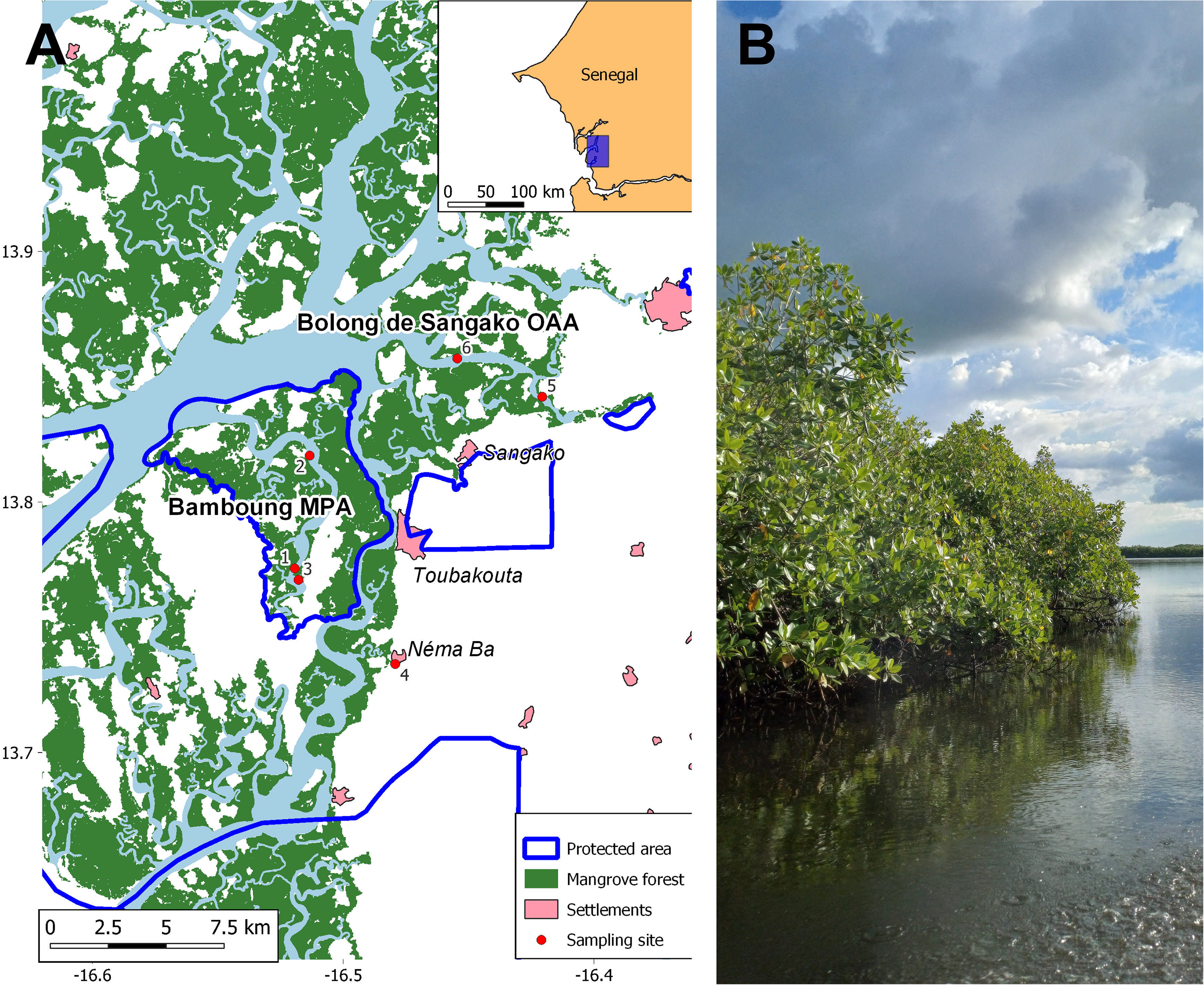
Overview of sampling sites: **A** sampling locations (points) in the Saloum Delta including the Bamboung Marine Protected Area (MPA), the Bolong de Sangako Open-Access Area (OAA), and a streamlet in the village of Néma Ba, with 2020 mangrove land cover (raster layer, green) (Bunting et al. 2022), extent of protected areas (boundaries, blue), settlements (polygons, red), and sampling sites (points, red; numbers refer to Table 1). **B** Main tidal channel in the Bamboung MPA surrounded by mangrove trees (*Avicennia* sp.).

**Table 1.** List of sampling sites along the Saloum Delta, Senegal (see Figure 1). Abbreviations: MPA, Bamboung Marine Protected Area; OAA, open-access area of Bolong de Sangako.

| **Name** | **Site ID** | **Status** | **Date** | **Latitude** | **Longitude** |
|---|---|---|---|---|---|
| Bamboung | 1 | MPA | 11/09/2022 | 13.77344 | -16.51935 |
| Bamboung | 2 | MPA | 12/09/2022 | 13.81845 | -16.51338 |
| Bamboung | 3 | MPA | 10/09/2022 | 13.76888 | -16.51783 |
| Village of Néma Ba | 4 | OAA | 14/09/2022 | 13.73528 | -16.47925 |
| Bolong de Sangako | 5 | OAA | 13/09/2022 | 13.84193 | -16.42060 |
| Bolong de Sangako | 6 | OAA | 14/09/2022 | 13.85722 | -16.45450 |

The sampling design of the campaign was opportunistic, aiming to maximise host diversity. Fishes and crabs were dissected on site. The gills and gastrointestinal tract of fishes as well as the soft-body parts of the crabs were stored in absolute ethanol at 4°C. Among these samples, 129 individuals belonging to 13 fish species and one crab species were screened for macroparasites (Table 2). Eight fish species were considered target species: *Coptodon guineensis* (Günther, 1862) (Cichliformes: Cichlidae), *Caranx hippos* (L., 1766) (Carangiformes: Carangidae), *Ethmalosa fimbriata* (Bowdich, 1825) (Clupeiformes: Dorosomatidae), *Gerres nigri* Günther, 1859 (Acanthuriformes: Gerreidae), *Hemichromis fasciatus* Peters, 1857 (Cichliformes: Cichlidae), *Ilisha africana* (Bloch, 1795) (Clupeiformes: Pristigasteridae), *Monodactylus sebae* (Cuvier, 1829) (Acanthuriformes: Monodactylidae), *Periophthalmus barbarus* (L., 1766) (Gobiiformes: Oxudercidae), and *Sarotherodon melanotheron* Rüppell, 1859 (Cichliformes: Cichlidae). These fish species were selected due to their presence in sufficient numbers both in the Bamboung MPA and the Bolong de Sangako OAA (Sadio et al. 2015; Thiaw et al. 2021) and among the collected tissue samples (Table 2). We specifically targeted gills and the gastrointestinal tract (referred to as “gut” in the following) as they are reportedly the organs where most parasite individuals are found (Justine et al. 2012). The target species include arthropodivorous (*C. hippos*, *G. nigri*, *H. fasciatus*, *M. sebae*, *P. barbarus*), omnivorous (*C. guineensis, S. melanotheron*), and planktivorous (*E. fimbriata*, *I. africana*) fishes (Froese and Pauly 2025).

**Table 2.** Biological data for the fishes examined for microparasites from Saloum Delta, Senegal (see Figure 1 for details on the sampling sites). The standard length (SL) of the fishes is provided in cm.

| **Species** | **Site 1 (MPA)** |  | **Site 2 (MPA)** |  | **Site 3 (MPA)** |  | **Site 4 (OAA)** |  | **Site 5 (MPA)** |  | **Site 6 (OAA)** |  |
|---|---|---|---|---|---|---|---|---|---|---|---|---|
|  | **n** | **SL** | **n** | **SL** | **n** | **SL** | **n** | **SL** | **n** | **SL** | **n** | **SL** |
| Acanthuriformes: Gerreidae<br>*Gerres nigri* Günther | 4 | 11.5–12.8 | 4 | 12.5–15.0 | 3 | 13.0–17.1 | – | – | 2 | 12.5–13.4 | 4 | 12.7–17.7 |
| Acanthuriformes: Monodactylidae<br>*Monodactylus sebae* (Cuvier) | 5 | 4.5–14.5 | 3 | 8.0–8.1 | 3 | 9.4–10.6 | – | – | 2 | 10.7–11.4 | 2 | 10.2–11.2 |
| Carangiformes: Carangidae<br>*Caranx hippos* (L.) | – | – | 5 | 11.6–13.2 | 1 | 13.5 | – | – | – | – | 1 | 13.6 |
| Carangiformes: Polynemidae<br>*Polydactylus quadrifilis* (Cuvier) | 5 | 38.7–62.2 | – | – | 2 | 25.8–32.0 | – | – | – | – | – | – |
| Carangiformes: Sphyraenidae<br>*Sphyraena afra* Peters | 3 | 47.0–54.0 | 2 | 29.7–33.7 | – | – | – | – | 2 | 43.4–50.7 | 1 | 35.5 |
| Cichliformes: Cichlidae<br>*Coptodon guineensis* (Günther) | 1 | 20.6 | 1 | 12.2 | 2 | 20.8–22.0 | – | – | – | – | 3 | 13.1–17.9 |
| *Hemichromis fasciatus* Peters | 2 | 18.5–19.5 | 3 | 15.9–18.2 | – | – | – | – | – | – | 1 | 16.3 |
| *Sarotherodon melanotheron* Rüppell | 3 | 17.7–21.2 | 5 | 10.5–15.9 | – | – | – | – | – | – | 1 | 16.3 |
| Clupeiformes: Dorosomatidae<br>*Ethmalosa fimbriata* (Bowdich) | 1 | 7.3 | 3 | 16.1 | 7 | 13.2–15.7 | – | – | – | – | 1 | 15.6 |
| Clupeiformes: Pristigasteridae<br>*Ilisha africana* (Bloch) | – | – | – | – | 1 | 15 | – | – | – | – | 1 | 12.8 |
| Elopiformes: Elopidae<br>*Elops lacerta* Valenciennes | 3 | 35.2–43.4 | 3 | 26.4–27.6 | – | – | – | – | 2 | 36.3–38.8 | 1 | 24.8 |
| Gobiiformes: Oxudercidae<br>*Periophthalmus barbarus* (L.) | 1 | 5.4 | – | – | – | – | 21 | 3.8–15.2 | – | – | – | – |
| Perciformes: Epinephelidae<br>*Epinephelus aeneus* (Geoffroy) | – | – | – | – | 4 | 19.9–30.8 | – | – | – | – | 1 | 22 |

Furthermore, we opportunistically explored the parasites of piscivorous host species *Elops lacerta* Valenciennes, 1847 (Elopiformes: Elopidae), *Epinephelus aeneus* (Geoffroy Saint-Hilaire, 1817) (Perciformes: Epinephelidae), *Polydactylus quadrifilis* (Cuvier, 1829) (Carangiformes: Polynemidae), and *Sphyraena afra* Peters, 1844 (Carangiformes: Sphyraenidae). However, we only focused on the parasite species infecting the gills of those fishes due to time limitations. Populations of these piscivores are the most affected by fisheries pressure in the Saloum Delta (Sadio et al. 2015; Thiaw et al. 2021); therefore, comparisons of the associated parasite communities are limited by the low host numbers available in the OAA. Nonetheless, we incorporated the data of piscivore-infecting parasites into the present study for their valuable new geographical and host records.

We also screened the soft body parts (muscle and connective tissues, internal organs, gills) of three swimming crabs, *Callinectes amnicola* Rochebrune, 1883 (Decapoda: Portunidae), as these were regularly caught in the same nets. These crabs were collected in a follow-up expedition in September 2024 using the same sampling techniques and slightly different sites in the MPA (geocoordinates: 13.77058, −16.52145) and OAA (geocoordinates: 13.83358, - 16.41452).

### Parasite screening and identification

Fish guts and gills and the soft-body parts of the crabs were dissected on site and stored in absolute (≥ 99.8%) ethanol at 4°C. The tissues were screened later under a stereomicroscope using entomological needles and tweezers, while remaining submerged in ethanol to minimise DNA degradation. All samples were screened twice by the same authors in alternating order (AJCL and LVP) to standardise sampling effort during the screening process.

The treatment of the parasite specimens depended on the taxonomic group, but we obtained tissue for at least one specimen of each parasite species. Some of the monopisthocotylan, polyopisthocotylan, trematode, cestode, and acanthocephalan specimens were cut prior to mounting, with the posterior portion used for DNA barcoding. Other specimens were mounted whole. Monopisthocotylans and some polyopisthocotylans were mounted in Hoyer’s medium (Anderson 1954), left to dry for 1–2 weeks prior to sealing the slides. Trematodes, cestodes, and some polyopisthocotylans were stained with borax carmine following the approach of Pantoja et al. (2022). Specimens were cleared in lactophenol (up to 15 min), rinsed in 70% ethanol (30 min), hydrated in 30% ethanol and deionised water (5 min each), stained in aqueous borax carmine (3 h), rinsed in deionised water (twice 5 min), and dehydrated in a graded ethanol series of 70% (5 min), 80% (5 min), 95% (5 min), and 100% (5 min). Finally, specimens were cleared with clove oil and mounted on Euparal (Chroma-Gesellschaft Schmid, Stuttgart, Germany) for permanent storage and dried for 48 h at 50°C. For acanthocephalan specimens, we followed the same procedure but, in the clearing steps, we used methyl salicylate. Nematode larvae were first cut into three pieces with the anterior and posterior portions kept for morphological identification and the remaining body used for DNA barcoding. Heads and tails were cleared in lactophenol until appearing transparent (15– 30 min), before mounting these pieces in glycerine. Copepods were cleared enzymatically with proteinase K (30 mins, 55°C) in a DNA extraction buffer solution (see ‘Molecular reference database’) before mounting in glycerine. The remaining buffer solution was kept for DNA barcoding.

One nematode specimen (*Echinocephalus* sp.) was also imaged using scanning electron microscopy, as outlined in Shigoley et al. (2025). The specimen was post-fixed in 4% OsO (osmium tetroxide), rinsed in distilled water to remove excess osmium tetroxide, and dehydrated through a graded ethanol series of 30%, 50%, 70% (three times), 80%, 95%, and 100% (three times), with each step lasting 30 min. Specimens were subsequently dried overnight under a fume hood using hexamethyldisilazane. Dried specimens were mounted on aluminium SEM stubs using double-sided adhesive tape and sputter-coated with gold at 30 mA using a JEOL JFC-1300 sputter coater (JEOL Ltd., Tokyo, Japan), following (Shigoley et al. 2025). Imaging was performed using a Phenom XL G2 Desktop scanning electron microscope (Thermo Fisher Scientific, Waltham, MA, USA) at an accelerating voltage of 5 kV.

All morphological vouchers were sealed with Glyceel and deposited in the Zoological Collection of Hasselt University (Diepenbeek, Belgium) (HU XXX–XXX), the Muséum National d’Histoire Naturelle (Paris, France) (MNHN HELXXXX–XXXX), and the MNHN collection at the Institut Scientifique de Rabat (Rabat, Morocco) (MNHN XXXXXX). Specimen photographs collected for this study were deposited on iNaturalist (project ID: 300535, URL: https://www.inaturalist.org/projects/300535) and MorphoBank (XXXXXXX).

### Parasite identification

We identified parasites using an Olympus BX53 light microscope (Evident Corporation, Tokyo, Japan) equipped with both differential interference contrast (DIC) and phase contrast (PhC), with the latter being applied to visualise the hard parts of attachment and reproductive organs of monopisthocotylan and polyopisthocotylan flatworms and the chitin skeleton of parasitic copepods.

Parasites were assigned to major taxonomic groups (Eoacanthocephala, Cestoda, Copepoda, Monopisthocotyla, Chromadorea, Polyopisthocotyla, and Trematoda) using the key of Kuchta et al. (2018). For further identification, we used taxon-specific identification keys, generic diagnoses, and species descriptions (see *Results*). For parasite specimens belonging to hitherto undescribed species, we assigned temporary names (cheironyms), pending the formal descriptions of these species. Undescribed species were delimited using the same morphological characters as for identifications, where present. Otherwise, we used molecular methods for delimiting species (see below).

Some parasite specimens could not be identified through morphological characters alone because of the lack of some of these characters in larval stages or in specimens cut for DNA barcoding. This limitation applied to trematodes and larval nematodes. Accordingly, we supplemented the morphological identification with molecular identifications through DNA barcoding (see ‘Molecular reference database’). Specifically, we used NCBI BLAST (Camacho et al. 2009) to match sequences with sequences available on GenBank (https://www.ncbi.nlm.nih.gov/genbank/).

We considered sequences with >99% nucleotide identity in the internal transcribed spacer 1 (ITS1) and 2 (ITS2) and 98% for cytochrome *c* oxidase I (*cox*1) to be conspecific. Following previous studies on flatworms, nematodes, and copepods (Moszczynska et al. 2009; Baek et al. 2016; Poon et al. 2017), these thresholds were set conservatively high to minimise the risk of misidentifications. This precaution is necessary because barcoding gaps can vary substantially among closely related species (Vanhove et al. 2013; Kmentová et al. 2020; Decru et al. 2022). We applied the same criteria to discern whether specimens that we could not identify to species level morphologically (specifically nematodes and trematodes) belonged to the same species. Above 97%, we assigned the same genus names (but see ‘Results’ for detailed considerations on morphological and molecular identifications). We recognise that these molecular identifications, particularly at the genus level, will have to be verified in future studies through more extensive analysis of the morphological characters, but these IDs provide a reliable approximation of the species identity, suitable for the present ecological study.

### Molecular reference database

Apart from some nematode and monopisthocotylan species (see ‘Results’), many of the parasite species we found had never been sequenced before. For those species that had been sequenced, no molecular data were available from Senegal. Therefore, we sequenced the DNA barcoding markers for all collected parasite species, focusing on the markers that have been most widely applied to species belonging to the same genus or family.

We performed DNA extractions according to the protocol published by Kmentová et al. (2021), using enzymatic digestion through proteinase K and DNA precipitation through ethanol to isolate whole-genomic DNA. However, we altered the original enzymatic digestion period (1 h) to recuperate the carapace of copepods for morphological identification to 30 min and to fully digest the tough outside cuticle of nematodes to 12 h. The detailed protocol has also been deposited at https://protocols.io/ under DOI https://dx.doi.org/10.17504/protocols.io.kqdg3re8eg25/v1.

We barcoded the main regions that are regularly used for each parasite group. In all of these groups, the target regions include one or several of the following markers: 18S and 28S nuclear ribosomal DNA, internal transcribed spacers 1 (ITS1) and 2 (ITS2), and the cytochrome *c* oxidase subunit I mitochondrial DNA (*cox*1). To amplify these regions, we used 12.5 µl of 2× MangoMix (Bioline, London, UK) and 0.5 µl of 50 mM MgCl_2_ with the amount of 10 µM primers and ddH2O adapted to each primer combination (and by extension parasite group), to reach a total volume of 25 µl. The selected primers and target taxa are listed in Table 3. Primer concentration and PCR conditions are supplied in Supplementary Table S1. Successful reactions were confirmed by means of gel electrophoresis. PCR products were purified using the GeneJet Gel Extraction Kit (ThermoFisher Scientific, Waltham, MA, USA) or ExoSAP-IT PCR Product Cleanup Reagent (ThermoFisher Scientific, Waltham, MA, USA) according to manufacturer instructions. Sanger sequencing was outsourced to Macrogen Europe (Amsterdam, The Netherlands).

**Table 3.**
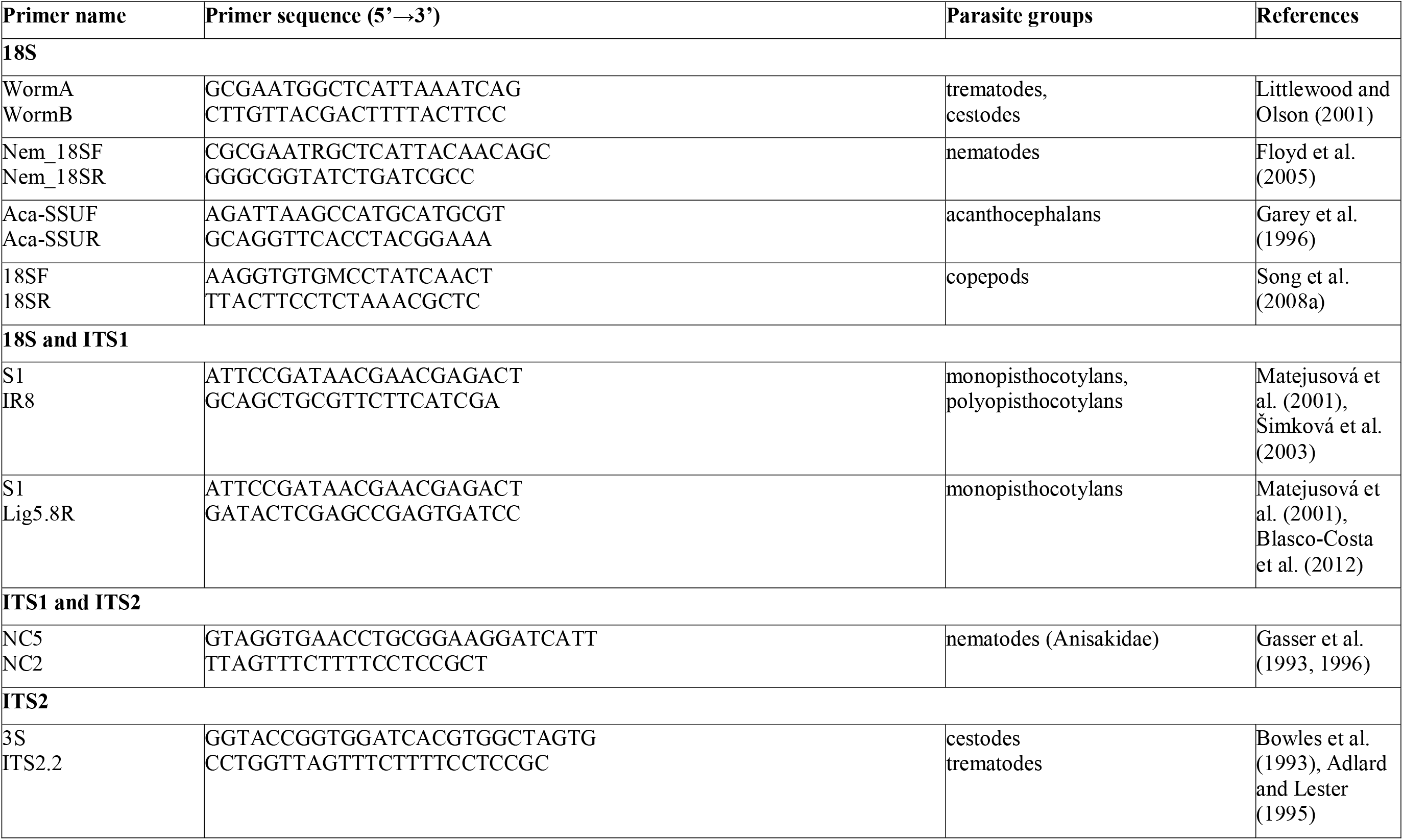
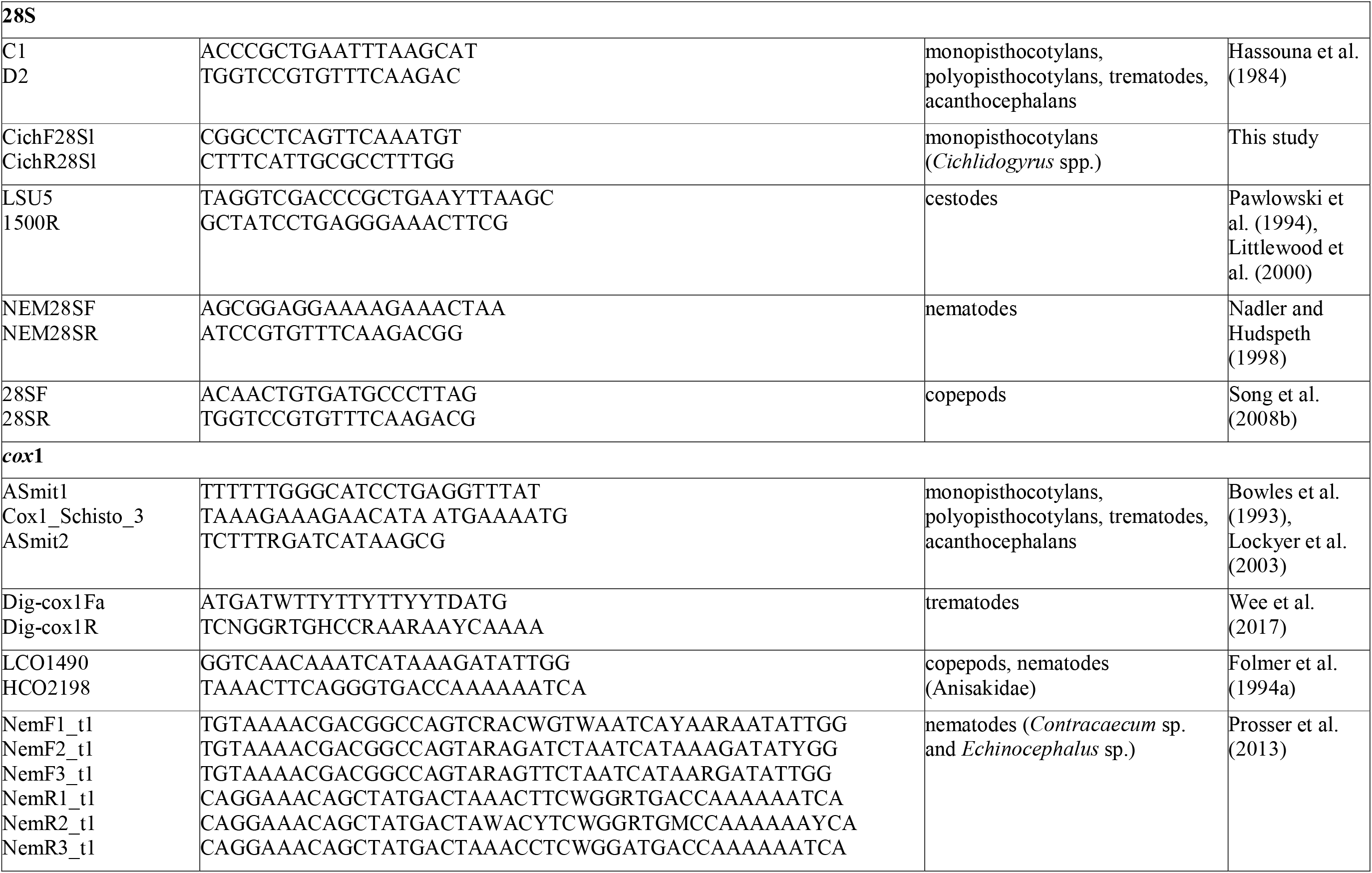
List of primers used in present study for amplification and sequencing of barcoding regions of different parasite groups.

Some of our sequences from monopisthocotylans were similar to reference sequences of *Euryhaliotrema* sp. deposited on GenBank (MG586859–MG586860). These samples were originally collected from *Lutjanus griseus* (L., 1758) (Acanthuriformes: Lutjanidae) at Campeche Bank, Playa Seyba, Mexico (coordinates: 19.710, −90.736) in 2017 by Mendoza-Franco et al. (2018). However, these sequences are presently considered ‘unpublished’ records without available hologenophores. Because these records currently lack associated hologenophores and formal publication, we formally establish and publish them in this study.

### Comparative analysis of parasite communities

We tested for differences between the parasite communities in the MPA and OAA, both at the level of species diversity associated with the individual hosts (infracommunities) and the host populations (component communities). We emphasise that samples have only been collected in a single month and the samples per host and per site are limited. Therefore, the aim of these analyses is not a representative comparison of the impact of this specific MPA or MPAs in general on parasite communities. Rather, we aimed to obtain a first insight into the suitability of the Saloum Delta as a study system to detect differences in fish-parasite community composition.

We performed four separate analyses to compare the biodiversity of the MPA with the OAA. All analyses were performed for three separate datasets: **(A)** gut parasites, **(B)** gill parasites, and **(C)** combined gut and gill parasites, restricted to host individuals screened for both organs.

*(i) A generalised linear model to test the effect of sampling site, host size, and host species on the biodiversity of parasite infracommunities*. Previous studies show that the size of the host individual is often positively associated with abundance (Lo et al. 1998; Braicovich et al. 2016; Henriksen et al. 2023) and the species richness (Luque et al. 2004; Braicovich et al. 2016; Calhoun et al. 2018) of parasite infracommunities. To test the effect of host size, we calculated standard biodiversity indices (Simpson and Shannon indices) for the parasite infracommunity infecting each host individual, rather than the extrapolation-based indices applied in (ii) and (iii), to reflect the biodiversity of the infra-communities. We opted for this approach as the number of individuals of the infracommunities may be considered as fully captured, at least for the targeted organs, because the samples had been screened extensively.

We built models for each dataset and index as follows:

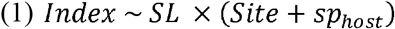

Where *Index* is the respective biodiversity index, *SL* is the standard length of the host, *Site* is the sampling area (MPA or OAA), and *sp_host_* is the identity of the host species. We accounted for potential interaction effects between the *SL* and *Site*, as fisheries pressure may affect host sizes. Furthermore, we modelled an interaction effect between *SL* and *sp_host_*. Models were built using glmmTMB v1.1.13 (McGillycuddy et al. 2025) in R v4.5.0 (R Core Team 2025). For the Shannon index, we applied a zero-inflated Tweedie distribution due to an excess of null values. For the Simpson index, we applied a beta-regression. This distribution is suitable for continuous data bounded strictly within the open interval (0, 1). Because our dataset included exact values of 0 and 1, we transformed the Simpson estimates as suggested by Smithson and Verkuilen (2006):

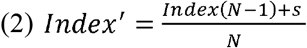

where *Index* is the respective biodiversity index*, N* is the sample size (total number of screened hosts) and *s* is a constant, which was set to 0.5. Coefficients from the Beta regression models were exponentiated (OR = *e^β^*) and reported as odds ratios (OR). Because the Simpson index is inversely related to species richness, an OR 1 represents an increase in the odds of observing higher Simpson values (i.e., lower infracommunity diversity), whereas an OR 1 indicates a shift toward higher diversity.

Models were simplified using a backward selection procedure using ‘drop1’ with a χ² test in R. All effects that failed to significantly improve model fit were removed, starting from the interaction effects. Based on the minimum adequate model, we predicted biodiversity indices through emmeans v1.11.2.8 (Lenth 2025) in R. The resulting figures were plotted through the emmeans and ggplot2 4.0.0 (Wickham 2016).

*(ii) Biodiversity indices of component communities based on parasite individuals and abundance data (counts).* In natural observation data, a number of species are likely to remain unobserved. The number of potentially unseen species decreases with repeated observations. This trend can be approximated using extrapolation-based biodiversity indices that provide estimations of the total number of species present. We estimated Chao, Gini-Simpson, and Shannon entropy indices for abundance data (number of observed parasite individuals) as implemented in iNEXT v3.0.2 (Chao et al. 2014) in R. We selected this Hill-number framework over traditional estimators (e.g., Jackknife) because it provides a unified scale for richness and diversity metrics while allowing comparisons based on sample coverage (sampling completeness). These indices are widely used for estimating biodiversity and statistically significant differences can be assessed through the 95% confidence intervals of each estimate (Chao et al. 2014). Note that for individual host GLMs (i), the response variable is the standard Simpson index, where higher values denote single-species dominance (lower diversity). In contrast, extrapolation analyses in iNEXT report Simpson diversity as Hill number order 2 (Gini-Simpson index), where higher values denote higher diversity.

We performed additional analyses to verify trends observed in (ii). First, biodiversity indices can also be extrapolated based on host individual and incidence (presence-absence), which more explicitly accounts for sampling effort. This analysis was performed as in (ii), but species matrices were transformed to presence-absence data. Second, interacting species can also be modelled as ecological networks, with species represented as discrete units (nodes) and interactions represented as links (edges) (Runghen et al. 2021). Ecological network diversity can be then estimated by extrapolating individual host-parasite interactions.

Extrapolation can be used to infer the number of unobserved interactions in the networks by modelling individual interactions as observed units. We compared the observed ecological networks in the MPA with the OAA by estimating Chao, Gini-Simpson, and Shannon entropy indices through iNEXT.link v1.0.0 (Chiu et al. 2023), an extension of the iNEXT package for ecological networks. For the results of these additional biodiversity analyses, see Supplementary File S2.

## Results

### Parasite survey: overview

We screened 126 gills and 81 guts from 14 host species, of which 104 gills and 59 guts were from target host species (*C. guineensis*, *C. hippos*, *E. fimbriata*, *G. nigri*, *H. fasciatus*, *I. africana, M. sebae*, and *S. melanotheron*) and the other samples were from the remaining five host species (*E. lacerta*, *E. aeneus*, *P. quadrifilis*, *S.afra*, and *C*. *amnicola)*. We collected 5,020 macroparasite individuals belonging to 27 species; 23 of which were found to infect eight of the nine target host species (no parasites were found on 22 specimens of *P. barbarus*). A large majority (96.8%) of parasite specimens belonged to Monopisthocotyla, followed by Acanthocephala (1.1%), Nematoda (0.9%), Polyopisthocotyla (0.7%), Trematoda (0.4%), Copepoda (0.1%), and Cestoda (<0.01%). Eighteen parasite species could be assigned to a described species. Six species could be assigned generic names, three of which we recognised as undescribed species (Monopisthocotyla: *Euryhaliotrema* spp.). For an overview of host-parasite combinations see Table 4. For details on the species- and genus-level identifications see the following section. The collection data of individual hosts and parasites can be found in Supplementary Table S3. Furthermore, we provide 208 DNA barcodes from 26 species identified here, all of which represent the first molecular data for these species in West Africa (Table 5). For eight (31%) of these species, these are the first molecular data worldwide; for 14 (54%), we provide the first sequence(s) for at least one DNA region.

**Table 4.**
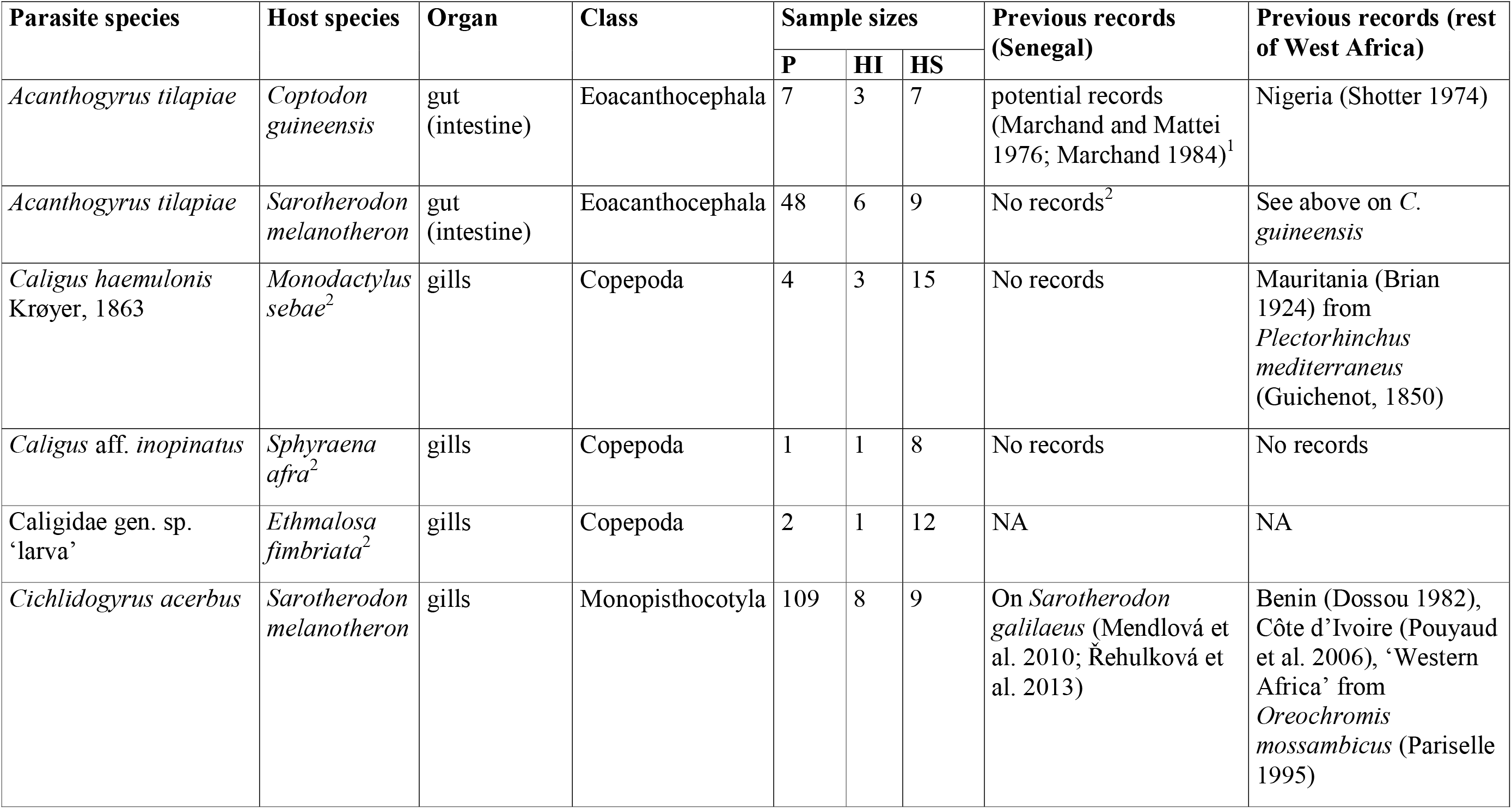
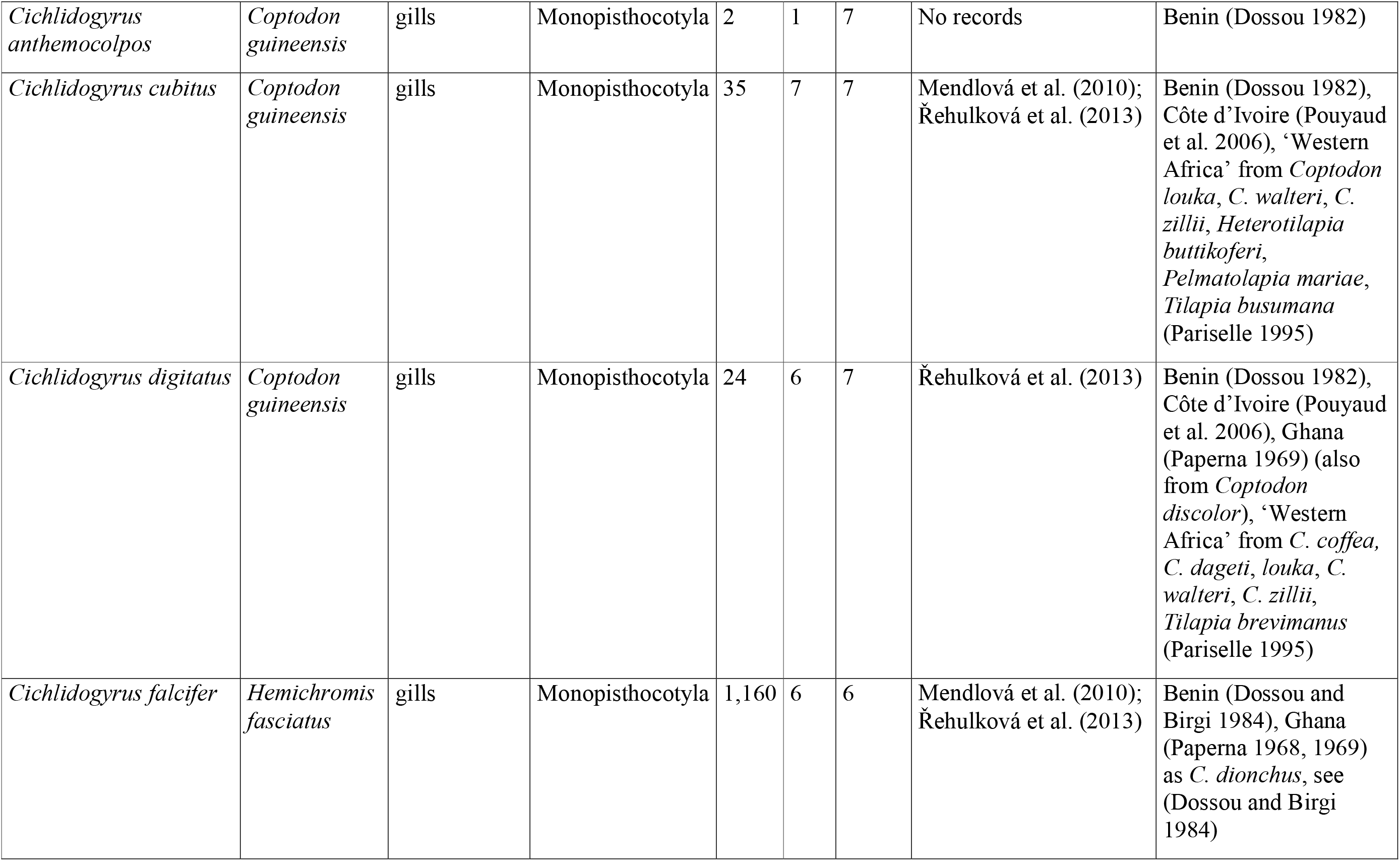
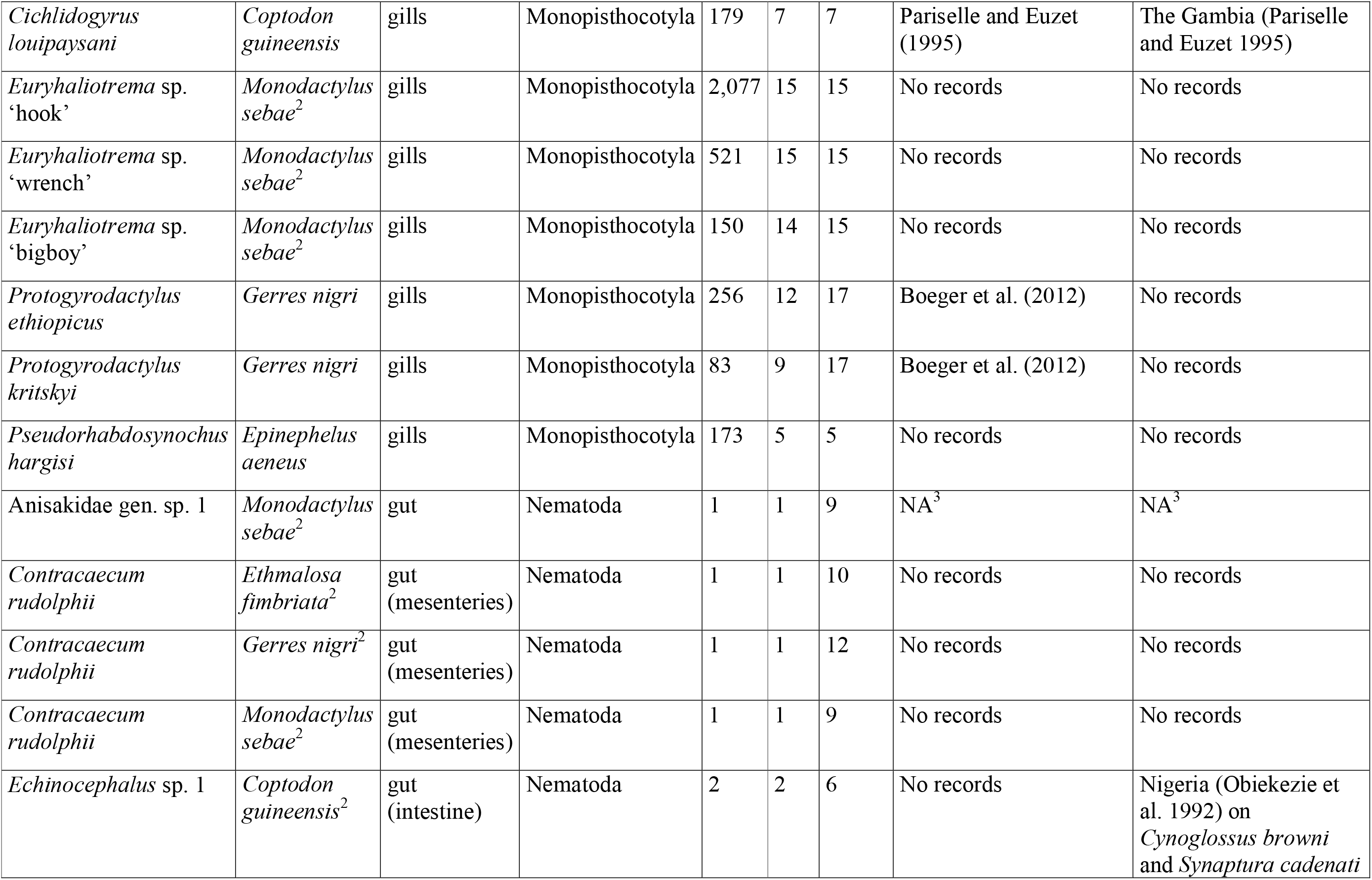
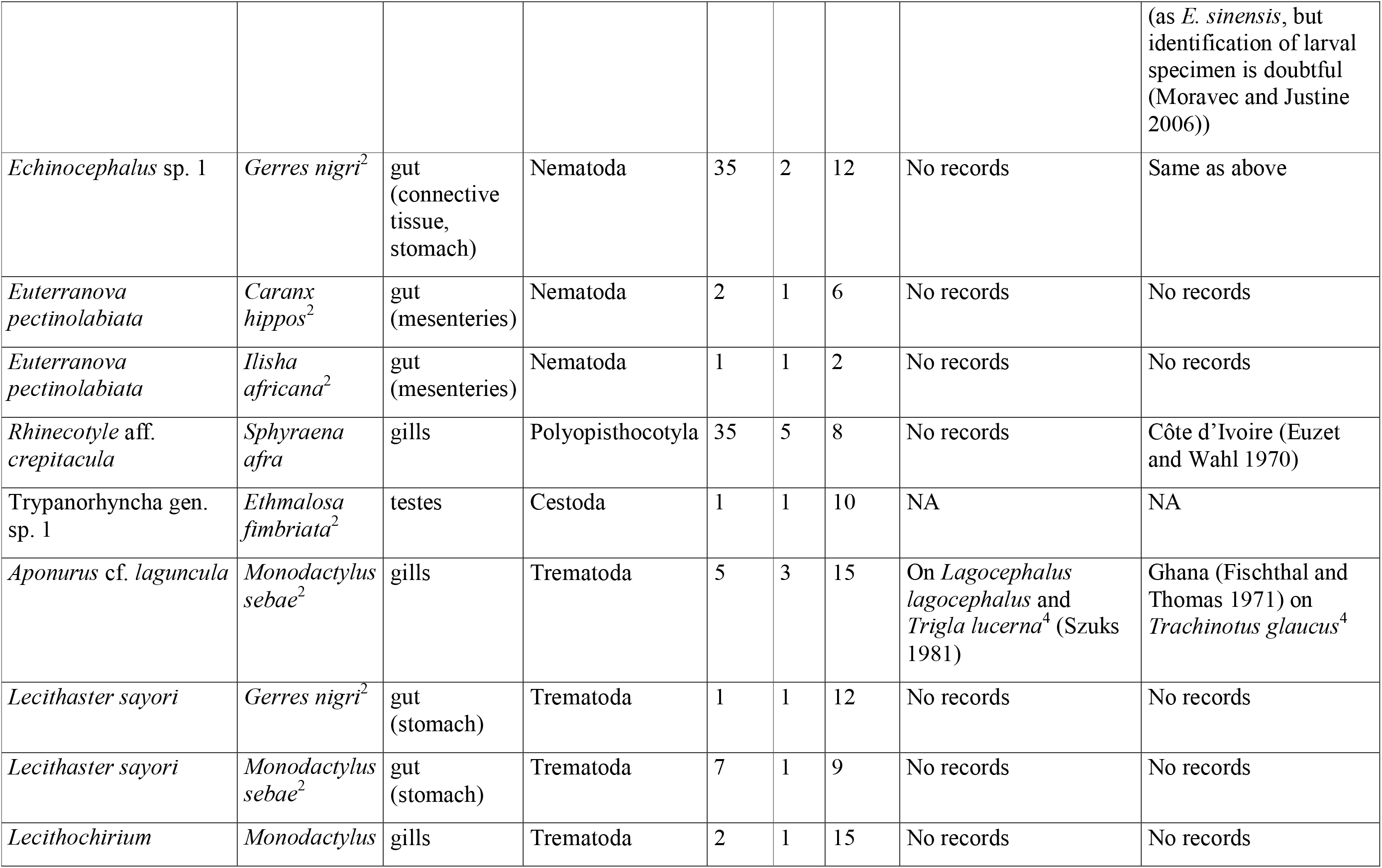
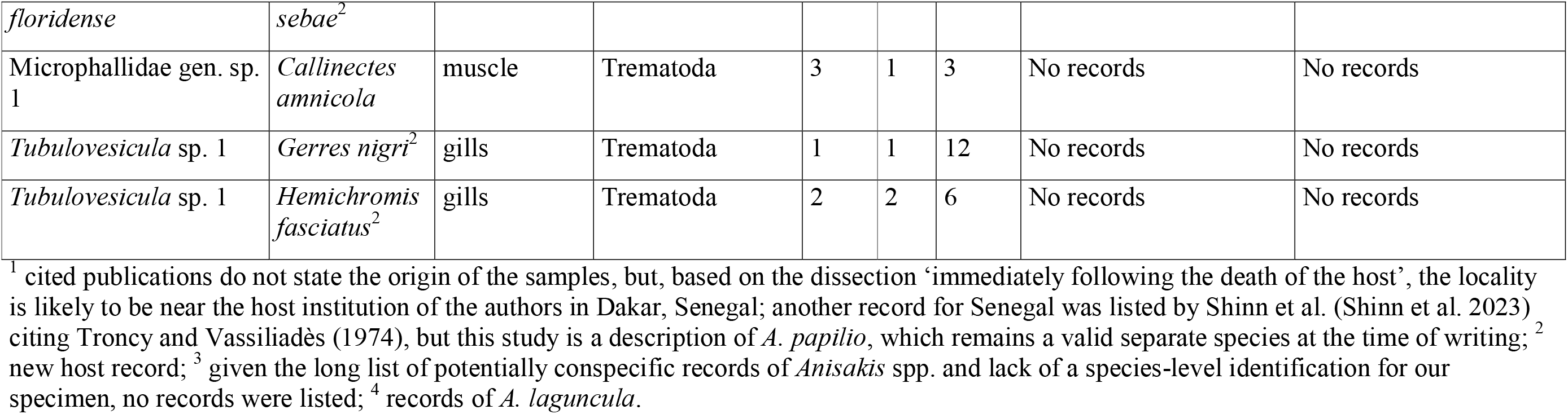
Host-parasite combination found in the Saloum Delta in the Bamboung MPA and the Bolong de Sangako OAA, sample sizes of parasites (P), hosts infected (HI), and hosts screened (HS) and information on previous records from Senegal and other West African countries (as defined by UNSTATS). Previous records from a different host species are listed alongside the host name (NA = previous records not applicable due to lack of species-level identification).

| **Parasite species** | **Host species** | **Organ** | **Class** | **Sample sizes** |  |  | **Previous records (Senegal)** | **Previous records (rest of West Africa)** |
|---|---|---|---|---|---|---|---|---|
|  |  |  |  | **P** | **HI** | **HS** |  |  |
| *Acanthogyrus tilapiae* | *Coptodon guineensis* | gut (intestine) | Eoacanthocephala | 7 | 3 | 7 | potential records (Marchand and Mattei 1976; Marchand 1984)<sup>1</sup> | Nigeria (Shotter 1974) |
| *Acanthogyrus tilapiae* | *Sarotherodon melanotheron* | gut (intestine) | Eoacanthocephala | 48 | 6 | 9 | No records<sup>2</sup> | See above on *C. guineensis* |
| *Caligus haemulonis* Krøyer, 1863 | *Monodactylus sebae*<sup>2</sup> | gills | Copepoda | 4 | 3 | 15 | No records | Mauritania (Brian 1924) from *Plectorhinchus mediterraneus* (Guichenot, 1850) |
| *Caligus* aff. *inopinatus* | *Sphyraena afra*<sup>2</sup> | gills | Copepoda | 1 | 1 | 8 | No records | No records |
| Caligidae gen. sp. ‘larva’ | *Ethmalosa fimbriata*<sup>2</sup> | gills | Copepoda | 2 | 1 | 12 | NA | NA |
| *Cichlidogyrus acerbus* | *Sarotherodon melanotheron* | gills | Monopisthocotyla | 109 | 8 | 9 | On *Sarotherodon galilaeus* (Mendlová et al. 2010; Řehulková et al. 2013) | Benin (Dossou 1982), Côte d’Ivoire (Pouyaud et al. 2006), ‘Western Africa’ from *Oreochromis mossambicus* (Pariselle 1995) |
| *Cichlidogyrus anthemocolpos* | *Coptodon guineensis* | gills | Monopisthocotyla | 2 | 1 | 7 | No records | Benin (Dossou 1982) |
| *Cichlidogyrus cubitus* | *Coptodon guineensis* | gills | Monopisthocotyla | 35 | 7 | 7 | Mendlová et al. (2010); Řehulková et al. (2013) | Benin (Dossou 1982), Côte d’Ivoire (Pouyaud et al. 2006), ‘Western Africa’ from *Coptodon louka*, *C. walteri*, *C. zillii*, *Heterotilapia buttikoferi*, *Pelmatolapia mariae*, *Tilapia busumana* (Pariselle 1995) |
| *Cichlidogyrus digitatus* | *Coptodon guineensis* | gills | Monopisthocotyla | 24 | 6 | 7 | Řehulková et al. (2013) | Benin (Dossou 1982), Côte d’Ivoire (Pouyaud et al. 2006), Ghana (Paperna 1969) (also from *Coptodon discolor*), ‘Western Africa’ from *C. coffea,* *C. dageti*, *louka*, *C. walteri*, *C. zillii*, *Tilapia brevimanus* (Pariselle 1995) |
| *Cichlidogyrus falcifer* | *Hemichromis fasciatus* | gills | Monopisthocotyla | 1,160 | 6 | 6 | Mendlová et al. (2010); Řehulková et al. (2013) | Benin (Dossou and Birgi 1984), Ghana (Paperna 1968, 1969) as *C. dionchus*, see (Dossou and Birgi 1984) |
| *Cichlidogyrus louipaysani* | *Coptodon guineensis* | gills | Monopisthocotyla | 179 | 7 | 7 | Pariselle and Euzet (1995) | The Gambia (Pariselle and Euzet 1995) |
| *Euryhaliotrema* sp. ‘hook’ | *Monodactylus sebae*<sup>2</sup> | gills | Monopisthocotyla | 2,077 | 15 | 15 | No records | No records |
| *Euryhaliotrema* sp. ‘wrench’ | *Monodactylus sebae*<sup>2</sup> | gills | Monopisthocotyla | 521 | 15 | 15 | No records | No records |
| *Euryhaliotrema* sp. ‘bigboy’ | *Monodactylus sebae*<sup>2</sup> | gills | Monopisthocotyla | 150 | 14 | 15 | No records | No records |
| *Protogyrodactylus ethiopicus* | *Gerres nigri* | gills | Monopisthocotyla | 256 | 12 | 17 | Boeger et al. (2012) | No records |
| *Protogyrodactylus kritskyi* | *Gerres nigri* | gills | Monopisthocotyla | 83 | 9 | 17 | Boeger et al. (2012) | No records |
| *Pseudorhabdosynochus hargisi* | *Epinephelus aeneus* | gills | Monopisthocotyla | 173 | 5 | 5 | No records | No records |
| Anisakidae gen. sp. 1 | *Monodactylus sebae*<sup>2</sup> | gut | Nematoda | 1 | 1 | 9 | NA<sup>3</sup> | NA<sup>3</sup> |
| *Contracaecum rudolphii* | *Ethmalosa fimbriata*<sup>2</sup> | gut (mesenteries) | Nematoda | 1 | 1 | 10 | No records | No records |
| *Contracaecum rudolphii* | *Gerres nigri*<sup>2</sup> | gut (mesenteries) | Nematoda | 1 | 1 | 12 | No records | No records |
| *Contracaecum rudolphii* | *Monodactylus sebae*<sup>2</sup> | gut (mesenteries) | Nematoda | 1 | 1 | 9 | No records | No records |
| *Echinocephalus* sp. 1 | *Coptodon guineensis*<sup>2</sup> | gut (intestine) | Nematoda | 2 | 2 | 6 | No records | Nigeria (Obiekezie et al. 1992) on *Cynoglossus browni* and *Synaptura cadenati* (as *E. sinensis*, but identification of larval specimen is doubtful (Moravec and Justine 2006)) |
| *Echinocephalus* sp. 1 | *Gerres nigri*<sup>2</sup> | gut (connective tissue, stomach) | Nematoda | 35 | 2 | 12 | No records | Same as above |
| *Euterranova pectinolabiata* | *Caranx hippos*<sup>2</sup> | gut (mesenteries) | Nematoda | 2 | 1 | 6 | No records | No records |
| *Euterranova pectinolabiata* | *Ilisha africana*<sup>2</sup> | gut (mesenteries) | Nematoda | 1 | 1 | 2 | No records | No records |
| *Rhinecotyle* aff. *crepitacula* | *Sphyraena afra* | gills | Polyopisthocotyla | 35 | 5 | 8 | No records | Côte d’Ivoire (Euzet and Wahl 1970) |
| Trypanorhyncha gen. sp. 1 | *Ethmalosa fimbriata*<sup>2</sup> | testes | Cestoda | 1 | 1 | 10 | NA | NA |
| *Aponurus* cf. *laguncula* | *Monodactylus sebae*<sup>2</sup> | gills | Trematoda | 5 | 3 | 15 | On *Lagocephalus lagocephalus* and *Trigla lucerna*<sup>4</sup> (Szuks 1981) | Ghana (Fischthal and Thomas 1971) on *Trachinotus glaucus*<sup>4</sup> |
| *Lecithaster sayori* | *Gerres nigri*<sup>2</sup> | gut (stomach) | Trematoda | 1 | 1 | 12 | No records | No records |
| *Lecithaster sayori* | *Monodactylus sebae*<sup>2</sup> | gut (stomach) | Trematoda | 7 | 1 | 9 | No records | No records |
| *Lecithochirium floridense* | *Monodactylus sebae*<sup>2</sup> | gills | Trematoda | 2 | 1 | 15 | No records | No records |
| Microphallidae gen. sp. 1 | *Callinectes amnicola* | muscle | Trematoda | 3 | 1 | 3 | No records | No records |
| *Tubulovesicula* sp. 1 | *Gerres nigri*<sup>2</sup> | gills | Trematoda | 1 | 1 | 12 | No records | No records |
| *Tubulovesicula* sp. 1 | *Hemichromis fasciatus*<sup>2</sup> | gills | Trematoda | 2 | 2 | 6 | No records | No records |

<sup>1</sup> cited publications do not state the origin of the samples, but, based on the dissection ‘immediately following the death of the host’, the locality is likely to be near the host institution of the authors in Dakar, Senegal; another record for Senegal was listed by Shinn et al. (Shinn et al. 2023) citing Troncy and Vassiliadès (1974), but this study is a description of *A. papilio*, which remains a valid separate species at the time of writing; <sup>2</sup> new host record; <sup>3</sup> given the long list of potentially conspecific records of *Anisakis* spp. and lack of a species-level identification for our specimen, no records were listed; <sup>4</sup> records of *A. laguncula*.

**Table 5.**
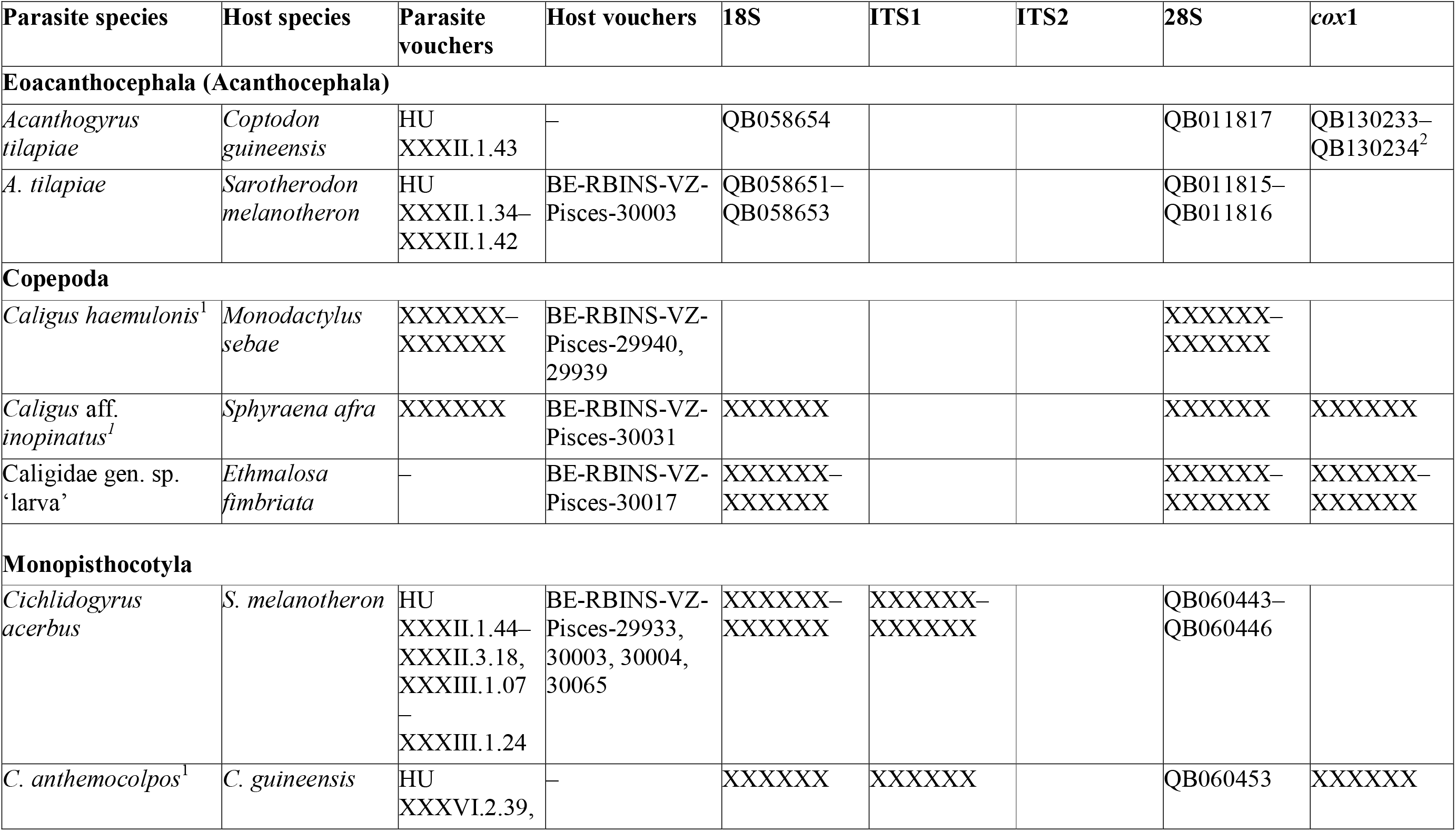
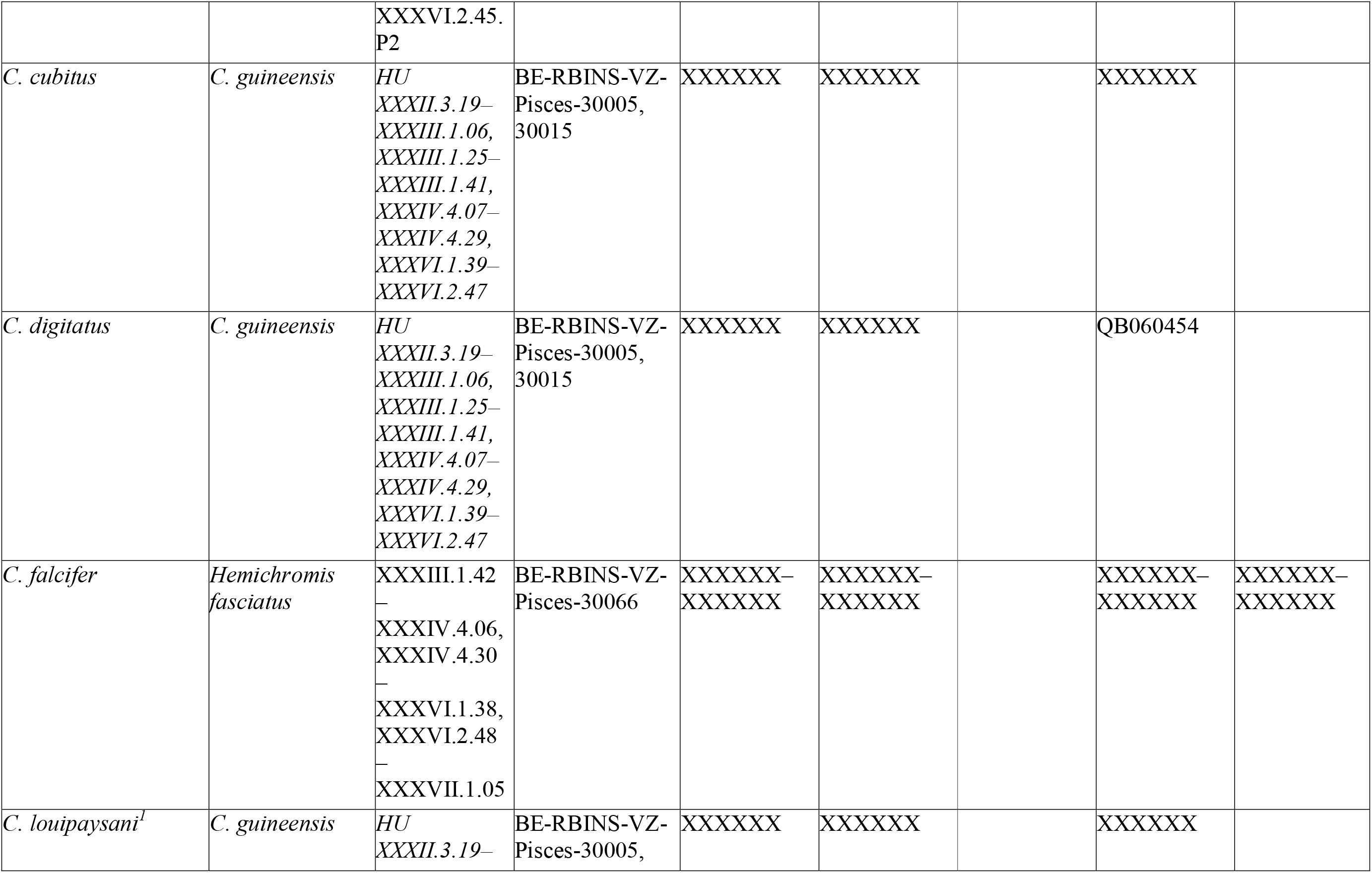
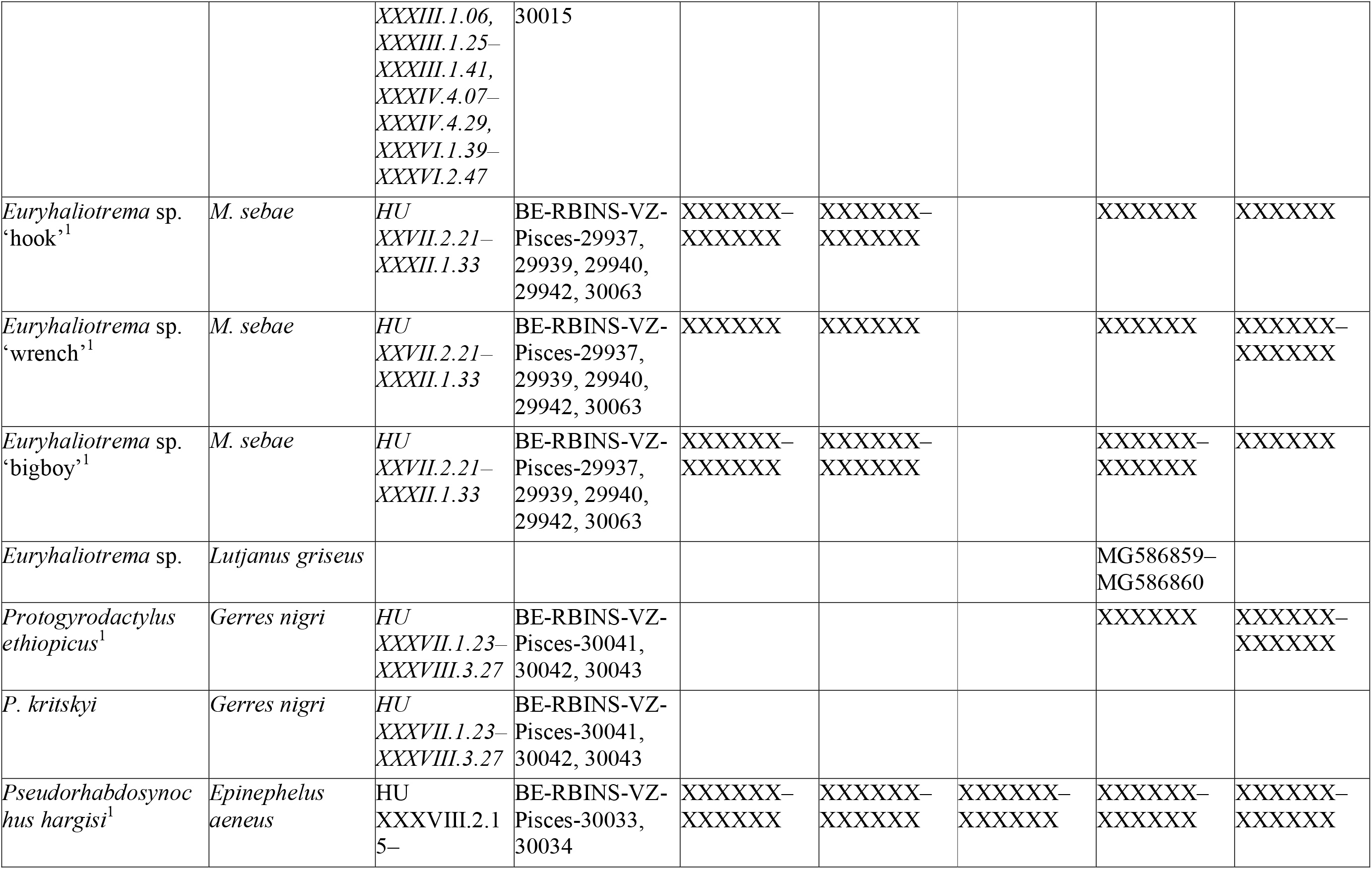
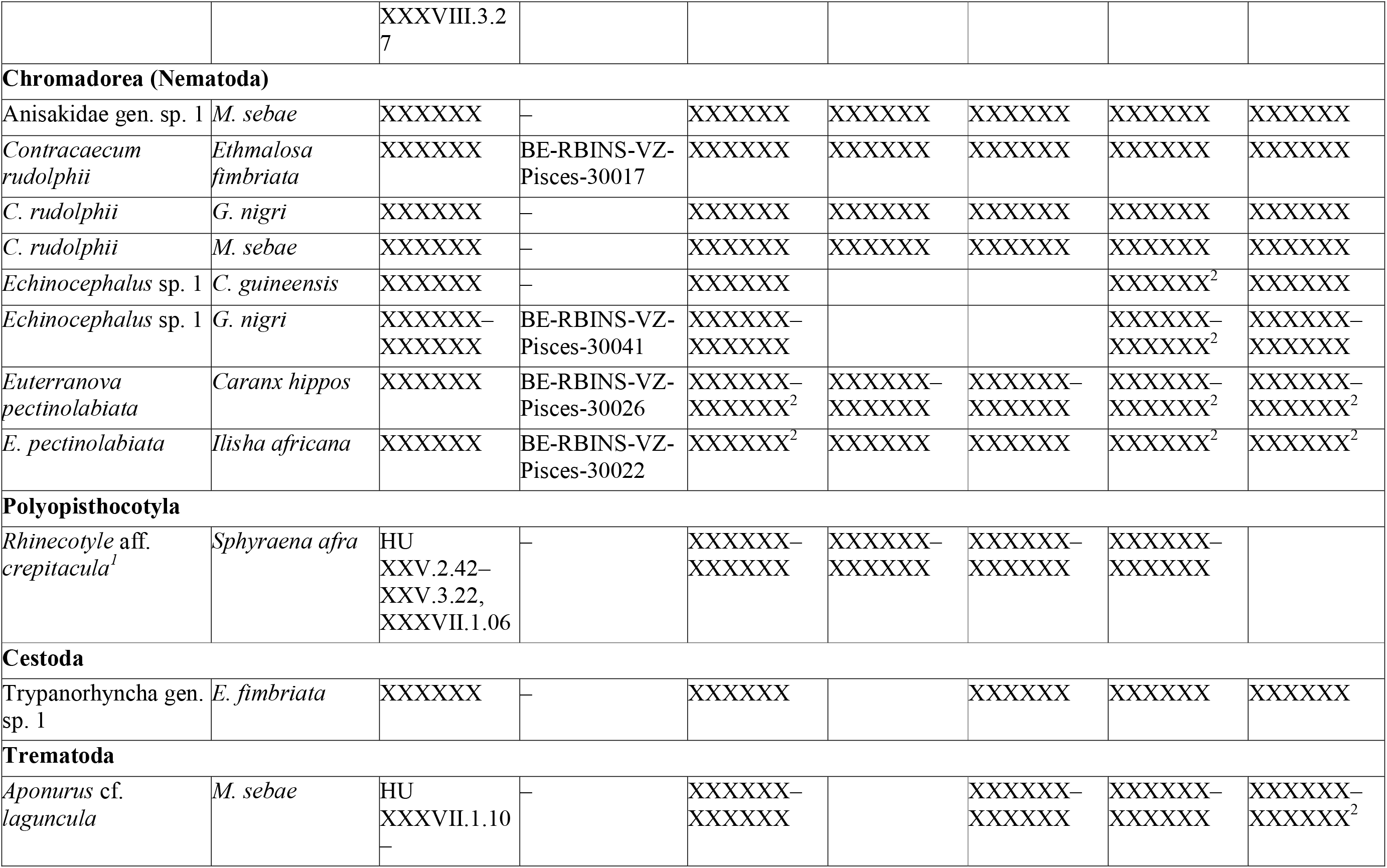
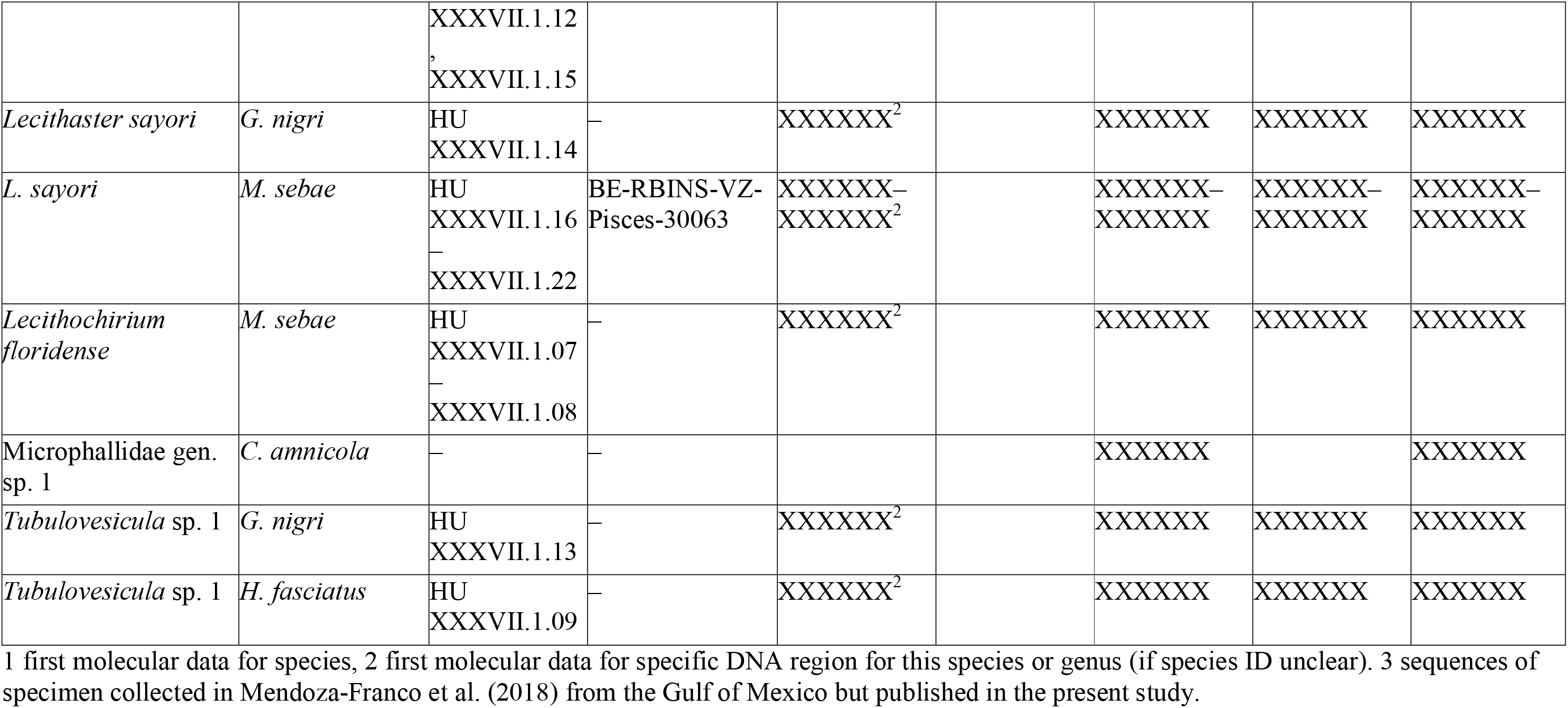
List of parasite and host species examined in this study, including museum voucher numbers and GenBank accession numbers for the sequenced molecular markers (18S, ITS1, ITS2, 28S, and *cox*1). Parasite vouchers in italics are grouped by host species; for complete list of hosts and parasite individuals see Supplementary Table S3.

**Table 6.** Generalised linear model (GLM) selection for parasite infracommunity diversity (Shannon–Weaver and Simpson indices) across infection sites (gills vs. combined gills and gut). Shown are the retained predictors in the minimum adequate models following backward stepwise selection. Reported ΔAIC values reflect the loss of model fit associated with removing each predictor from the final model.

| **Dependent variable** | **Infection site** | **Effects in minimum adequate models** | **ΔAIC** |
|---|---|---|---|
| Shannon-Weaver index | gills | Host species | 92 |
|  | gills + gut | Host species | 16.6 |
| Simpson index | gills | Host species | 1.54 |
|  | gills + gut | Site<br>SL<br>Host species | 7.8<br>7.4<br>1.7 |

### Monopisthocotyla Brabec, Salomaki, Scholz & Kuchta, 2023

#### Dactylogyridea Bychowsky, 1937

#### Dactylogyridae Bychowsky, 1933

Most monopisthocotylan specimens (other than for the parasite of *E. aeneus*) were identified as members of Dactylogyridae, based on the presence of a two-lobed opisthohaptor with well-developed anchors and supporting bars, as well as the absence of adhesive plaque-like or tubular structures (Yamaguti 1963).

#### *Cichlidogyrus* Paperna 1960

*Cichlidogyrus* spp.

The monopisthocotylan flatworms infecting the gills of cichlid fishes (*C. guineensis*, *H. fasciatus*, *S. melanotheron*) were assigned to *Cichlidogyrus*, based on their two pairs of anchors, and two transverse bars, with the dorsal bar possessing two auricles and the ventral bar being V-shaped. Species of *Cichlidogyrus* have been reported multiple times from these same host species (Paperna 1969, 1979; Dossou 1982; Dossou and Birgi 1984; Pariselle 1995;

Pariselle and Euzet 1995, 2004; Pariselle et al. 2013).

The specimens found on *S. melanotheron* all matched the description of *C. acerbus* Dossou, 1982 (Fig. 2B–C), with its characteristic morphology of the male copulatory organ (MCO), including a tubular, slightly arched copulatory tube, and a complex accessory piece with many irregularly shaped protuberances (Dossou 1982) – the only species with this type of MCO to be reported from *S. melanotheron*. However, our specimens fell within two groups based on the morphology of the accessory piece, with *C. acerbus* A (Fig. 2B; 3A) presenting a large plate-like structure with a distal hook and *C. acerbus* B (Fig. 2C; 3B–C) possessing a similar structure but with much smaller, incurved distal hook and serrations on its surface. These plate-like structures were not explicitly mentioned or illustrated in the description of *C. acerbus* (Dossou 1982). However, the three-dimensional structure of the accessory piece is inherently complex, which may be poorly reflected by two-dimensional drawings. We also noted that the serrations and distal hook of the plate-like structure of *C. acerbus* A are reminiscent of the MCO morphology of *C. ophioglossus* Moons, Kmentová, Pariselle, Vanhove & Cruz-Laufer, 2023, a parasite of *Chromidotilapia kingsleyae* Boulenger, 1898 and *C. elongata* Lamboj, 1999 from Central Africa. Therefore, we suggest that the relationship between *C. acerbus* and these other taxa should be explored in future studies.

**Figure 2.**
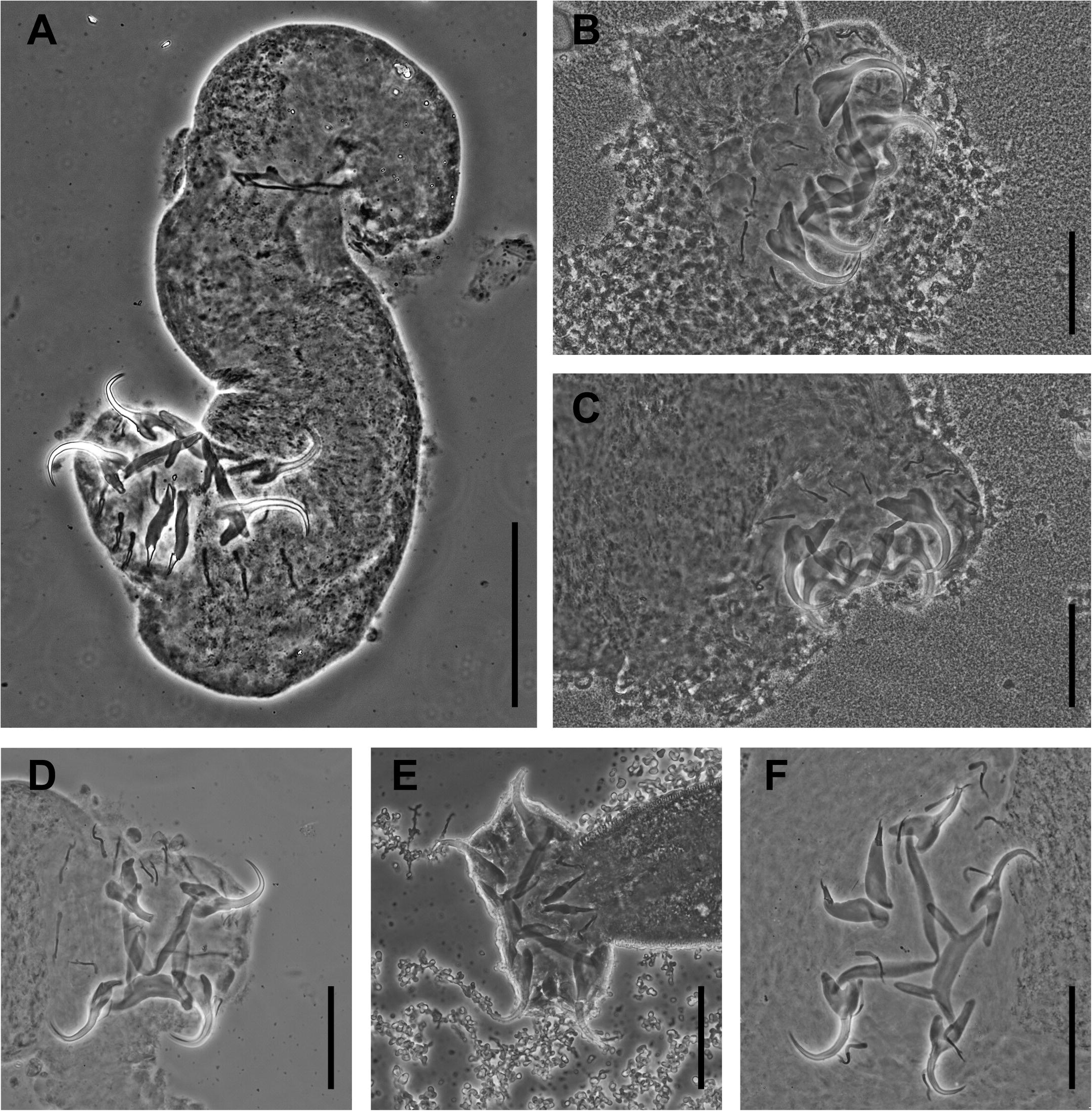
Microphotographs of *Cichlidogyrus* spp. collected off the gills of cichlid fishes using phase contrast, part 1. **A** Overview of whole body (*C. falcifer*). Haptoral sclerites of **B** *C. acerbus* A ; **C** *C. acerbus* B; **D** *C. cubitus*; **E** *C. louipaysani*; and **F** *C. digitatus*. The distal half of the body including the haptoral sclerites of *C. anthemocolpos* were used for DNA extraction. Scale bars: 100 µm (**A**); 50 µm (**B–F**).

The specimens found on *C. guineensis* belonged to four different species, including *Cichlidogyrus cubitus* Dossou, 1982 (Fig. 2D; 3D), *C. louipaysani* Pariselle & Euzet, 1995 (Fig. 2E; 3E), *C. digitatus* Dossou, 1982 (Fig. 2F; 3F), and *C. anthemocolpos* Dossou, 1982 (Fig. 3G, H). Specimens of *C. cubitus* were recognised by the ribbon-like accessory piece with a forked distal end (Dossou 1982), *C. louipaysani* by the accessory piece like *C. cubitus*, but with serration at the forked distal end (Pariselle and Euzet 1995), *C. digitatus* by two hooks at the distal end of the accessory piece (Dossou 1982), and *C. anthemocolpos* by the haptor with elongated marginal hooks 3–7, MCO with a complex, S-shaped accessory piece, and sclerotised vagina with reniform plate (Dossou 1982). The specimens found on *H. fasciatus* were identified as *C. falcifer* (Fig. 2A; Fig. 3I), based on the MCO that presented an arched, tubular copulatory tube and an accessory piece in the shape of a filament with a thickened nodular portion in its midsection (Dossou and Birgi 1984).

**Figure 3.**
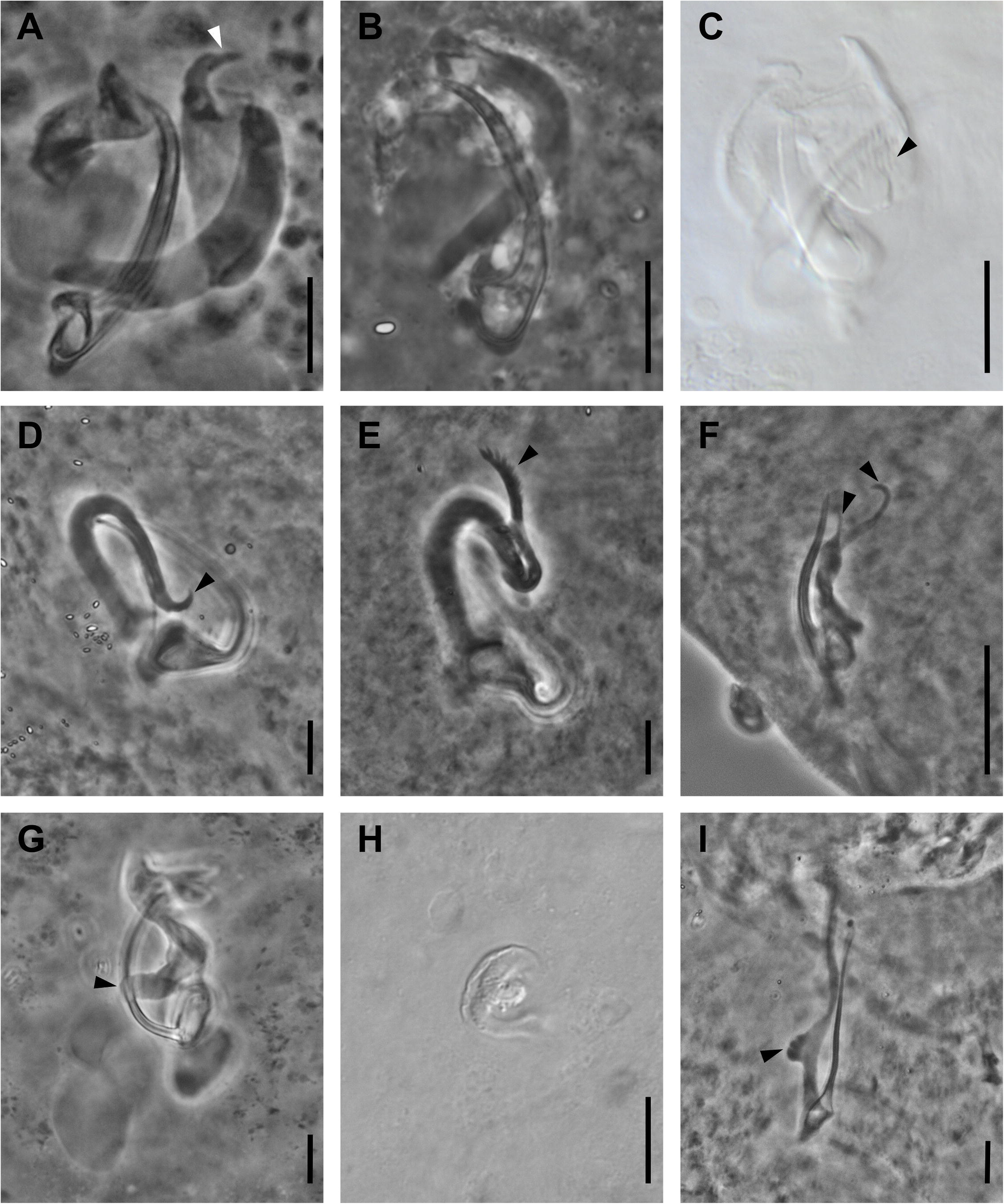
Microphotographs of *Cichlidogyrus* spp. collected off the gills of cichlid fishes using phase contrast or differential interference contrast, part 2. **A** Male copulatory organ (MCO) of *C. acerbus* A possessing a large distal hook in the plate-like portion of the accessory piece; **B**, **C** MCO of *C. acerbus* B with serrations in the plate-like portion of the accessory piece; **D** MCO of *C. cubitus*; **E** MCO of *C. louipaysani* with feather-like distal end of the accessory piece (which is absent in *C. cubitus*); **F** MCO of *C. digitatus* with distal hooks; **G** MCO of *C. anthemocolpos* with proximal flap of accessory piece; **H** reniform vaginal sclerites of *C. anthemocolpos*; **I** MCO of *C. falcifer* with node-like thickening in mid-portion of accessory piece. Scale bars: 10 µm (**A–I**).

Mirroring the observed morphological differences, sequences of *C. acerbus* clustered in two distinct groups, *C. acerbus* A and *C. acerbus* B, with substantial differences in their DNA sequence data (28S: 97.2%, 18S: 99.0%, ITS1: 86.3%). All sequences were most similar to those of specimens of *C. acerbus* collected in Côte d’Ivoire from the same host species (Pouyaud et al. 2006) (group A – ITS1: 93.3%, 18S: 98.6%; group B – ITS1: 99.0%, 18S: 98.9%), but our specimens differed substantially from the specimens of *C. acerbus* from eastern Senegal from *Sarotherodon galilaeus* (L. 1758) (Mendlová et al. 2010, 2012) (group A – ITS1: 90.0%, 18S: 99.6%, 28S: 93.8%; group B – ITS1: 94.0%, 18S: 99.4%, 28S: 94.8%). Given these low levels of sequence identity between our own specimens and the specimens from other studies, we suggest that these specimens may constitute at least two distinct species: *Cichlidogyrus acerbus* A collected in the present study and *C. acerbus* B collected in the present study and in Côte d’Ivoire. The specimens from *S. galilaeus* in eastern Senegal may constitute a third lineage, but the authors provided no morphological data or deposited voucher specimens (Mendlová et al. 2010, 2012). Therefore, their identification could not be verified. We suggest that these differences in the morphological and molecular information call for a revision of *C. acerbus* as well as further studies into its status as a single species. Pending a more detailed taxonomic study, we treated *C. acerbus* A and *C. acerbus* B as the same species for the species richness analyses (see below).

We provide the first molecular data for *C. louipaysani*. Sequences of *C. cubitus* and *C. louipaysani* showed high sequence identity (28S: 100%, 18S: 99.4%–99.8%, ITS1: 99.8– 100%), which reflects the suggested close relationship between these two species (Pariselle and Euzet 1995). Our sequences of *C. cubitus* were also similar to sequences of specimens previously assigned to *C. cubitus* from *C. guineensis* from eastern Senegal (28S: 100%, 18S: 100%, ITS1: 99.4%–100%) (Mendlová et al. 2010, 2012), Côte d’Ivoire (18S: 98.9%–99.3%, ITS1: 98.0%) (Pouyaud et al. 2006), and Morocco (28S: 99.5%) (Cruz-Laufer et al. 2022), and from *Pelmatolapia mariae* (Boulenger, 1899) in Côte d’Ivoire (28S: 100%) (Cruz-Laufer et al. 2022). Sequences of *C. louipaysani* were also similar to these reference sequences of *C. cubitus* (Senegal – 18S: 100%, ITS1: 99.6%; Côte d’Ivoire/*C. guineensis* – 18S: 98.5%, ITS1: 98.0%; Morocco – 28S: 99.5%, Côte d’Ivoire/*P. mariae* – 99.5%). While the nuclear sequence data identical or very similar among members of the two species, we could unambiguously discriminate the two species based on the morphological characters in the accessory piece of the MCO (see remarks above). We suggest that the close relationship of these two species may necessitate the use of more variable barcoding markers, such as *cox*1. However, this marker was not amplified successfully for these representatives of *Cichlidogyrus* in the present study.

Sequences of a specimen of *C. digitatus* showed high similarity with reference sequences of a specimen from the same host collected in eastern Senegal (28S: 100%, 18S: 99.6%, ITS1: 99.8%). The molecular data, therefore, supported our identification. No *cox*1 sequences could be obtained.

We provide the first molecular data for *C. anthemocolpos*. Sequences of a single specimen were most similar to those of *C. tiberianus* Dossou, 1982 from *Coptodon rendalli* (Boulenger, 1897) in the Lower Congo Basin (28S: 99.6%, 18S: 100%, ITS1: 98.7%) and Madagascar (28S: 99.5%, 18S: 99.8%, ITS1: 99.6%), and from *C. guineensis* in eastern Senegal (28S: 99.6%, 18S: 100%, ITS1: 100%). No reference sequences for the *cox*1 region of *C. tiberianus* were available. The high sequence similarity in the nuclear regions suggests that *C. anthemocolpos* and *C. tiberianus* are close relatives. Morphologically, these species belong to the same clade of *Cichlidogyrus*, sharing their marginal hooks 3–7 with extended shafts and MCO with a tubular, arched copulatory tubes and complex S-shaped accessory piece (Cruz-Laufer et al. 2022). *Cichlidogyrus anthemocolpos* differs from *C. tiberianus* with regard to its kidney-shaped sclerotised vagina (Dossou 1982), unlike the curled tubular structure observed in *C. tiberianus* (Paperna 1960; Ergens 1981; Dossou 1982). *Cichlidogyrus anthemocolpos* has been described from *Coptodon zillii* (Dossou 1982), with later reports finding the parasite on *C. camerunesis* (Lönnberg, 1903) and *C. guineensis* in Cameroon (Pariselle et al. 2013). *Cichlidogyrus tiberianus* has been described from *C. zillii* in Israel (Paperna 1960), with later reports from Egypt and Benin on the same host species (Ergens 1981; Dossou 1982); the parasite has now been reported from a total of 16 host species (Cruz Laufer et al. 2022).

Given the high similarity of our sequences of *Cichlidogyrus anthemocolpos* with those assigned to *C. tiberianus*, we suggest that an in-depth systematic revision of these two species may be required. Furthermore, in the absence of morphological reference data or vouchers (Mendlová et al. 2010, 2012), we suggest that the specimen assigned to *C. tiberianus* from eastern Senegal should be considered a representative of *C. anthemocolpos*, given the geographical proximity, identical host species, and identical ITS1 sequences.

Sequences of *C. falcifer* matched closely with those of specimens from the same species collected in Côte d’Ivoire (28S: 99.4%–99.5%) (Cruz-Laufer et al. 2022), eastern Senegal (28S: 97.2%–97.5%, 18S: 99.6%) (Mendlová et al. 2010, 2012), and the Lower Congo Basin in the Democratic Republic of the Congo (28S: 95.3%–95.4%, 18S: 100%) (Jorissen et al. 2022). However, ITS1 sequences diverged more substantially (86.0%–89.6%) from the Senegalese and Congolese specimens, which may warrant further studies into the potential status of *C. falcifer* as a species complex. The only available *cox*1 sequence (Jorissen et al. 2022) failed to align to our sequences as the amplified regions only marginally overlapped.

#### *Protogyrodactylus* Johnston & Tiegs, 1922

*Protogyrodactylus* spp.

The monopisthocotylans infecting *G. nigri* (Fig. 4) were assigned to *Protogyrodactylus* as the material conformed well to the most recent generic diagnoses (Gussev 1973; Galli and Kritsky 2008), particularly in the presence of an accessory sclerite associated with the tip of the inner root of the ventral anchor (Fig. 4B, D: i), an elongate superficial root usually with small distal hook-like extension and recurved point of the dorsal anchor (Fig. 4B, D: ii), and an MCO with a clockwise-coiled copulatory tube (Fig. 4C, E). Species of *Protogyrodactylus* were previously described from the same locality and host species (Boeger et al. 2012), and these descriptions conformed well to the morphology of the specimens collected in the present study. Specimens belonging to *P. kritskyi* presented a characteristic disc-shaped basal bulb of the MCO and the elongated shape of the anterior projections of the ventral bar (Fig. 4B: iii), whereas specimens belonging to *P. ethiopicus* presented an additional marked heel associated with the basal bulb (Fig. 4C) (Boeger et al. 2012).

**Figure 4.**
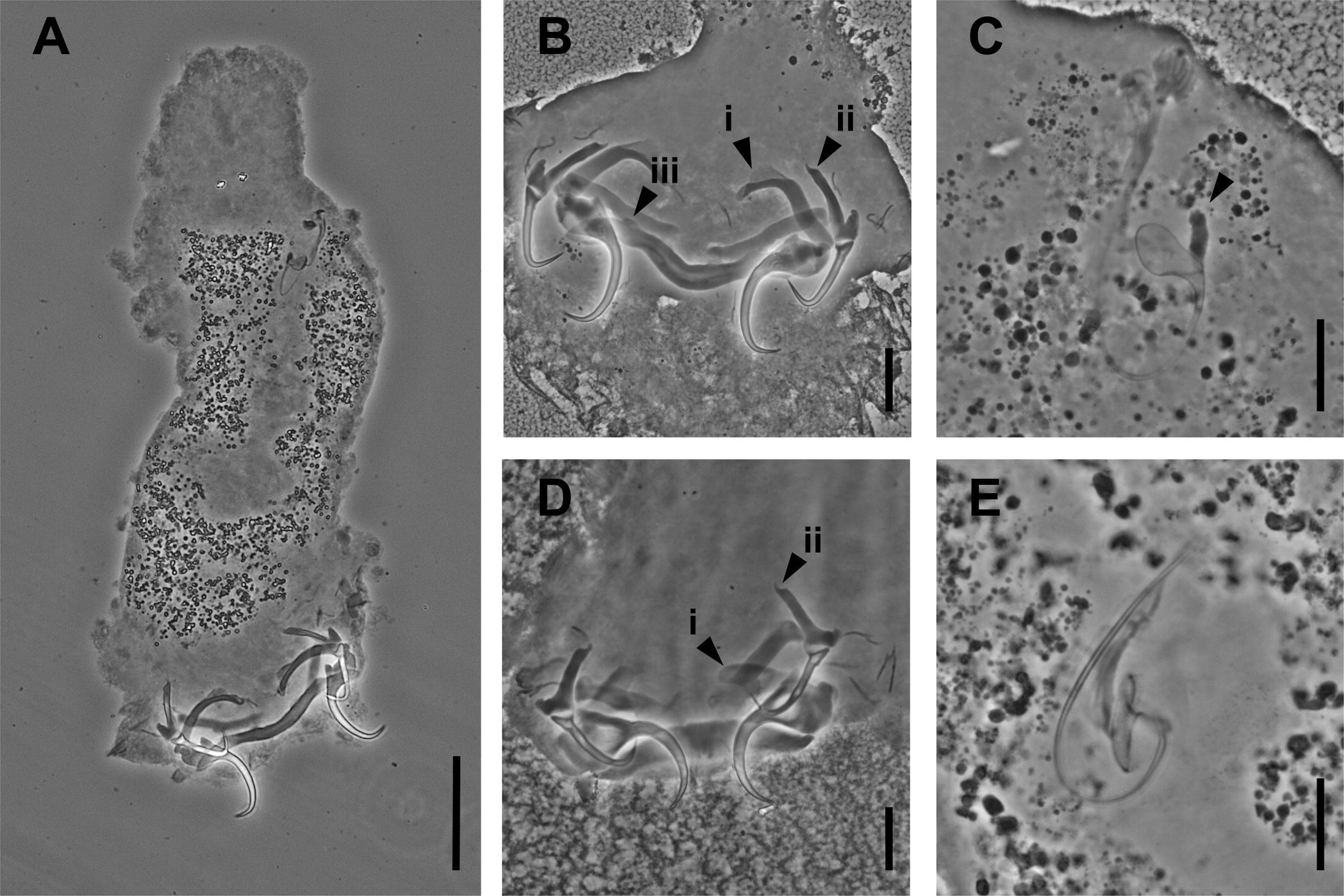
Microphotographs of *Protogyrodactylus* spp. ex *Gerres nigri* using phase contrast. *Protogyrodactylus kritskyi*: **A** Overview of whole body ; **B** haptor sclerites with accessory sclerite associate associated with tip of the inner root of the ventral anchor (i), elongate superficial root usually with small distal hook-like extension and recurved point of the dorsal anchor (ii), and rounded anterior projections of the ventral bar (iii); **C** MCO with disc shaped basal bulb with a marked heel (arrow). *Protogyrodactylus ethiopicus*: **D** haptor sclerites (i) and (ii) as in (Fig. 4B); **E** MCO lacking the marked heel at the basal bulb. Scale bars: 50 µm (**A**); 20 µm (**B**, **D**); 10 µm (**C**, **E**).

Herein, we provide the first molecular data for *P. ethiopicus*. The 28S sequence matched most closely with sequences assigned to two *Protogyrodactylus* sp. 1 (82.6%) and *P.* sp. 2 (84.2%) infecting *Gerres oblongus* Cuvier, 1830 in China, as well as *P. alienus* Bychowsky & Nagibina, 1974 (81.9%) infecting *G. filamentosus* Cuvier, 1829 in China (Wu et al. 2006). Our *cox*1 sequence is the first published for any member of *Protogyrodactylus*. None of the targeted markers could be amplified for *P. kritskyi*.

#### *Euryhaliotrema* Kritsky & Boeger, 2002

*Euryhaliotrema* spp.

The monopisthocotylans found on the gills of *M. sebae* (Fig. 5A–C) were assigned to *Euryhaliotrema* as these specimens conformed well to the original and emended diagnoses of the genus (Kritsky and Boeger 2002; Kritsky 2012), particularly in the vas deferens looping the left intestinal caecum, the presence of ventral and dorsal anchor-bar complexes (Fig. 5D, H, M), a dextral vaginal pore, and the MCO with a coiled (with counterclockwise rings) or meandering tubular copulatory tube, often with a bulbous base (Fig. 5E, H, I, L) and an accessory piece that can be present or absent.

**Figure 5.**
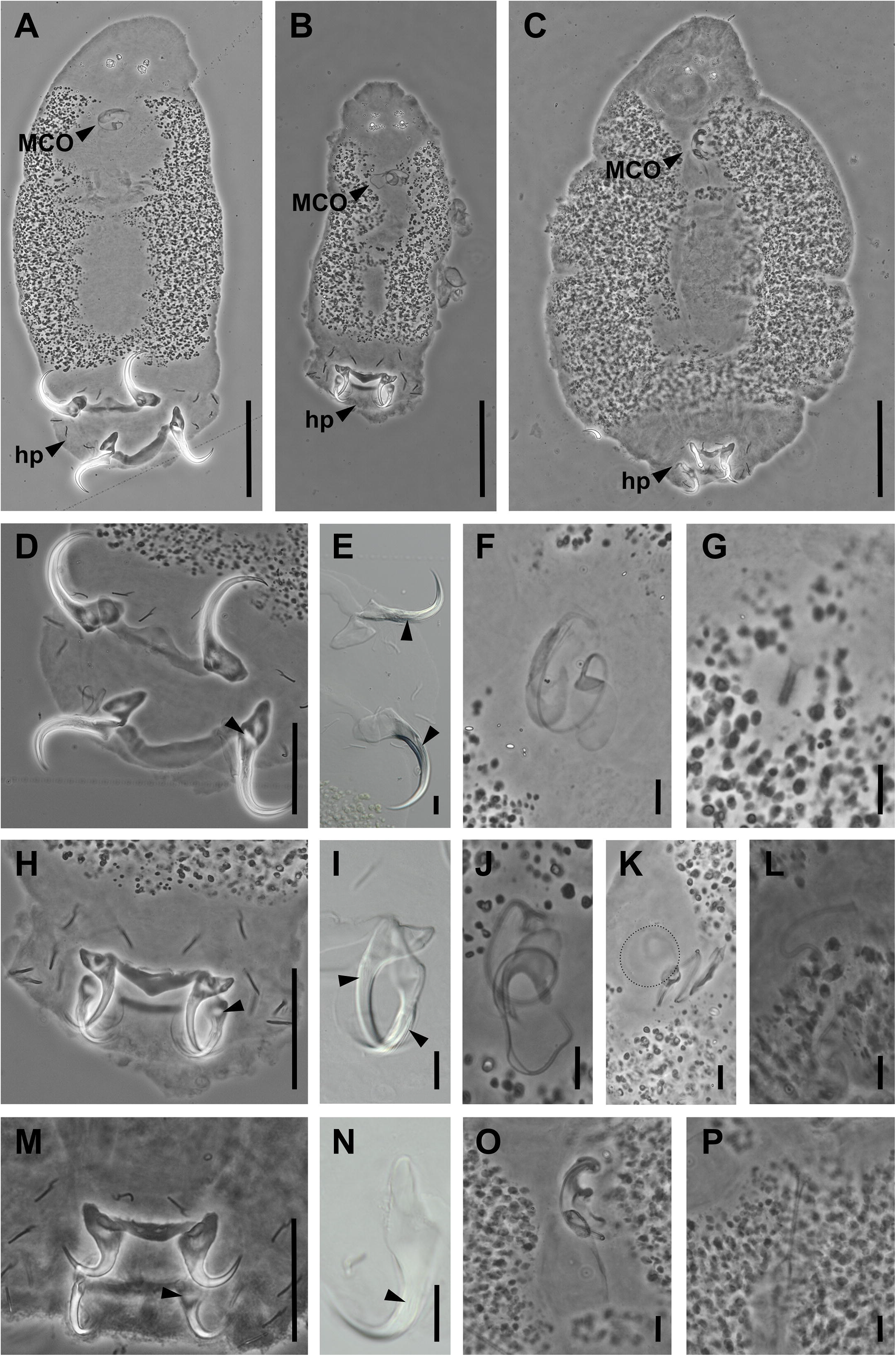
Microphotographs of *Euryhaliotrema* spp. ex *Monodactylus sebae* using phase contrast and differential interference contrast microscopy. Whole body of **A** *Euryhaliotrema* sp. ‘hook’; **B** *Euryhaliotrema* sp. ‘wrench’; and **C** *Euryhaliotrema* sp. ‘bigboy’. *Euryhaliotrema* sp. ‘hook’: **D** haptor; **E** haptoral anchors with longitudinal grooves; **F** male copulatory organ; and **G** sclerotised vagina. *Euryhaliotrema* sp. ‘wrench’: **H** haptor; **I** haptoral anchors with longitudinal grooves; **J** male copulatory organ with **K** basal bulb and accessory piece; and **L** sclerotised vagina. *Euryhaliotrema* sp. ‘bigboy’: **M** haptor; **N** dorsal haptoral anchor with longitudinal grooves; **O** male copulatory organ; and **P** sclerotised vagina. Arrows: thinning in the basal of the dorsal anchors (**D**, **H**, **M**) or longitudinal grooves in anchors (**E**, **I**, **N**). Dotted line (K): outline of weakly sclerotised basal bulb. Scale bars: 100 µm (**A–C**); 50 µm (**D**, **H**, **M**); 10 µm (**E–G**, **I–L**, **N–P**).

Based on the hard parts of the attachment and reproductive organs, we identified three distinct morphospecies that we used for our downstream biodiversity analyses. No species of *Euryhaliotrema* have previously been reported from either the East Atlantic or *M. sebae*, but all three species examined here presented a thinning in the base of the dorsal anchor (Fig. 5D, H, M) similar to species collected from snappers (Lutjanidae) in the Gulf of Mexico including *E. fastigatum* (Zhukov, 1976), *E. cardinale* Kritsky & Justine, 2012, *E. cognatus* Kritsky & Galli, 2012, *E. diplops* Kritsky, Justine & Young, 2012, and *E. tormocleithrum* Kritsky & Galli, 2012, as well as longitudinal grooves along the surfaces of the anchor points and shafts (Fig. 5E, I, N), which have been reported from *E. anecorhizion* Kritsky & Mendoza-Franco, 2012, *E. fastigatum* (Zhukov, 1976), and *E. diplops* Kritsky, Young & Justine, 2012 (Kritsky 2012). *Euryhaliotrema solenophallus* Kritsky, 2019 is the only other species described from a monodactylid host. This species has been described from *M. argenteus* off the Australian coast, but bears little resemblance to its congeners from *M. sebae* by lacking the thinning of the dorsal anchors, longitudinal grooves of both anchor pairs, and an accessory piece (Kritsky 2019). The three species infecting *M. sebae* can be differentiated as follows:

- *Euryhaliotrema* sp. ‘hook’ is characterised by an MCO with a sheath-like and J-shaped accessory piece and a basal bulb of more than half the length of the accessory piece (Fig. 5F) as well as a short tubular vagina (Fig. 5G).
- *Euryhaliotrema* sp. ‘wrench’ is characterised by an MCO with a halfpipe-shaped accessory piece (Fig. 5J) and a basal bulb larger than the accessory piece (Fig. 5K) as well as a long meandering tubular vagina (Fig. 5L).
- *Euryhaliotrema* sp. ‘bigboy’ is characterised by an MCO with a pincer-like accessory piece and a basal bulb approximately 1.5 times the length of the accessory piece (Fig. 5O) as well as a straight delicate, tubular vagina (Fig. 5P). This species was also the largest of the three in terms of length and width, creating a bulky appearance (Fig. 5C).

We acknowledge that to properly delimit these three proposed species, formal descriptions are still required, which would need to include quantitative characters (see Kritsky 2012).

However, we are confident enough in these morphological groups to represent a minimum number of species within our samples, suitable for downstream community analyses.

The molecular data supported the morphological identification of our specimens as members of *Euryhaliotrema*. For 28S, *Euryhaliotrema* sp. ‘hook’ (28S: 93.7%–95.6%), *Euryhaliotrema* sp. ‘wrench’ (28S: 93.9%–95.4%), and *Euryhaliotrema* sp. ‘bigboy’ (28S: 93.4%–95.7%) showed the highest similarity with sequences assigned to *E.* sp. (Zhukov, 1976) from *Lutjanus griseus* in the Gulf of Mexico in a previous study (Mendoza-Franco et al. 2018; see comments on the status of these sequences in ’Material and methods’). For 18S, *Euryhaliotrema* sp. ‘hook’ (97.2%–97.6%), *Euryhaliotrema* sp. ‘wrench’ (98.0%), and *Euryhaliotrema* sp. ‘bigboy’ (96.1%–96.5%) were most similar to a sequence of a specimen annotated as *E*. sp. ‘HQDD’ from *L. rhodopterus* off the coast of China (Wu et al. 2007). For *cox*1, *Euryhaliotrema* sp. ‘hook’ (79.6%) and *Euryhaliotrema* sp. ‘wrench’ (79.8%–82.1%) matched most closely with a sequence assigned to *E. johni* (Tripathi, 1959) from *Lutjanus argentimaculatus* (Forsskål, 1775) off the coast of China (Zhang et al. 2019). The *cox*1 sequence of *Euryhaliotrema* sp. ‘bigboy’ was most similar to a sequence of *E. pirulum* (Plaisance & Kritsky, 2004) (*cox*1: 84.1%) from *Chaetodon vagabundus* from Palau (Plaisance et al. 2008). No ITS1 reference sequences of species of *Euryhaliotrema* were available at the time of writing.

#### Diplectanidae Monticelli, 1903

*Pseudorhabdosynochus Yamaguti, 1958*

*Pseudorhabdosynochus hargisi (Oliver & Paperna, 1984)*

Monopisthocotylan flatworms infecting the gills of *E. aeneus* were assigned to Diplectanidae due to the presence of well-developed anchors, supporting bars, and accessory adhesive plaques (‘squamodiscs’)in the opisthohaptor (Yamaguti 1963; Domingues and Boeger 2008) (Fig. 6A). The specimens corresponded well with the most recent diagnosis of *Pseudorhabdosynochus*, including the four-chambered male copulatory organ (Fig. 6B) and the anterior rows of rodlets in the squamodiscs with closed rings (Fig. 6A) (Domingues and Boeger 2008).

**Figure 6.**
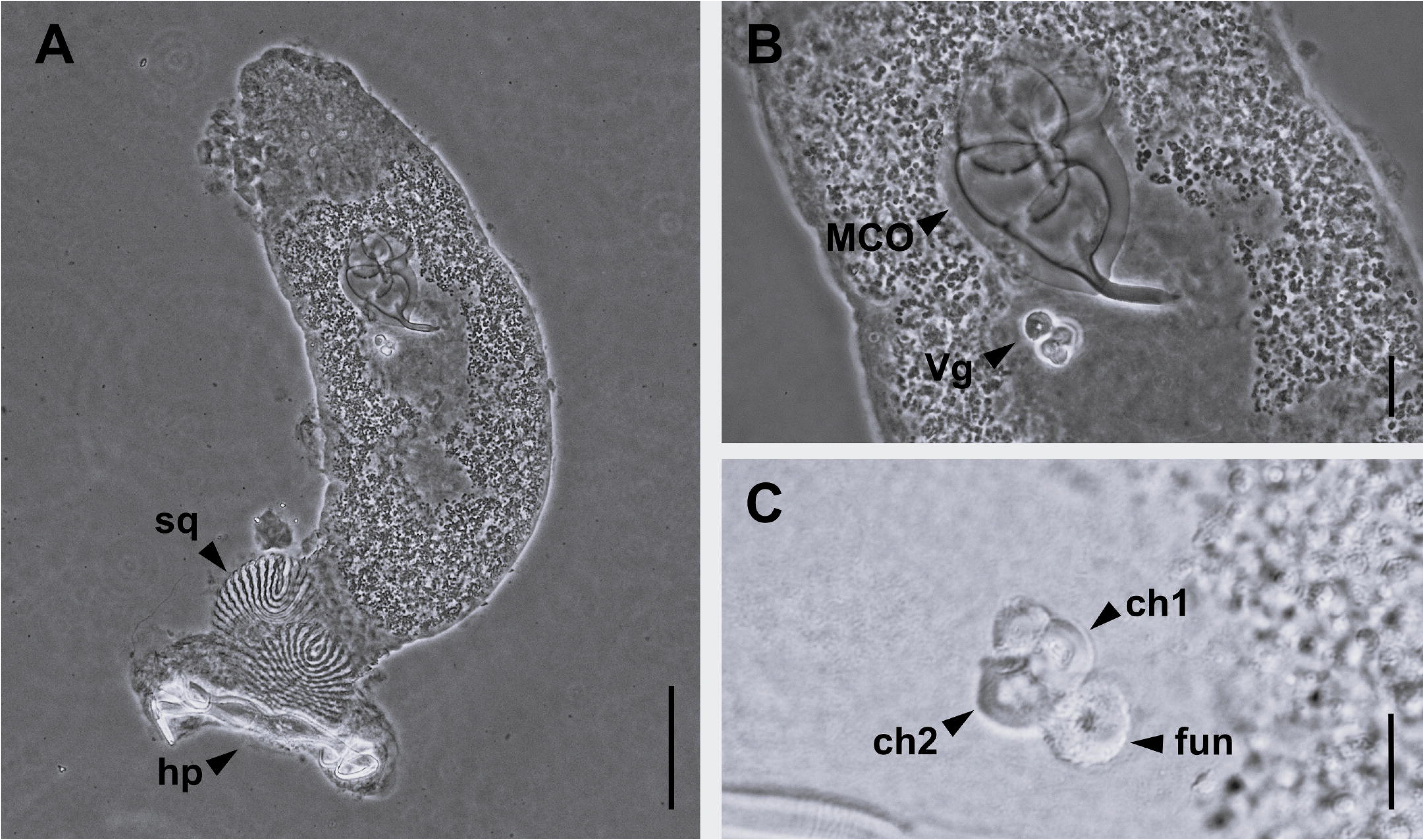
Microphotographs of *Pseudorhabdosynochus hargisi* ex *Epinephelus aeneus* using phase contrast and differential interference contrast. **A** Whole body of specimen with squamodiscs (sq) and haptor (hp) indicated; **B** reproductive organs including chambered male copulatory organ (MCO) and sclerotised vaginal (Vg); **C** vaginal sclerite, with two thick-walled chambers (ch1, ch2) and distal funnel (fun) highlighted. Scale bars: 100 µm (**A**); 20 µm (**B**); 10 µm (**C**).

Our material is morphologically most similar to *P. hargisi* and *P. americanus* Price, 1937 in possessing a sclerotised vagina with a distal funnel and two large thick-walled chambers (Oliver and Paperna 1984; Kritsky et al. 2015) (Fig. 6C). Due to their morphological similarity, *P. hargisi* was previously synonymised with *P. americanus* (Tingbao et al. 2005). However, the synonymy was later rejected by Kritsky et al. (2015) due to the lack of ‘diagnostic’ drawings of the vaginal sclerites of *P. hargisi* (Oliver and Paperna 1984; Tingbao et al. 2005), as well as the distinct ‘confirmed’ (i.e., vouchered) host and geographical records, with *P. americanum* reported from *Epinephelus itajara* (Lichtenstein, 1822) in the West Atlantic (Kritsky et al. 2015) and *P. hargisi* reported from *E. aeneus* from the eastern Mediterranean (Oliver and Paperna 1984). As our material was found on *E. aeneus* in the Eastern Atlantic and was indistinguishable from the description of *P. hargisi*, we assigned these specimens to this species. However, as noted by Kritsky et al. (2015), *P. hargisi* requires redescription as the existing drawings of the vaginal sclerites are of insufficient quality to be of diagnostic value.

We provide the first molecular data for *P. hargisi*, including partial 18S, ITS1, ITS2, 28S, and *cox*1 sequences. The 18S and 28S sequences most closely resembled those of other members of Diplectanidae infecting groupers (Serranidae), including *Cryptoplectanum grouperi* (Bu, Leong, Wong, Woo & Foo, 1999) (18S: 96.7%–96.9%, 28S: 87.1%–88.0%) from *Epinephelus coioides* (Hamilton, 1822) off the coast of China (Wu et al. 2005; Tingbao et al. 2006) and *P. lantauensis* (Beverley-Burton & Suriano, 1981) (18S: 98.4%) from *Epinephelus brunneus* Bloch, 1793 in China (Wu et al. 2005). Due to the poor taxonomic coverage of ITS1 and ITS2 sequences for these species and other members of *Pseudorhabdosynochus*, our sequences showed no close matches (>90%) with any reference sequences on GenBank. The *cox*1 sequences matched most closely with sequences of *P. sosia* Neifar & Euzet, 2007 (86.9%) from *Epinephelus costae* (Steindachner, 1878) off the coast of Tunisia (Chaabane et al. 2016).

*Pseudorhabdosynochus hargisi* was described from the same host species in the Eastern Mediterranean (Oliver and Paperna 1984), and was later reported off the coast of “south-west Africa” (no precise locality was specified) (Aljoshkina 1984) as *P. americanus* (later reassigned to *P. hargisi*; Kritsky et al. 2015). Our report, therefore, constitutes a new geographical record for this species.

### Polyopisthocotyla Brabec, Salomaki, Scholz & Kuchta, 2023

Mazocraeidea Bychowsky, 1957

Rhinecotylidae Lebedev, 1979 *Rhinecotyle* Euzet & Trilles, 1960 *Rhinecotyle* aff. *crepitacula*

Polyopisthocotylan flatworms found on the gills of *S. afra* (Fig. 7A–C) were identified as members of *Rhinecotyle*, based on the possession of an oblong haptor with a ventral concavity divided by a longitudinal septum and an additional series of transverse muscular septa, creating the appearance of two parallel rows of loculi (Fig. 7A) (Euzet and Trilles 1960; Euzet and Wahl 1970). Our material corresponds well to the redescription of *R. crepitacula* sensu Euzet and Wahl (1970) on *S. afra*. As noted by Bouguerche (2025), the specimens from the original description collected in the Mediterranean off *S. sphyraena* (Euzet and Trilles 1960) differ from the specimens collected by Euzet & Wahl (1970) from *S. afra* (cited as *S. piscatorum*) off the coast of Côte d’Ivoire with regard to the number and size of clamps and the number of hooks in the genital atrium (Fig. 7B) (see Bouguerche 2025). Our specimens morphologically correspond to those reported from Côte d’Ivoire. Given these differences, we refer to our specimens as *Rhinecotyle* aff. *crepitacula*, pending an in-depth morphological and molecular study of the populations infecting *S. afra*.

**Figure 7.**
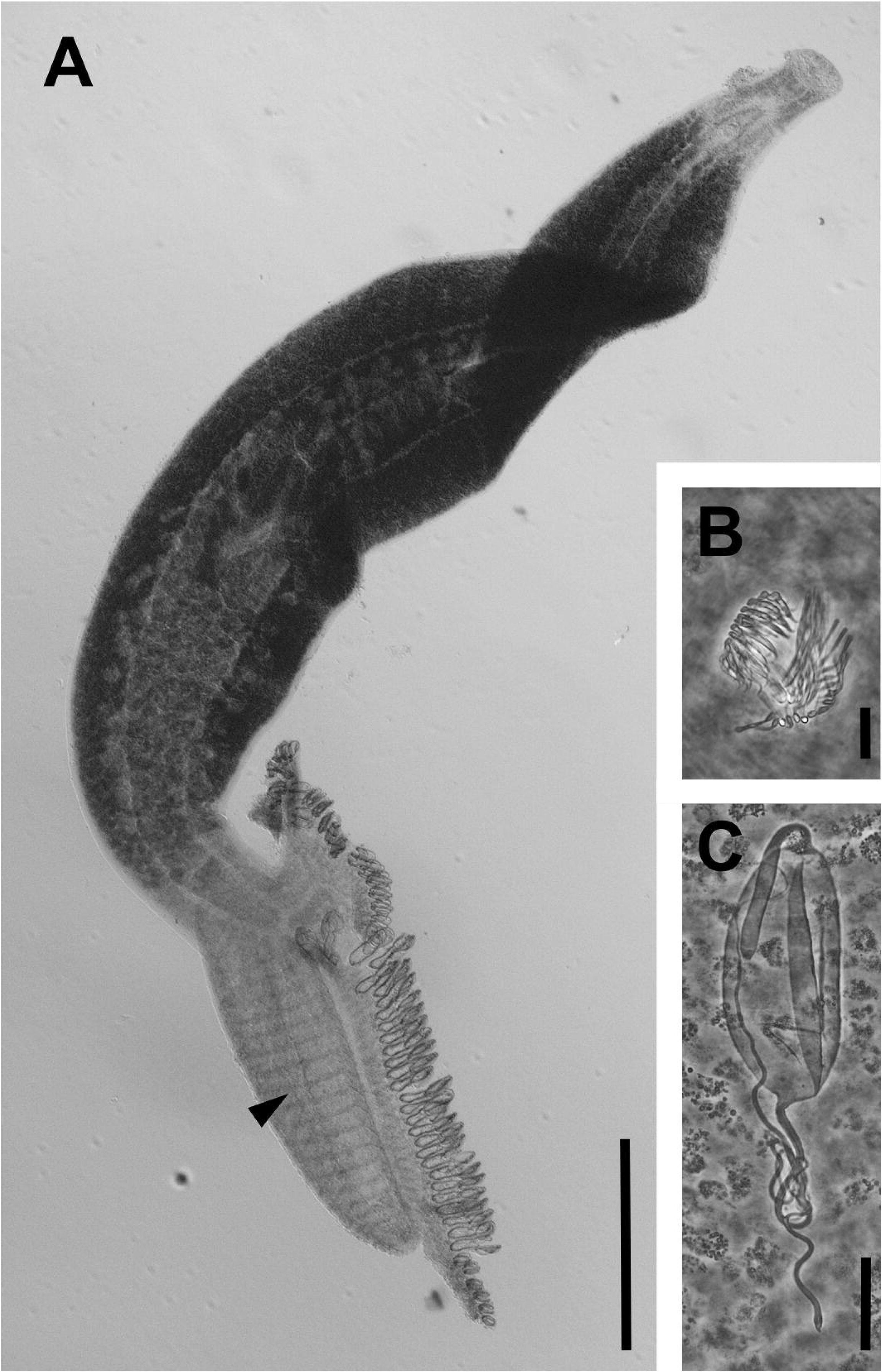
Microphotographs of *Rhinecotyle* aff. *crepitacula* ex *Sphyraena afra* using differential interference contrast and phase contrast: **A** whole body with septum in haptor highlighted by arrow; **B** genital atrium with many spines; **C** egg. Scale bar: 500 µm (**A**); 20 µm (**B**); 50 µm (**C**).

We obtained the first DNA sequences for species of *Rhinecotyle*. Our sequences were most similar to those assigned to other mazocraeidean species, including *Atriaster ibamba* Vermaak, Bouguerche, Acosta & Smit, 2025 (Microcotylidae) (28S: 90.7%) from *Diplodus capensis* in South Africa (Vermaak et al. 2025), *Microcotyle erythrinii* Van Beneden & Hesse, 1863 (Microcotylidae) (18S: 94.9%-95.1%) from *Pagellus erythrinus* (L., 1758) in France (Sinnappah et al. 2001), *Zeuxapta seriolae* (Meserve, 1938) (Heteraxinidae) (ITS1: 73.2%) from *Seriola lalandi* Valenciennes, 1833 in Brazil (Camargo and Santos 2020), *Diplostamenides sciaenae* (Goto, 1984) (ITS2: 74.1%) from *Argyrosomus japonicus* (Temminck and Schlegel, 1843) in Australia (Hayward et al. 2007), and *Pyragraphorus rohdei* Kritsky & Martin, 2026 (ITS2: 74.5%) from *Trachinotus baillonii* (Lacépède, 1801) (Kritsky and Martin 2026).

### Trematoda Rudolphi, 1808

Hemiuroidea Looss, 1899

Within the study context, trematodes presented specific identification challenges. To preserve a voucher specimen while obtaining DNA (i.e., hologenophore), we removed the posterior portion of each organism for molecular analysis. Although the posterior region contains key morphological characters, removing any tissue section inherently impacts morphological diagnostic capacity. Consequently, species identification relied heavily on matching DNA sequences to reference data in GenBank (a strategy similarly applied to nematodes). Because all sequenced trematodes closely matched GenBank reference sequences for Hemiuroidea, we were able to use established taxonomic keys for this group (detailed below) to confirm specimen identities.

Lecithophyllidae Skrjabin & Guschanskaja, 1954

*Aponurus* Looss, 1907

*Aponurus* cf. *laguncula*

Several specimens collected from the gills of *M. sebae* were assigned to Lecithophyllidae, following the recently proposed diagnosis and revised classification of this family (Vainutis and Voronova 2025). The specimens collected herein conform well to the generic diagnosis of *Aponurus* as provided by Gibson and Bray (1986), Gibson (2002b), and Vainutis and Voronova (2025), particularly in possessing two testes, an oval to globular ovary, globular vitelline lobes, and a small genital atrium (Fig. 8A). Morphologically, our material is most similar to *A. laguncula*, originally described by Looss (1907) from the carangid *Lichia amia* (L., 1758), the sparid *Spicara maena* (L., 1758) (= *Maena vulgaris* Valenciennes, 1830), the engraulid *Engraulis encrasicolus* (L., 1758), the belonid *Belone belone* L., 1761 (= *Belone acus* Risso, 1827), and other unidentified fishes from Trieste, Italy, sharing the following characters: a ventral sucker approximately twice as large as the oral sucker; obliquely arranged testes located in the anterior part of the hindbody; an oval sinus sac; a seminal vesicle positioned anteriorly or slightly reaching the anterior margin of the ventral sucker (Fig. 8A); and eggs of similar size, narrow and more or less pointed at the opercular end (27.8 µm × 14.8 µm *vs*. 27 µm × 16 µm).

**Figure 8.**
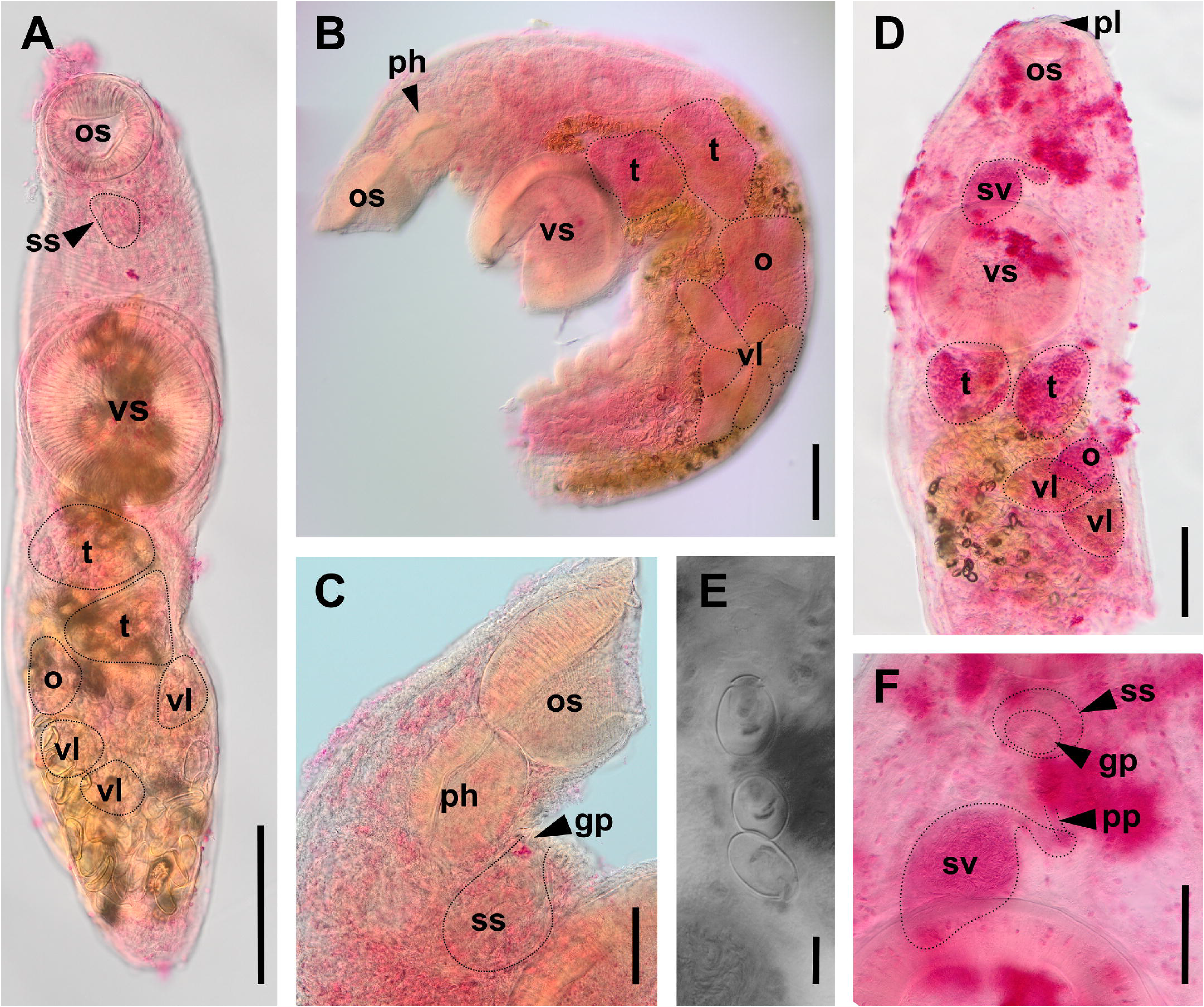
Microphotographs of trematodes using differential interference contrast microscopy. *Aponurus* cf. *laguncula*: **A** whole specimen. *Lecithaster sayori*: **B** partial specimen and **C** forebody. *Lecithochirium floridense*: **D** partial specimen; **E** eggs; and **F** genitalia. Abbreviations: gp – genital pore, os – orals sucker, o – overary, ph – pharynx, pl – pre-oral lobe, pp – *pars prostatica*, ss – sinus sac, sv – seminal vesicle, t – testes, vl – vitellarian lobe, vs – ventral sucker. Scale bars: 100 µm (**A**, **B**, **D**); 50 µm (**C**, **F**); 10 µm (**E**).

*Aponurus laguncula* has been reported from a wide range of hosts belonging to multiple families and orders and is characterised by considerable intraspecific morphological variation (Bray and MacKenzie 1990; Madhavi and Bray 2018). Bray and MacKenzie (1990) provided a comprehensive account of its host repertories and geographic distribution, noting that several congeners, although currently regarded as valid species, exhibit pronounced morphological similarity to *A. laguncula*. Subsequent studies have raised concerns regarding the taxonomic boundaries of this species. In particular, evidence for cryptic diversity within *Aponurus* has been reported from the western Mediterranean (Carreras-Aubets et al. 2011), suggesting that what is currently recognised as *A. laguncula* may comprise a complex of morphologically similar but genetically distinct lineages. In addition, morphological variation observed among specimens attributed to *A. laguncula* has been highlighted as further complicating species delimitation, indicating that species boundaries within this taxon may be less clear-cut than is traditionally assumed (Madhavi and Bray 2018).

In this context, the relatively low nucleotide identity observed in the 28S rDNA with sequences identified as *A. laguncula* by Claxton et al. (2017) from *Rhomboplites aurorubens* (Cuvier, 1829) in the Gulf of Mexico (96.5%–96.8%) is noteworthy. In fact, our sequences were most similar to those of *Aponurus mulli* Carreras-Aubets, Repulles-Albelda, Kostadinova & Carrasson, 2011 from *Mullus surmuletus* (L., 1758) (97.3%–97.6%) and *M. barbatus* (L., 1758) (97.3%–97.4%) (Pankov et al. 2006; Carreras-Aubets et al. 2011) from the Mediterranean. The ITS2 sequences were also more similar (96.5%) to those of *A. mulli* from *M. surmuletus* (Pankov et al. 2006) than to those of *A. laguncula* from the Gulf of Mexico (Claxton et al. 2017) (95.9%). However, a robust interpretation of these molecular differences is constrained by the lack of corresponding morphological information for some of the DNA sequence data attributed to *A. laguncula*, thereby limiting any meaningful comparative assessments. Furthermore, *A. mulli* presents eggs with a rounded shape (Carreras-Aubets et al. 2011), unlike the eggs with a pointy shape at the opercular end found in our present specimens (Fig. 8E). Given the unresolved taxonomic boundaries of *A. laguncula* and relatively low nucleotide identity, we refer to our specimens as *A.* cf. *laguncula*.

We would also like to point out the low nucleotide identity of our 28S sequences and those of *A. laguncula* published by Claxton et al. (2017) from those of Machado et al. (2024) from *Chaetodipterus faber* from the Brazilian Southeast (87.3%–87.6%) and those of Martínez-Flores et al. (2023) from *Trachinotus rhodopus* Gill, 1863 from the Mexican Pacific coast (85.0%–85.6%), which call into question the identification of these specimens for the latter two, given that these authors, unlike Claxton et al. (2017), did not provide a detailed morphological characterisation. Moreover, the images provided in Machado et al. (2024) show a larger distance between the posterior margin of the ventral sucker and the testes compared to previous reports (Bray and MacKenzie 1990), adding further doubt to their identification.

Lecithophyllids are known to parasitise the stomach and intestines of marine teleosts (Vainutis and Voronova 2025). The specimens collected in the present study were found exclusively associated with the gills and were absent from both the stomach and intestines, which may reflect *post-mortem* displacement from the original site of infection. A similar finding was reported for *A. balisti* Szuks, 1981 from *Balistes capriscus* Gmelin, 1789 from Mauritania (Szuks 1981). This species was included in a list of potential synonyms of *A. laguncula* (Bray and MacKenzie 1990).

*Aponurus laguncula* has been reported across the globe from the Atlantic, Pacific, and Indian Oceans (Bray and MacKenzie 1990; Machado et al. 2024). No previous records of *A. laguncula* from monodactylid hosts exist. Therefore, the present record of *Aponurus* cf. *laguncula* infecting *M. sebae* off the Senegalese coast potentially represents a new host and geographical record.

Lecithasteridae Odhner, 1905

*Lecithaster* Lühe, 1901

*Lecithaster sayori* Yamaguti, 1938

The specimens examined in the present study were assigned to Lecithasteridae, following the diagnosis of Gibson (2002b). The specimens collected herein conform well to the generic diagnosis of *Lecithaster* as provided by Gibson and Bray (1986) and Vainutis and Voronova (2025), particularly in possessing two testes, a lobed ovary with four lobes, tear-shaped vitelline lobes located in the anterior third to middle of the hindbody (Fig. 8B), and a proximal portion of the seminal vesicle situated posterior to the ventral sucker (Fig. 8C).

Based on the generic key (Vainutis and Voronova 2025), our specimens most closely resembled *L. sayori* as described by Yamaguti (1938) from the intestine of *Hyporhamphus sajori* (Temminck & Schlegel, 1846) in Lake Hamana, Japan, because the genital pore was positioned ventral to the pharynx (Fig. 8C), the testes were oval-shaped, and the seminal vesicle was posterodorsal to the ventral sucker (Fig. 8C). The only members of *Lecithaster* reported from West Africa are *L. africanus* Fischthal & Thomas, 1971 and *L. ghanensis* Fischthal & Thomas, 1971, which were described from Ghana (Fischthal and Thomas 1971), but the genital pore in both species is situated on the posterior margin of the oral sucker. The general morphology of our specimens fits the description (Yamaguti 1938) and redescription of *L. sayori* (Atopkin et al. 2020). However, our specimens differed in some characteristics. Notably, the sucker width ratio was larger, with 1.6–2.1 *vs.* 1.1, for specimens collected in Vietnam (Atopkin et al. 2020). This ratio resembles specimens reported as *L. stellatus* by Bray et al. (1993). However, these specimens are also likely representatives of *L. sayori* due to the positioning of vitellarium, ovaries, and genital pore as remarked by Atopkin et al. (2020), suggesting that sucker ratios might be somewhat variable for this species. The identification as *L. sayori* was further supported by the molecular data, which matched closely with the sequences of *L. sayori* from Vietnam (Atopkin et al. 2020) (28S: 97.9%–99.2%; ITS2: 97.5%–97.7%). The *cox*1 sequences we provided did not match with the only reference sequence, possibly because the region of *cox*1 targeted by our primers differed from that of Atopkin et al. (2020). Considering their morphological similarities alongside the relatively low sequence divergence between our specimen and the Vietnamese specimens, we consider them conspecific.

Prior to our study, *L. sayori* was reported as a parasite of fishes belonging to Beloniformes. Other than the original description, the species was recorded in *Cololabis saira* (Brevoort, 1856) (Scomberesocidae) from the Pacific off the coast of Hokkaido, Japan (Machida et al. 1972); in *Strongylura anastomella* (Valenciennes, 1846) (= *Ablennes anastomella*) (Belonidae) from the Yellow Sea and the Bohai Sea (Shen and Qiu 1995); and in *Hemiramphus marginatus* (Forsskål, 1775) (Hemiramphidae) from coastal waters off northern Vietnam (Atopkin et al. 2020). Our records of *L. sayori* infecting *M. sebae* and *G. nigri* off the Senegalese coast represent new host and geographical records for this species. The material was collected from the stomachs of one specimen of *M. sebae* and one of *G. nigri*.

Hemiuridae Looss, 1899

*Lecithochirium* Lühe, 1901

*Lecithochirium floridense* (Manter, 1934)

A single specimen belonging to *Lecithochirium* was found on the gills of *M. sebae*. The specimen corresponds well with the generic diagnosis of *Lecithochirium* by Gibson and Bray (1986) and Gibson (2002a) in having a pre-oral lobe (Fig. 8D), tubular *pars prostatica*, vitellarium of two lateral masses divided into three or four short lobes (Fig. 8F), and eggs without polar filament (Fig. 8E). The specimen appears to belong to the “musculus” group based on the absence of the internal elevations in ventral sucker, the absence of the presomatic pit, the presence of the vitellarium of compact masses with distinct and short digitiform lobes (Fig. 8D), the terminal genitalia of the “musculus” type, and the non-muscular seminal vesicle (Fig. 8F) according to a key for *Lecithochirium* (Bray 1991). The species-level identification was based on morphological characters consistent with the original description of *L. floridense* (Manter 1947), based on material from the stomach of *Paralichthys* sp. (Pleuronectiformes: Paralichthyidae) and other marine fishes from Florida, USA, mainly in possessing a preoral lobe (Fig. 8D), a genital pore at the level of the pharynx (Fig. 8F), two obliquely symmetrical testes (Fig. 8D), and a bipartite seminal vesicle (Fig. 8F). This identification was further supported through the 28S rDNA sequences that showed high identity (99.5%–99.9%) with sequences from specimens collected in the Caribbean (Claxton et al. 2017; Vidal-Martínez et al. 2019) and off the coast of Brazil (Pantoja et al. 2022). Furthermore, the ITS2 sequence obtained from the specimen confirmed this identification (99% match through BLAST) with the specimens collected off the coast of Brazil.

Most records of this species are from the western Atlantic Ocean. The present record of *L. floridense* in *M. sebae* from the coast of Senegal, in the north-eastern Atlantic, represents an extension of the known geographical range of this species and constitutes a new host record. The inclusion of this monodactylid host increases the number of fish families known to harbour this species from 17 to 18 (Pantoja and Kudlai 2022).

Tubulovesicula Yamaguti, 1934

Tubulovesicula sp. 1

Two specimens recovered in the present study were identified to the genus level based on the 28S, ITS2, and *cox*1 DNA barcodes. However, the condition of the specimens did not allow a detailed morphological examination, precluding reliable species-level identification.

Trematodes belonging to *Tubulovesicula* parasitise a wide range of marine fishes and, occasionally, marine snakes, with records spanning the Atlantic, Indian and Pacific Oceans (Gibson 2002b; Martin et al. 2023). According to the World Register of Marine Species (WoRMS) (https://marinespecies.org/, accessed: 17^th^ July 2026), the genus currently comprises 25 valid species. Despite this diversity, molecular data are available for only two species, *Tubulovesicula laticaudi* Parukhin, 1969 and *T. lindbergi* (Layman, 1930).

Alignments with BLAST showed high levels of sequence identity (28S: 99.4%–99.7%; ITS: 99.7%–99.8%; *cox*1: 89.8%–97.8%) between our material and *T. laticaudi* collected from marine elapid snakes in Sri Lanka (Martin et al. 2023), suggesting that the Senegalese specimens are closely related. However, because no detailed morphological description is provided in the present study, the specimens are conservatively identified at the generic level. Three specimens belonging to *Tubulovesicula* were collected from the gills of *H. fasciatus* and *G. nigri*. Previously, only *T. lindbergi* was reported from the Atlantic, including Brazil (Pantoja et al. 2024), Ghana (Fischthal and Thomas 1972), Nigeria (Siddiqi and Hafeezullah 1975), and Puerto Rico (Siddiqi and Cable 1960). However, the conspecificity of these Atlantic specimens with the North Pacific specimens (including the type locality) has been questioned recently based on *cox*1 data (Sokolov et al. 2026).

Plagiorchiida La Rue, 1957 Microphallidae Ward, 1901

Microphallidae gen. sp. 1

Three small specimens of larval trematodes were found encysted in the muscle tissue of *C. amnicola*. Using an ITS2 barcode, we identified these specimens as members of Microphallidae, with 87.9% identity with a specimen assigned to *Longiductotrema tethepae* Kudlai, Cribb & Cutmore, 2016 (Kudlai et al. 2016) and 86.1% identity with a specimen assigned to *Microphallus calidris* Belopolskaja, 1963 (Galaktionov et al. 2012). We also obtained a *cox*1 barcode, but no reference sequences available on GenBank showed > 90% identity. The small size of the specimens and their use as genetic material did not allow for any morphological examination. To our knowledge, microphallids have not been studied in West Africa. However, these species use crustaceans as intermediate hosts – which corresponds to our record – and primarily birds as definitive hosts (Bray et al. 2008). Our DNA sequence, therefore, constitutes the first evidence of the presence of this family of trematodes in the region, and the molecular data may contribute to understanding the life cycle of this group in future studies.

### Cestoda Rudolphi, 1808

#### Trypanorhyncha Diesing, 1863 Trypanorhyncha gen. sp. 1

One small encysted larval cestode was found in the testes of *E. fimbriata*. Using 18S and 28S barcodes, we identified the specimen as a member of Trypanorhyncha, with 95.9% identity with 28S reference sequences assigned to *Pterobothrium kingstoni* Campbell & Beveridge, 1996 from *Hypanus americanus* (Hildebrand & Schroeder, 1928) in Mexico (Adán-Torres et al. 2025) and 95.3% with *Grillotellia exilis* (Linton, 1909) from *Scomberomorus commerson* (Lacepède, 1800) in Indonesia (Palm et al. 2009), and 95.5% with 18S reference sequences assigned to Australian specimens of *Grillotia pristiophori* Beveridge & Campbell, 2001 from *Pristiophorus nudipinnis* Günther, 1870 (Olson et al. 2010), *Pterobothrium lintoni* (MacCallum, 1916) from *Choerodon venustus* (De Vis, 1884) (Littlewood and Olson 2001), and *P. platycephalum* (Shipley & Hornell, 1906) from *Himantura* sp. (Olson et al. 2010). We also obtained ITS2 and *cox*1 barcodes, but no reference sequences available on GenBank were similar (> 90% identity). The immature state of the specimen did not allow for any morphological examination.

*Acanthocephala Rudolphi, 1808* Eoacanthocephala Van Cleave, 1936 Gyracanthocephala Van Cleave, 1936 Quadrigyridae Van Cleave, 1920 *Acanthogyrus* Thapar, 1927 *Acanthosentis* Verma & Datta, 1929

Acanthogyrus (Acanthosentis) tilapiae (Baylis, 1948)

The acanthocephalan specimens collected from the intestines of *C. guineensis* and *S. melanotheron* were assigned to Quadrigyridae based on the small proboscis (Fig. 9A, E) with few radially arranged hooks, and trunk with spines, and fish definitive hosts (Amin 1987). We identified our specimens (Fig. 9) as belonging to *Acanthosentis* – a subgenus of *Acanthogyrus* – because the proboscis was armed with three irregularly arranged circles of six rooted hooks each (Amin 2005). Our material most closely resembled *A.* (*A.*) *tilapiae* (Baylis, 1947), described from *Oreochromis lidole* (Trewavas, 1941) in Lake Malawi (Baylis, 1947) in the size of the anterior (35–37 µm, root: 19–23 µm), middle (14–18 µm), and posterior hooks (14–18 µm), as well as the number of rows (29–36) and length (2.9–4.0 µm) of the body spines (Fig. 9C). As noted by Louizi et al. (2022), five species currently included in *Acanthogyrus* (*Acanthosentis*) have been reported from fishes in Africa: *A.* (*A.*) *maroccanus* (Dollfus, 1951)*, A.* (*A.*) *nigeriensis* (Dollfus & Golvan, 1956), *A.* (*A.*) *malawiensis* Amin & Hendrix, 1999, *A.* (*A.*) *papilio* Troncy & Vassiliadès, 1974, and *A*. (*A*.) *tilapiae*. Our material differs from *A.* (*A.*) *maroccanus* and *A.* (*A.*) *nigeriensis* in having smaller proboscis hooks and spines. Furthermore, our material has a larger number of rows of body spines than *A.* (*A.*) *maroccanus*. Our material further differs from *A.* (*A.*) *malawiensis* in having middle hooks that are smaller than the anterior hooks. *Acanthogyrus* (*A.*) *papilio* Troncy & Vassiliadès, 1974 described from *P. barbarus* is possibly a synonym of *A.* (*A.*) *tilapiae* (Louizi et al. 2022). However, the original description of *A.* (*A.*) *papilio* was partially based on immature specimens (Troncy and Vassiliadès 1974) and the type material appears to be lost (Louizi et al. 2022). Therefore, the status of this species will have to be verified through new specimens collected from the type host and location.

**Figure 9.**
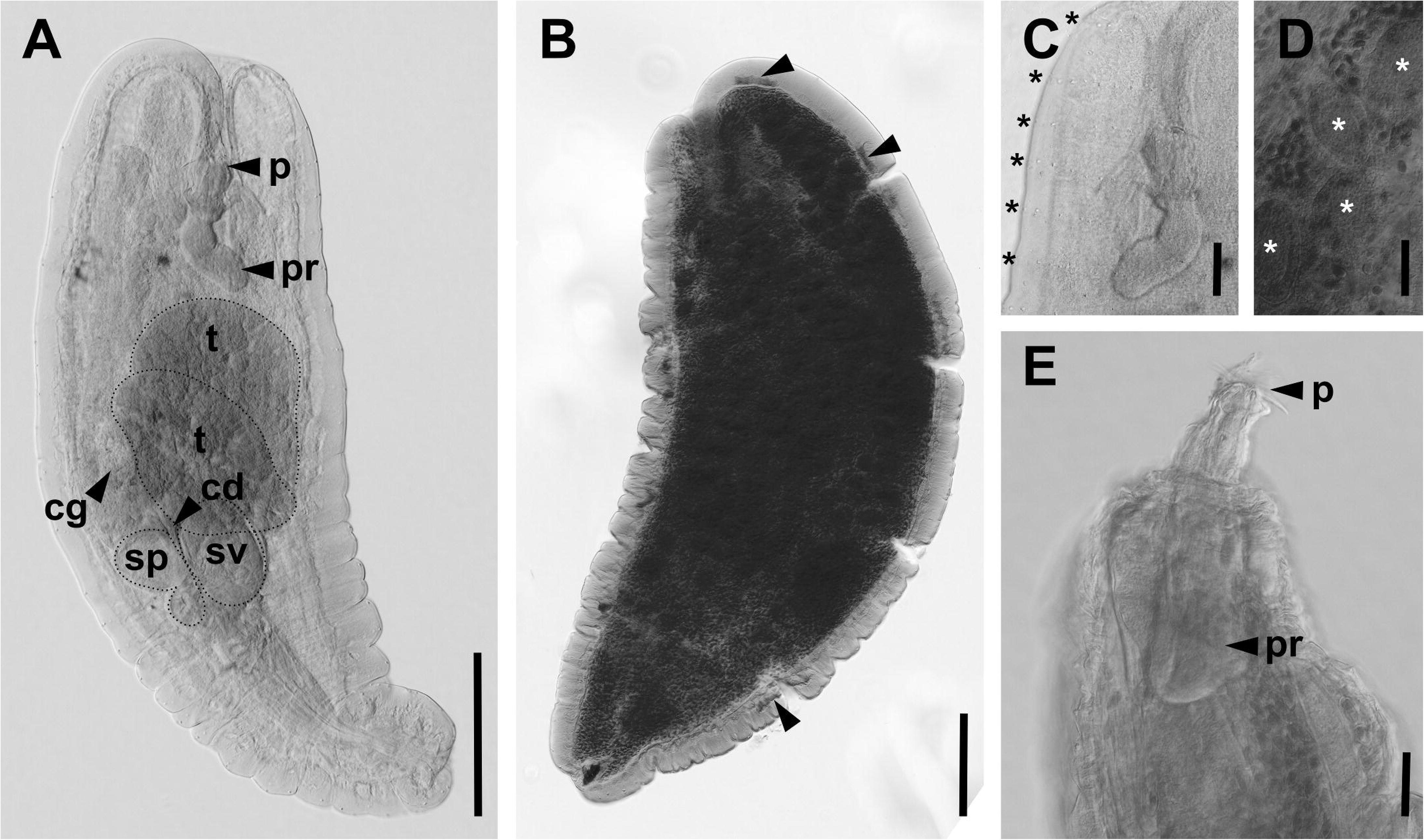
Microphotographs of *Acanthogyrus tilapiae.* **A** Whole body of male specimen with inverted proboscis; **B** whole body of gravid female with giant nuclei indicated by arrows; **C** anterior part of male tegument with rows of body spines; **D** ovarian balls in female; and **E** everted proboscis of male. Abbreviations: cg – cement duct, cg – cement glands, p – proboscis, pr – proboscis receptacle, sp – Saefftigen’s pouches, sv – seminal vesicle. Scale bars: 200 µm (**A**, **B**); 50 µm (**C–E**).

Based on the morphological resemblance, *A.* (*A.*) *tilapiae* appears as the most closely corresponding species and, therefore, we considered our material conspecific. However, we recognise that this species has previously been considered a species complex (Louizi et al. 2022) because of its reported geographical range across the African continent and its host repertoire including around 40 species of mostly African cichlids (Kuchta et al. 2018).

The molecular data of the 28S and 18S regions confirmed the affiliation of our material with specimens identified as *A.* (*A.*) *tilapiae*. Specifically, our sequences matched (18S: 99.9%, 28S: 96.9%–100%) those of specimens of *A.* (*A.*) *tilapiae* collected from *C. guineensis* and *Oreochromis aureus* (Steindachner, 1864) in Morocco (Louizi et al. 2022). Amplification of the *cox*1 sequences using the LCO1490/HCO2198 primer pair (Lisitsyna et al. 2025) was not successful for our material. However, we obtained partial sequences using the ASmit1/Schisto3/ASmit2 primers that aligned with a mitochondrial genome of *Pallisentis* (*Pallisentis*) *celatus* (Van Cleave, 1928) (Pan and Nie 2013), a member of the same family. Given these difficulties in obtaining sequences, we suggest that primers for this region may have to be developed specifically for *Acanthogyrus* to improve amplification success in the future. However, we note that the 28S sequence of one of our specimens diverged from the others (97.8%), which may indicate the presence of multiple lineages (or cryptic species) of *A.* (*A.*) *tilapiae* in the Saloum Delta.

Only a single definitive record of *A.* (*A.*) *tilapiae* exists for West Africa, from *C. zillii* in Nigeria (Shotter 1974). Our material constitutes the first explicit record of *A.* (*A.*) *tilapiae* in Senegal. However, two studies on the ultrastructure of acanthocephalans reported *A.* (*A.*) *tilapiae* from *S. melanotheron* (Marchand and Mattei 1976; Marchand 1984). Because they used material extracted from the hosts immediately after death, and because these authors were based at Senegalese institutions (see remarks in Table 4), we consider these the first records of *A.* (*A.*) *tilapiae* in Senegal, even though specific sampling sites were not detailed.

Furthermore, *A.* (*A.*) *papilio* Troncy & Vassiliadès, 1974, described from *P. barbarus* may constitute another potential record from Senegal, as the species has been suggested to be a synonym of *A.* (*A.*) *tilapiae* (Louizi et al. 2022; see comments above regarding its uncertain status). We found no specimens belonging to *Acanthosentis* in the intestines of *P. barbarus* in the present study.

### Arthropoda Gravenhorst, 1843

Copepoda Milne Edwards, 1840

Siphonostomatoida Thorell, 1859

Caligidae Burmeister, 1835

*Caligus* Müller O.F., 1785

All copepod specimens were assigned to Caligidae based on their dorsoventrally flattened bodies, the cephalothorax covered by a subcircular cephalothoracic shield, and the third legs being fused with a broad intercoxal sclerite to form an apron, which closes the posterior border of the cephalothoracic shield (Boxshall and Halsey 2004). All specimens were further identified as members of *Caligus* based on a recently published key for the family (Bernot et al. 2025). Specifically, they all possessed a cephalothoracic shield with frontal lunules, a leg 3 endopod bearing up to six setae, and a nauplius eye with small or laterally extending lenses but lacked dorsal plates or elongate posterior processes on the body somites. We provide the first molecular data for each of the following species.

#### *Caligus haemulonis* Krøyer, 1863

The four specimens collected from the gills of *M.* sebae – two females (Fig. 10A) and two males (Fig. 10B) – were identified as members of the *C. productus* group based on the reduction of the three plumose setae normally carried on the posterior margin of the distal exopodal segment of the first swimming leg (Fig. 10C) and a three-segmented leg 4 (i.e., two-segmented exopod) with four spines on the distal segment (Boxshall and El-Rashidy 2009).

**Figure 10.**
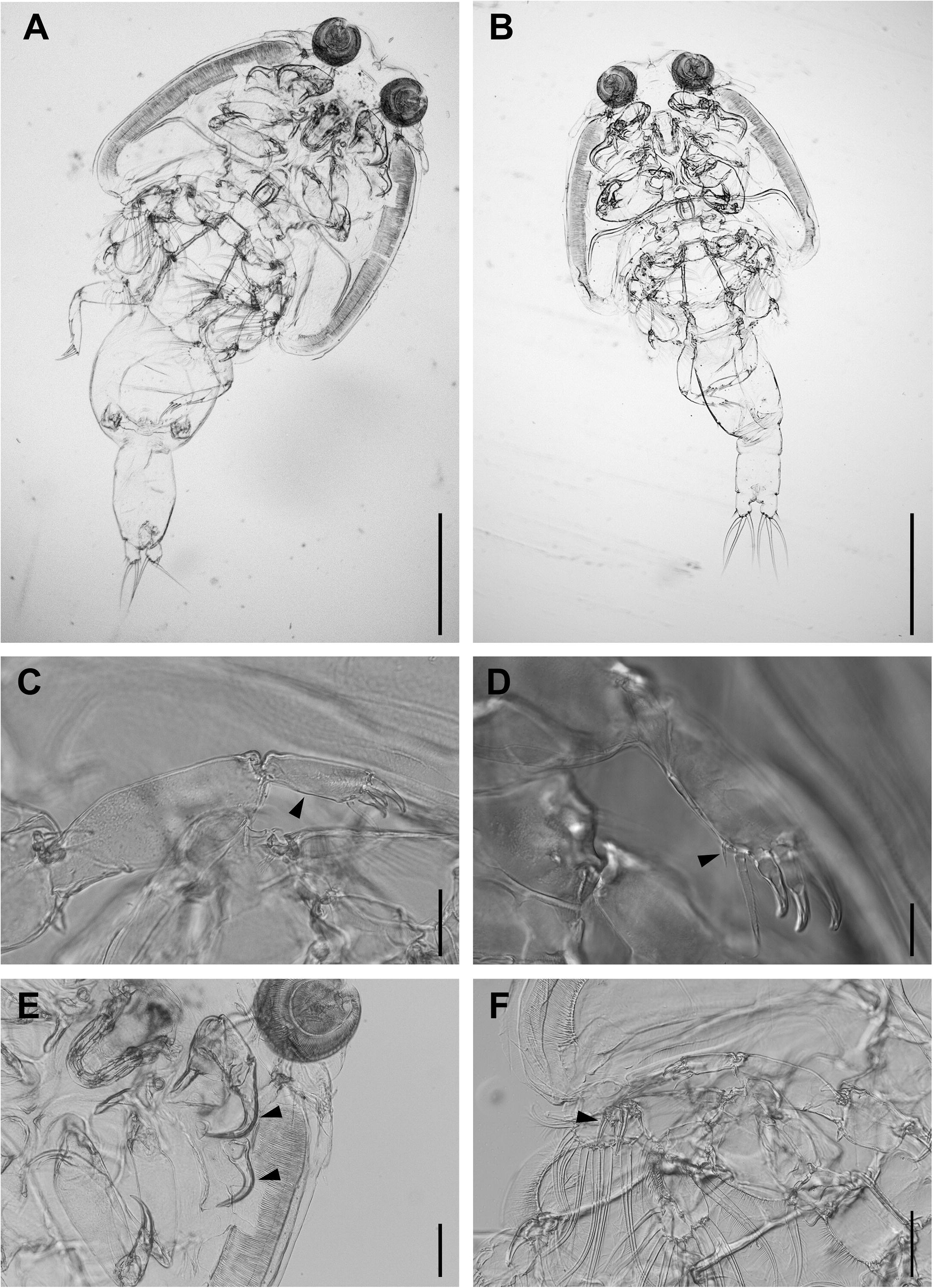
Microphotographs of *Caligus haemulonis* ex *Monodactylus sebae* using differential interference contrast. **A** Whole body of female and **B** male specimens. Morphological characters of female: **C** leg 1 without plumose setae carried on the posterior margin of the distal exopodal segment; **D** naked vestige of a seta of leg 1; **E** antennal claw and post-antennal process; and **F** spine on the leg 2 exopod segment 2. Scale bars: 500 µm (**A**, **B**); 50 µm (**C**); 20 µm (**D**); 100 µm (**E**, **F**).

Using a key for this species group (Boxshall and El-Rashidy 2009), the two female specimens most closely resembled *C. haemulonis* Krøyer, 1863, with the posterior margin of the distal exopodal segment of leg 1 possessing a naked vestige of a seta (Fig. 10D) (Boxshall 2018) and the genital complex and abdomen combined being about 1.3 times longer than the cephalothorax. Our specimens also resembled *C. schlegeli* Ho & Lin, 2003, which possesses similar characteristics, but lacks the strongly recurved shape of the antennal claw and post-antennal process (Fig. 10E) (Ho and Lin 2003; Boxshall 2018). Furthermore, the spine on the leg 2 exopod segment 2 is longer in *C. haemulonis* than in *C. schlegeli*. In *C. haemulonis* the spine extends to the margin of the distal segment (Cressey 1991), as seen in our specimens (Fig. 10F). In *C. schlegeli*, this spine has been described as not reaching the margin of the third segment (Ho and Lin 2003) or being much shorter than the third segment (Boxshall 2018). Therefore, we identified our specimens as *C. haemulonis*. Our specimens were smaller, i.e., 2.8 mm and 1.5 mm for the females, 2.8 mm and 2.5 mm for the males *vs.* 2.96–3.92 mm and 1.86–3.26 mm (Boxshall and El-Rashidy 2009), respectively, which may suggest that the female specimens were not yet mature.

We only obtained 28S sequences for *C. haemulonis*, with the closest matching reference sequence originating from a specimen assigned to *C. hoplognathi* Yamaguti & Yamasu, 1959 (98.3%), a member of the *C. bonito* species group (Boxshall 2018), collected in South Korea (Baek et al. 2016). However, the only sequences available for the *C*. *productus* group are 18S and *cox*1 sequences of *C. dakari* van Beneden, 1892 (Hayes et al. 2021). Therefore, using barcoding sequences to identify species of this group is currently severely limited. The sequences of all four specimens of *C. haemulonis* were identical.

*Caligus haemulonis* has previously been reported in West Africa, specifically off the Mauritanian coast (as *C. mauritanicus minuscula* Brian, 1924) on the gills of *Plectorhinchus mediterraneus* (Guichenot, 1850), *Carlarius heudeloti* (Valenciennes, 1840), *Sciaena umbra* L., 1758, and *Dentex* sp. (Brian 1924). The present record of *C. haemulonis* from *M. sebae* from Senegal, in the north-eastern Atlantic, represents an extension of the known geographical range of this species and constitutes a new host record. The inclusion of the monodactylid hosts raises the number of known host species from 30 to 31 (Boxshall and El-Rashidy 2009).

#### *Caligus* aff. *inopinatus*

One female specimen (Fig. 11A) infecting the gills of *S. afra* exhibited the characteristics of the *C. confusus* group including a raised cuticular rib and circular array of large denticles on the apron of leg 3 (Fig. 11B, i), a large recurved outer spine on the first exopodal segment of leg 3 (Fig. 11B, ii), a four-segmented leg 4 (i.e., three-segmented exopod) armed with I, I, II spines (Fig. 11A, i), and a postantennal process and maxillule posterior processes each bearing an accessory tine (Fig. 11C) (Boxshall 2018).

**Figure 11.**
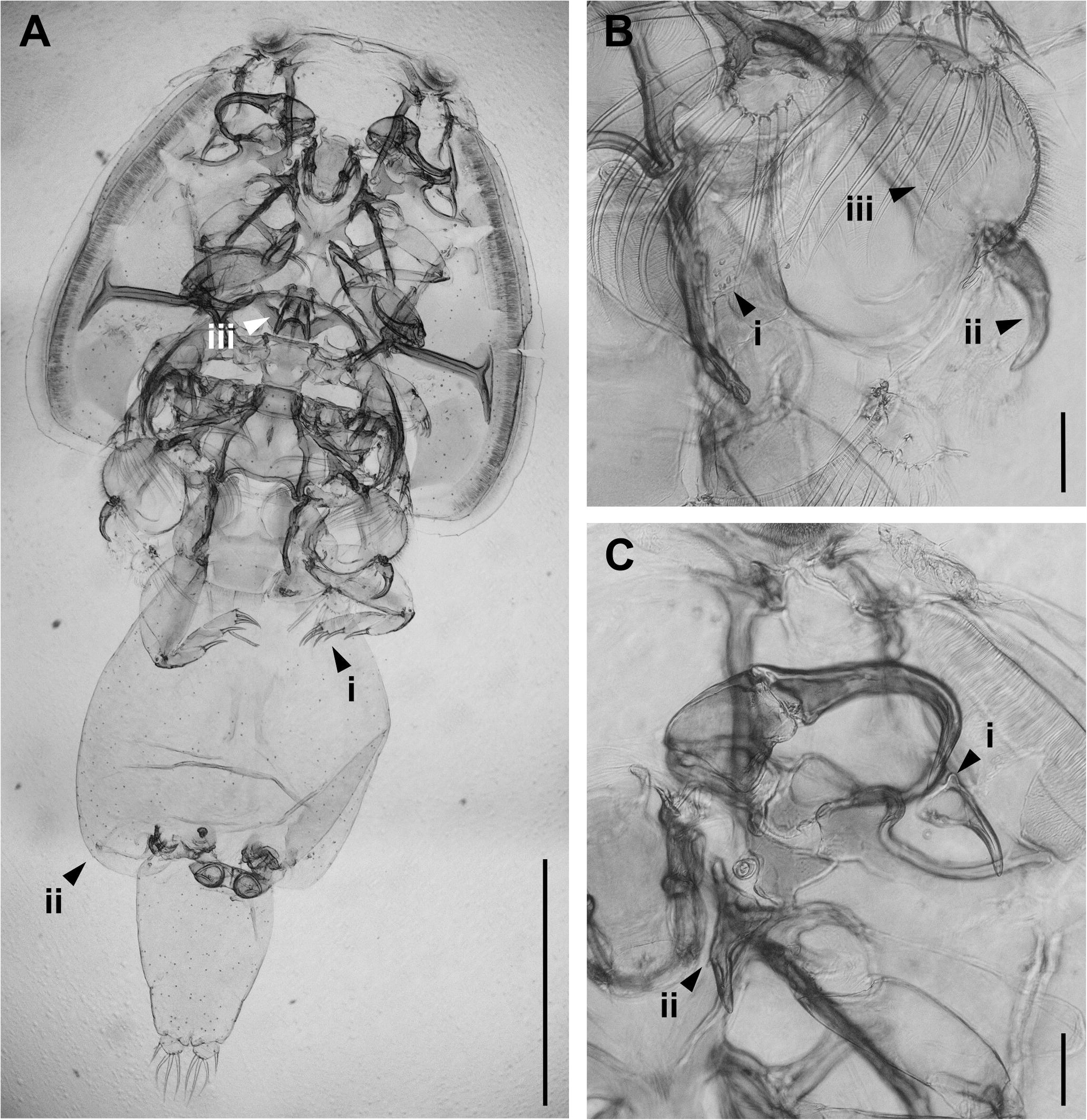
Microphotographs of female *Caligus* aff. *inopinatus* ex *Sphyraena afra* using differential interference contrast: **A** whole body with four-segmented leg 4 (i.e., three-segmented exopod) armed with I, I, II spines (i), an abdomen with rounded posterolateral corners (ii), and a sternal furca with widely set tines (iii); **B** Leg 3 with a raised cuticular rib and circular array of large denticles on apron (i), a large recurved outer spine on the first exopodal segment (ii), and a longitudinal row of denticles above the base of the exopod (iii); **C** antennae and maxillules, with a pronounced subconical swelling lateral the base of the postantennal process (i) and a maxillule posterior process with an accessory tine (ii). Scale bars: 1,000 µm (**A**); 100 µm (**B**, **C**).

Using a key for this species group (Özak et al. 2019), the specimen appeared most similar to *C. inopinatus* Kabata, 1994 and *C. abigailae* Boxshall, 2018, with an abdomen around 50% of the length of the genital complex (55% in our specimen) and rounded posterolateral corners (Fig. 11A, ii); in addition, both species infect barracudas (*Sphyraena* spp.) (Kabata 1994; Boxshall 2018). The relative length of the abdomen (1.6 times longer than wide) is more similar to *C. inopinatus*. Furthermore, our specimen presents characteristic features of *C. inopinatus* including a rounded post-antennal process with a pronounced subconical swelling lateral to its base (Fig. 11C, i) and a longitudinal row of denticles above the base of the exopod of leg 3 (Fig. 11B, iii) (Kabata 1994). However, compared to the original description of *C. inopinatus*, our specimen has a maxillule posterior process with an accessory tine (vs. single tine) (Fig. 11C, ii), caudal rami that are relatively much wider (length–width ratio 0.65 *vs.* 1.6), a much smaller genital complex–to–abdomen ratio (1.8 *vs.* 2.6), and a sternal furca with more widely set tines (Fig. 11A, iii). Furthermore, the present record is from the Tropical Atlantic whereas *C. inopinatus* has only been reported from the gills of *S. flavicauda* Rüppell, 1838 in the Red Sea (Kabata 1994). Therefore, we conclude that our specimen probably represents a new species resembling *C. inopinatus* and, therefore, we refer to it as *Caligus* aff. *inopinatus*.

We obtained 18S, 28S, and *cox*1 sequences, but only *cox*1 reference sequences were available for members of the *C. confusus* group, including *C. lichiae* Brian, 1906 (cited as *C. aesopus* Wilson, 1921), *C. confusus* Pillai, 1961, and *C. chorinemi* Krøyer, 1863 (Morales-Serna et al. 2014). Our *cox*1 sequence showed low similarity to these reference sequences (78%–79%).

For 18S, the most similar reference sequence was of a specimen assigned to *C. pelamydis* Krøyer, 1863 (Øines and Schram 2008), a member of the *C. diaphanus* species group (Boxshall 2018), collected in Norway from an unnamed host (Øines and Heuch 2005).

Therefore, as was the case for *C. haemulonis*, the number of available reference sequences for species of *Caligus* is currently insufficient for molecular identifications.

Caligidae gen. sp. ‘larva’

Two chalimus larvae, i.e. copepodids, were collected from a specimen of *E. fimbriata*. We recognised these specimens as caligids through the presence of a frontal filament, characteristic of larval caligids (Piasecki et al. 2023). The DNA sequences we obtained did not match closely (> 90%) with any of those currently available in GenBank or the adult specimens collected in the present study, but the sequences of the two specimens were identical. Therefore, we considered these specimens as an unidentified third species in our study and refer to these specimens as Caligidae gen. sp. ‘larva’.

### Nematoda Diesing, 1861

Chromadorea Inglis, 1983

Rhabditida Chitwood, 1933

Anisakidae Skrjabin & Karokhin, 1945

All nematodes collected in the present study (Fig. 12) were larval stages. Several specimens had a boring tooth at the anterior extremity (Fig. 12G–I), characteristic of the larval stages of nematodes belonging to Anisakidae (Weerasooriya et al. 1986). Seven specimens were tentatively placed in Anisakinae Railliet & Henry, 1912 based on the presence and position of the intestinal caecum and ventricular appendix (Fig. 12A, C, E) and the labial position of the excretory pore (Fig. 12G–I) (Anderson 2009; see following paragraphs). The species identification was based on the ITS1, ITS2, and *cox*1 barcodes.

**Figure 12.**
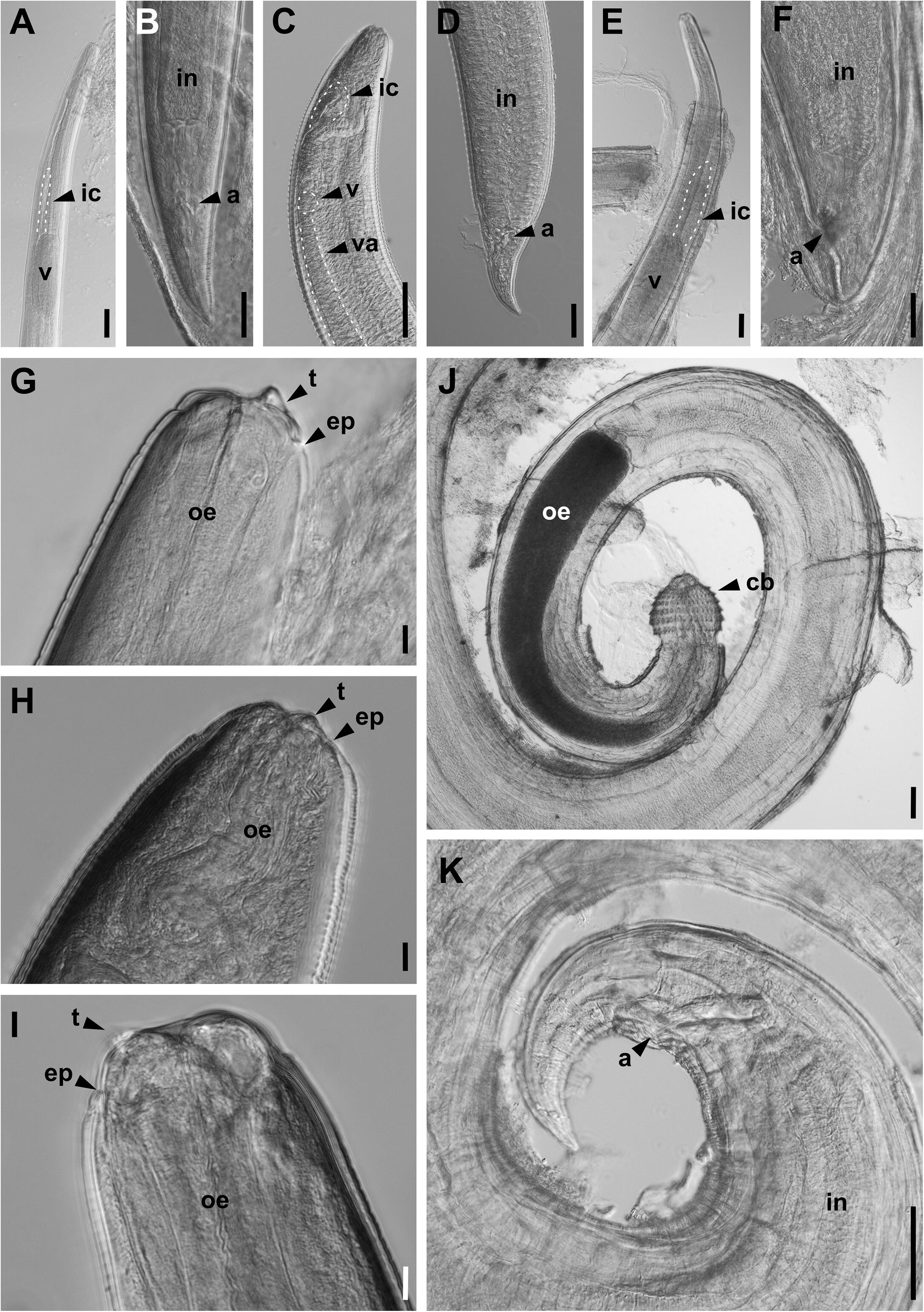
Microphotographs of larval nematodes using differential interference contrast. Anisakidae: *Euterranova pectinolabiata*, **A** anterior and **B** posterior ends; *Contracaecum rudolphii*, **C** anterior and **D** posterior ends; Anisakidae gen. sp. 1, **E** anterior and **F** posterior ends; cephalic region: **G** *E. pectinolabiata*; **H** *C. rudolphii*; and **I** Anisakidae gen. sp. 1. Gnathostomatidae: *Echinocephalus* sp. **J** anterior and **K** posterior ends. Labels: a – anus, cb – cephalic bulb, ep – excretory pore, ic – intestinal caecum, in – intestines, oe – oesophagus, t – boring tooth, v – ventriculus, va – ventricular appendix. Scale bars: 100 µm (**A**, **C**, **E**, **J**, **K**); 50 µm (**B**, **D**, **F**); 10 µm (**G**, **H**, **I**).

*Euterranova* Moravec & Justine, 2020

*Euterranova pectinolabiata* (Shamsi, Barton & Zhu, 2019)

Three specimens found in the mesentery of *C. hippos* and *I. africana* showed high levels of sequence identity to reference sequences assigned to *Euterranova pectinolabiata* (ITS1 and ITS2: 99.2–99.4% (Shamsi et al. 2019; Hossen et al. 2021)). These reference sequences encompassed those from the original description of the species from the elasmobranch, *Sphyrna mokarran* (Rüppell, 1837), off the coast of Australia (Shamsi et al. 2019) as well as larval specimens collected from species of *Platycephalus* Bloch,1795 off the coast of Australia (Hossen et al. 2021). Our specimens also possess a muscular oesophagus followed by an oval-shaped ventriculus that is less than half the length of the intestinal caecum (Fig. 12A) as well as an excretory pore at the base of the lips (Fig. 12G), as reported for larvae of

*E. pectinolabiata* (cited as *Terranova* larval type II (Cannon 1977; Hossen et al. 2021)).

Based on these morphological and molecular similarities, we considered our specimens conspecific with the Australian specimens. We provide the first published record of *cox*1 sequences for this species.

*Terranova*-type larval stages have been reported from Nigeria (Ogbeibu et al. 2014), but no species was assigned. Therefore, our records from *C. hippos* and *I. africana* constitute new host and geographical records, with elasmobranchs probably serving as the definitive hosts in the Saloum Delta.

*Contracaecum* Railliet & Henry, 1912

*Contracaecum rudolphii* Hartwich, 1964

The three specimens found in the mesentery of *E. fimbriata*, *G. nigri*, and *M. sebae*, respectively, showed high levels of sequence identity to specimens assigned to the *Contracaecum rudolphii* species complex, specifically the lineage referred to as *C. rudolphii* A (ITS1 and ITS2: 99.4–99.9%) collected from *Phalacrocorax carbo sinensis* (Staunton, 1796) in Israel (Caffara et al. 2023) and *Gulosus aristotelis* (L., 1761) in Italy (Farjallah et al. 2008). Both these records are accompanied by detailed morphological characterisations of the adult specimens in their definitive cormorant hosts. Therefore, we considered our specimens conspecific with *C. rudolphii* A. The only reference *cox*1 sequences of *C. rudolphii* A were amplified using the JB3/JB4.5 (= ASmit1/ASmit2) primer set (Lin et al. 2013), which does not overlap with the *cox*1 region targeted by the LCO1490/HCO2198 primer set (Folmer et al. 1994a). The latter region is frequently targeted for other nematode species using these or other primers (Prosser et al. 2013). This mismatch highlights the lack of a unified approach towards sequencing *cox*1 in this group of parasites (Prosser et al. 2013). Morphologically, our specimens possessed an excretory pore at the base of the lips (Fig. 12H) as well as a muscular oesophagus followed by a small ventriculus with an intestinal caecum and a ventricular appendix (Fig. 12C).

Larval nematodes belonging to *Contracaecum* have previously been reported from fishes in Senegal in the 1970s (Vassiliadès 1975) and other West African countries in the 1980s (Anyanwu 1983), but none of the records were identified to the species level. Therefore, our records from *E. fimbriata*, *G. nigri*, and *M. sebae* constitute new host and geographical records for *C. rudolphii* A, with the definitive hosts in the Saloum Delta probably being one of the cormorant species – *Microcarbo africanus* (Gmelin, 1789) or *P. c. lucidus* Lichtenstein, 1823 – occurring in the estuary (Borrow 2023).

#### Anisakidae gen. sp. 1

A single specimen found in the mesentery of *M. sebae* showed its highest sequence identity (28S: 99.5%, *cox*1: 95.4%) to a larval nematode collected off the Mexican coast from *Fundulus persimilis*, which was labelled as *Pseudoterranova* sp. (Espínola-Novelo et al. 2025). As the authors mentioned no morphological characters underpinning their identification, we consider this record unverifiable. The ITS and 18S sequences of our material have no close matches (> 95%) to any reference sequence in GenBank. Our specimen has an intestinal caecum and an oval-shaped ventriculus and no ventricular appendix (Fig. 12E) as well as an excretory pore at the base of the lips (Fig. 12I), reminiscent of the *Terranova* larval type II (see *E. pectinolabiata*). However, larvae of anisakid species belonging to *Euterranova*, *Neoterranova* Moravec & Justine, 2020, and *Pseudoterranova* Mozgovoi, 1951 all possess a similar intestinal and ventricular arrangement. As neither the DNA barcodes nor the morphological characteristics provide an unequivocal indication of the generic affiliation or species identity, we refer to this specimen as Anisakidae gen. sp. 1.

#### Gnathostomatidae Railliet, 1895

Echinocephalus Molin, 1858

*Echinocephalus* sp.

Several nematode specimens with a cephalic bulb armed with rows of hooks at their anterior end (Fig. 12J–K; Fig. 13) were found encapsulated mostly within stomach walls and connective tissues of *G. nigri*, but one specimen was also found in the intestine of *C. guineensis*. The only group of nematodes with this characteristic known to infect fishes are species of *Echinocephalus* (Anderson 2009). However, larval stages of *Echinocephalus* cannot be definitively identified to the species level based on current knowledge, as their reproductive organs and number of cephalic hooks are not fully developed (Moravec and Justine 2021; Saad et al. 2022).

**Figure 13.**
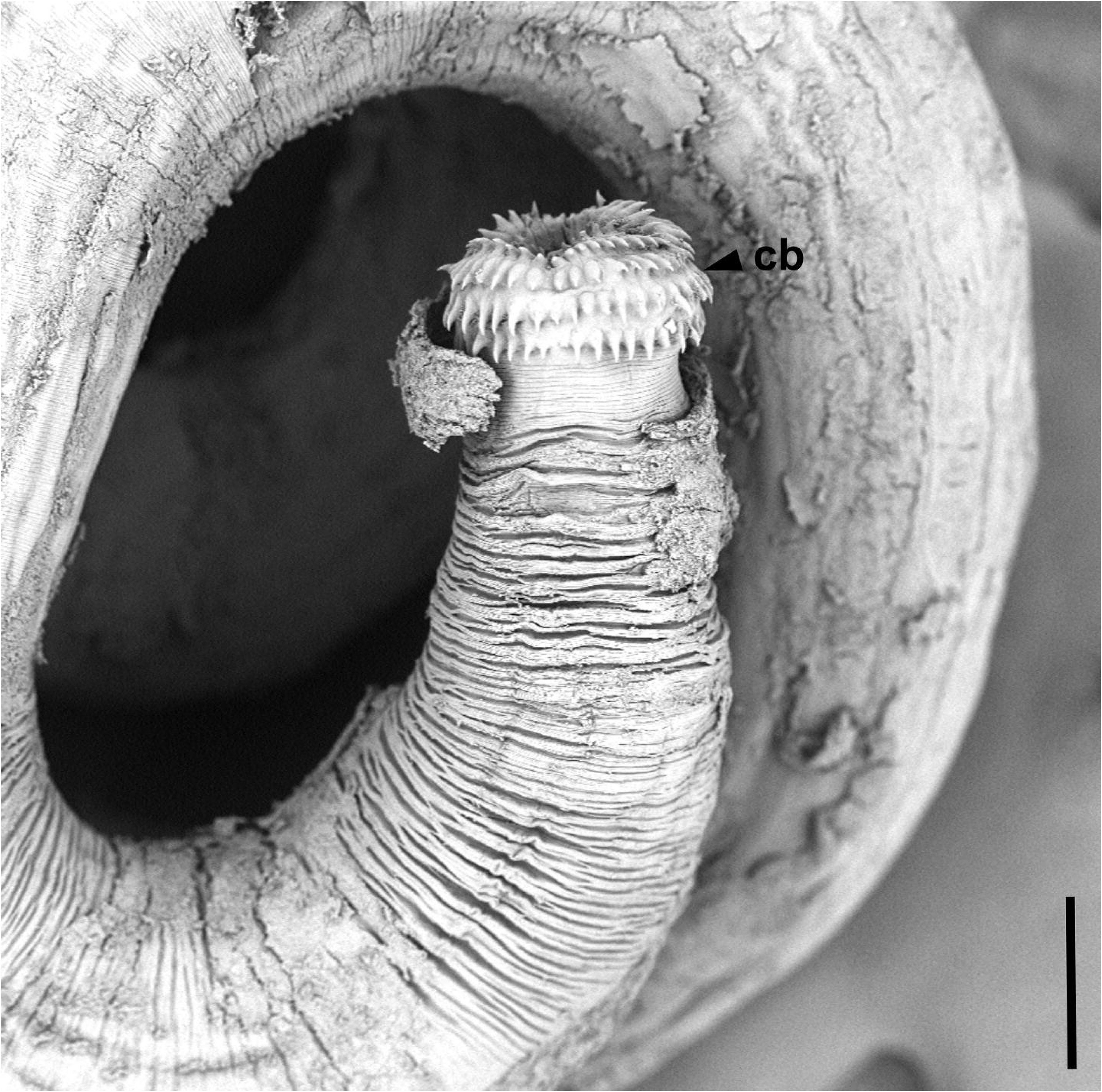
Scanning electron microscope image of *Echinocephalus* sp. with cephalic bulb (cb) armed with hooks.

Only 18S and *cox*1 reference sequences of species of *Echinocephalus* were available on GenBank. Our 18S sequences were highly similar (99.9–100%) to a sequence of *E. overstreeti* Deardorff & Ko, 1983 found on *Heterodontus portusjacksoni* (Meyer, 1793) off the Australian coast (Laetsch et al. 2012), although 18S may be too conserved to identify nematode species (Chan et al. 2021). Our *cox*1 sequence matched most closely with sequences assigned to *E. spinosus* Müller, Adriano, Oliveira & Corrêa, 2022 (81.0–86.4%) collected off the Brazilian coast from *Potamotrygon motoro* (Müller & Henle, 1941) (Müller et al. 2022). Furthermore, no reference sequences were available for the 28S regions. As no species name could be assigned morphologically or molecularly, we refer to our specimens as *Echinocephalus* sp.

In West Africa, only larval specimens of *Echinocephalus* have previously been reported, with the hosts being *M. sebae*, *Pseudotolithus senegalensis* (Valenciennes, 1833), and *P. elongatus* (Bowdich, 1825) in Nigeria (Ogbeibu et al. 2014). The present specimens from *G. nigri* and *C. guineensis* in Senegal, therefore, constitute new host and geographical records for this group of nematodes, with elasmobranchs probably being the definitive hosts in the Saloum Delta, similar to other species of this group (Moravec and Justine 2021). Here, we provide the first 28S sequences for *Echinocephalus*.

### Species richness: marine protected area vs. open-access area

Richness analyses were restricted to *C. guineensis*, *G. nigri*, *M. sebae*, and *S. melanotheron* as these were the only host species sampled in sufficient numbers to draw comparisons between the MPA and OAA. We included only host species with multiple corresponding parasite species as single observations could not mathematically yield biologically meaningful richness estimates through extrapolation, as implemented in iNEXT. However, we emphasise that singletons (i.e., unique observations) of parasite species were included for the four host species.

(i) Infracommunities: generalised linear models

In the GLM analysis of the Shannon and Simpson indices of the infracommunities, all interaction effects were unsupported, and therefore dropped. Only the host species ID remained as a significant effect in the models estimating the Shannon Index for the gill and combined datasets (Table 6). Similarly, only the standard length of the host remained as a significant effect on the Simpson index for the gill parasites. For the combined dataset (guts and gills), site (MPA vs. OAA) and host standard length had significant effects on the Simpson index. The odds of hosts harbouring infracommunities with a higher Simpson index values (lower diversity) were estimated to be 4.95 times higher in the MPA than in the OAA (95% CI: 0.08, 0.49). This suggests that infracommunities were estimated to be less species-rich in the MPA (Fig. 14). Furthermore, for every 1 mm increase in host standard length, the odds of observing a higher Simpson index decreased by 22.3% (Odds Ratio = 0.78, 95% CI: 0.61, 0.98), indicating that larger hosts harboured more species-rich parasite communities.

**Figure 14.**
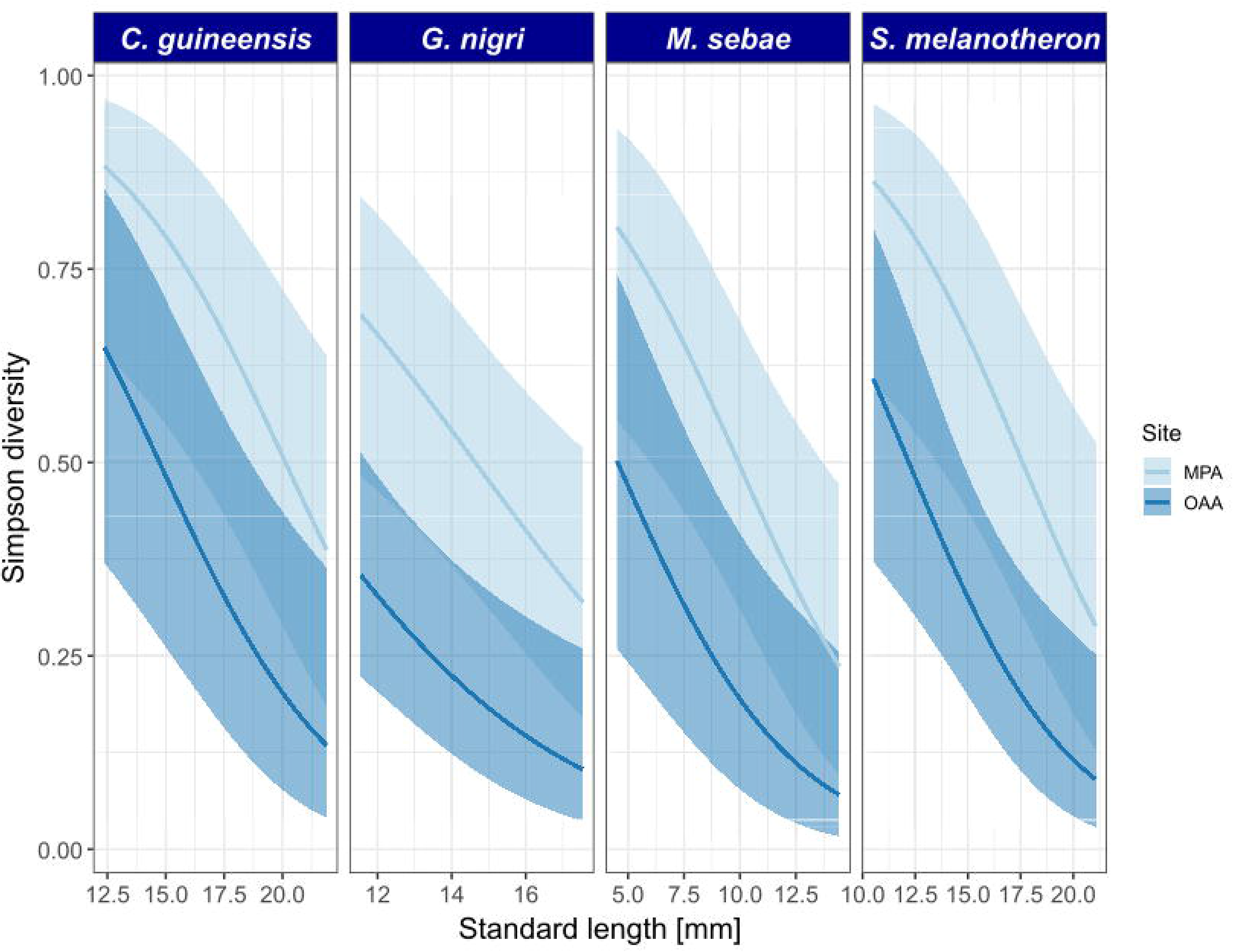
Predicted Simpson biodiversity for combined parasite communities (gills and guts) in the Bamboung Marine Protected Area (MPA) and the Bolong de Sangako Open-Access Area (OAA) as a function of standard lengths of fish hosts, according to the optimised generalised linear model; shaded areas indicate confidence intervals. Richness analyses were restricted to *C. guineensis*, *G. nigri*, *M. sebae*, and *S. melanotheron* to exclude host species with single observations, as these do not yield biologically meaningful richness estimates. Biodiversity was estimated to be significantly higher outside the MPA for all four host species, but the overlapping confidence intervals indicate that pairwise differences were not significant.

Whilst the effects of site and host size were significant overall, the predicted values of the indices overlapped at the 95% confidence level (Fig. 14), suggesting that the differences could not be resolved at the species-specific level given the present data.

(ii) Component communities: biodiversity indices

The difference in species diversity of parasite component communities varied by host species and dataset, i.e., gills *vs.* gills and guts (Figure 15). When estimating biodiversity indices for the gill parasites through individual-based extrapolation, component communities in the MPA presented lower values for extrapolated Shannon/Gini-Simpson diversity and species richness than in the OAA for *C. guineensis* and *M. sebae*. The reverse trend was observed in the parasite component communities infecting *S. melanotheron*.

**Figure 15.**
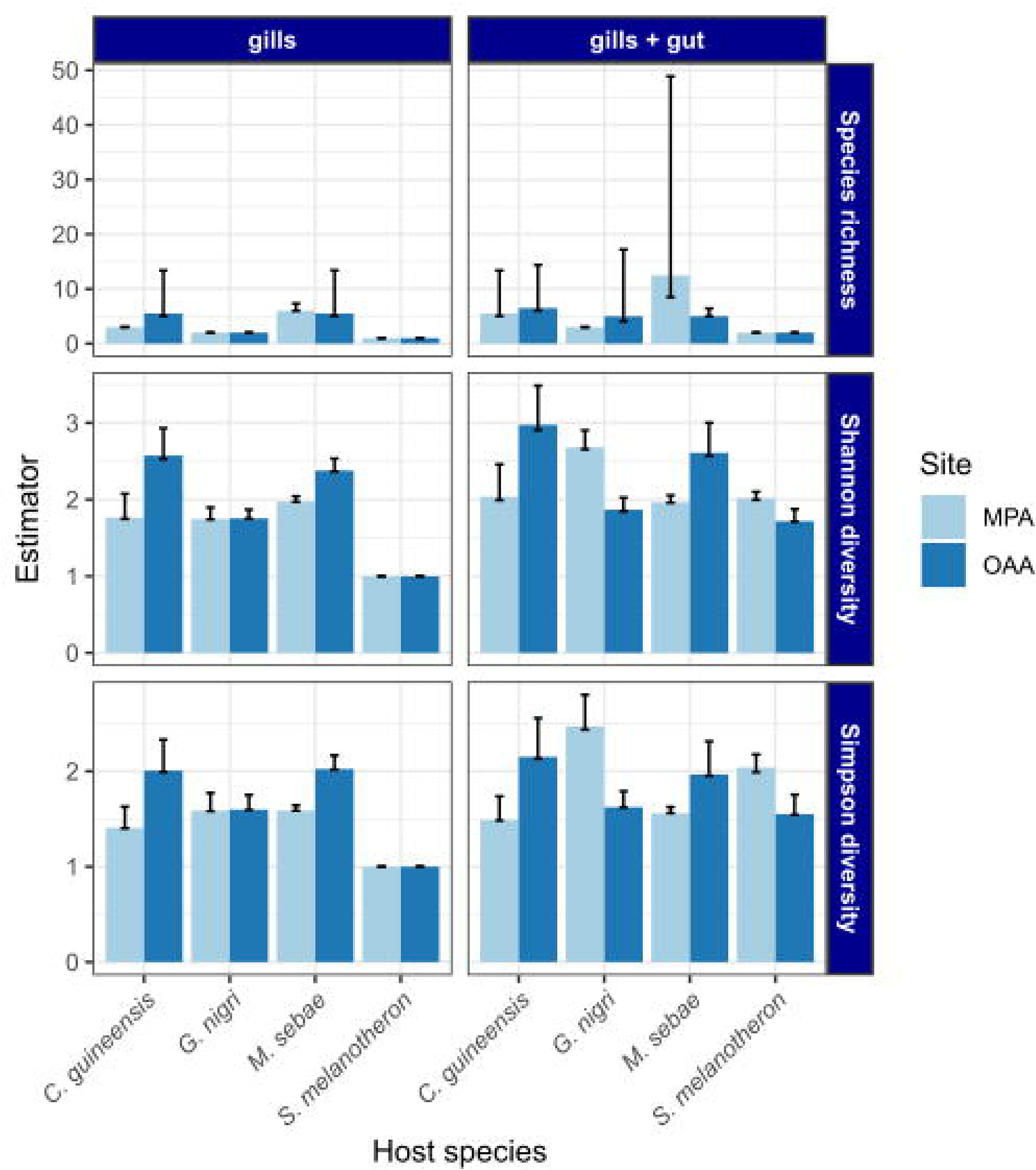
Estimated species diversity of component communities in the Bamboung Marine Protected Area (MPA) and the Bolong de Sangako Open-Access Area (OAA) based on rarefaction by parasite individuals and abundance data. Richness analyses were restricted to *C. guineensis*, *G. nigri*, *M. sebae*, and *S. melanotheron*, as all other host species were infected by only a single macroparasite species and therefore did not yield biologically meaningful richness estimates.

As gut parasites were observed in lower numbers than the gill parasites or even as singletons, biodiversity indices calculated based on these observations were not informative. However, the analysis of the combined data set (gut and gill parasites) indicated significant differences in the estimated biodiversity. For *C. guineensis*, the differences between MPA and OAA were smaller for the combined dataset than for the gill dataset. For the component communities infecting *G*. *nigri*, *M. sebae*, and *S. melanotheron*, Shannon and Simpson biodiversity were higher in the MPA than in the OAA.

The results of the additional analysis mostly conformed to the present results. However, the errors in the biodiversity indices through sampling unit–based extrapolation were markedly larger, which rendered some differences non-significant (Supplementary File S2). Host-parasite interaction diversity was lower in the MPA than in the OAA for the gill parasites, but no difference was detected for the gut and the combined dataset (Supplementary File S2).

## Discussion

### A new baseline for marine parasite diversity in West Africa

We present the first community-level survey of macroparasites in and around a West African MPA. In total, 5,020 parasite specimens belonging to 27 species were collected from 107 host specimens representing ten species. This is also the first study in West African wild fishes to simultaneously identify all major macroparasite groups at the species level. While previous research in West African wild fishes provided valuable taxonomic baselines, these efforts were typically restricted to individual taxonomic groups such as monopisthocotylans (Dossou 1982; Pariselle and Euzet 2004; Diamanka et al. 2011; Domingues et al. 2011; Boeger et al. 2012), polyopisthocotylans (Euzet and Wahl 1970), copepods (Brian 1924), acanthocephalans (Troncy and Vassiliadès 1974; Matouke et al. 2011), cestodes (Scholz et al. 2018a), and nematodes (Anyanwu 1983; Obiekezie et al. 1992), or identified specimens at higher taxonomic levels (Obiekezie et al. 1992; Palm et al. 1994; Boufahja et al. 2015).

Out of the 27 species, 15 (56%) represent first records for Senegal and 11 (41%) for West Africa. Globally, 21 out of 33 (64%) host-parasite combinations reported here constitute new host records. These new records improve the understanding of the geographical distribution and host repertoires of these parasites. The number of studies on marine parasites is particularly low along the coast of sub-Saharan Africa (Boufahja et al. 2015; Reed 2015). For instance, the last taxonomic study using newly collected material of marine trematodes in West Africa dates back 45 years (Szuks 1981) and that of marine caligid copepods 102 years (Brian 1924). Here, we provide the first record of *Pseudorhabdosynochus hargisi* since 1984 (Aljoshkina 1984). We provide only the second record of species of *Echinocephalus* from West Africa (Obiekezie et al. 1992). Furthermore, we also found three undescribed species of *Euryhaliotrema*. These examples illustrate the substantial knowledge gaps concerning the species diversity, host repertoire, and geographical distribution of parasites of marine fishes in the region.

The novelty of this study also lies in the potential of trophically transmitted parasites to act as biological tracers of food webs [see also Knudsen et al. (2014) and Pitt et al. (2026)]. Trophic transmission is supported here by larval nematodes (*Contracaecum rudolphii* and *Euterranova pectinolabiata*) that link infected prey fishes to bird and elasmobranch predators (Shamsi et al. 2019; Mattiucci et al. 2020), as well as trematodes closely related to the parasite of sea snakes (Martin et al. 2023), *Tubulovesicula laticaudi*. While the complete life cycles of these species in the Saloum Delta have yet to be fully characterised, our records provide a baseline to link these larval stages to their definitive hosts. Consequently, these findings reinforce the role of multi-host parasites as key indicators and tracers of marine ecosystem connectivity.

### A new molecular reference database for parasites of mangrove fishes

We provide 208 DNA barcodes covering 26 species identified in the present study, all of which represent the first molecular data for these species in West Africa. For eight (31%) of these species, these are the first molecular data worldwide; for 14 (54%), we provide the first sequences for at least one DNA region. The new sequences alleviate the issue of missing molecular data in those 26 parasite species for the most widely used barcoding markers.

However, we suggest that the broader value of these data lies in matching sequencing records in future studies as a first reference library for macroparasites in the region. Specifically, our data could inform studies using high-throughput sequencing to study parasite communities from tissue or gut content homogenates (Owens et al. 2024; Rijal et al. 2024) or environmental samples (Thomas et al. 2022; Hill-Spanik et al. 2025). These assays often target regions that are also used for standard barcoding including the nuclear ribosomal and mitochondrial DNA regions (Chan et al. 2021). Our DNA sequences may hence help build reference libraries for future metabarcoding studies. Ultimately, our data may support efforts to use high-throughput sequencing to monitor parasite communities in West Africa.

### A study system to compare species diversity of marine parasites

Beyond the novelty of species records and molecular data, we empirically estimated the species diversity of the parasite communities. We detected significant variations in biodiversity indices between a protected (Bamboung MPA) and an open-access (Bolon de Sangako OAA) mangrove ecosystems at the level of intra-host communities (infracommunities) and site-specific communities (component communities). Similar variations were also observed when diversity indices were applied to observed host-parasite interactions rather than individual parasite counts. Overall, these analyses suggested that parasite species richness on fish gills was higher in the OAA across three of the four host species (*C. guineensis*, *G. nigri*, and *M. sebae*) (Fig. 14–15; Supplementary File S2).

However, when combining gill and gut parasite data, these differences were diminished or reversed, though this reversal occurred exclusively at the component community level (Fig. 15). The patterns observed for the gill parasite communities may stem from the low sensitivity of parasites with single-host life cycles, including monopisthocotylan species, to environmental disturbance (Wood and Lafferty 2015; Sures et al. 2017; Wood et al. 2023). As fisheries in the Saloum Delta (Sadio et al. 2015; Thiaw et al. 2021) and elsewhere (Christensen et al. 2014) often target large piscivores, the smaller arthropodivores and omnivores evaluated here may experience predation release and competitive release, essentially leading to host population expansions, which could benefit monopisthocotylan parasites. Conversely, parasite species with complex life cycles involving more than one host species are probably more vulnerable to fisheries pressure (Wood et al. 2014) because fisheries can deplete the intermediate, paratenic, or definitive hosts required for completing their life cycles. This might explain why parasite species richness across gill and gut was higher in the MPA than the OAA for some host species.

However, we caution against extrapolating these trends given the small host sample size and single time point. Instead, we propose that our results demonstrate the effectiveness of the proposed study design (MPA *vs.* OAA) but also indicate limitations for testing the impact of fisheries on parasite biodiversity. If the sampling is repeated over multiple years, patterns in parasite species diversity shaped by the protection of fish assemblages in the MPA may be detected more reliably. For instance, one may explicitly test the impact of life cycle complexity in future studies. Our present data can support such approaches by enabling researchers parameterise a power analysis to determine the number of fishes required to detect differences between the MPA and OAA. Testing other similar sites with this study design – MPA *vs.* OAA – akin to studies on fish communities (Sadio et al. 2015; Thiaw et al. 2021), may also enable inferring more general trends concerning the impact of MPAs on macroparasite biodiversity.

Beyond the single time point, the present data also illustrate other limitations. Effectively studying the diversity of gut parasites may require a more substantial sampling effort than for the gill parasites. Specifically, monopisthocotylans – as the main group of parasites we collected from the gills – were substantially more abundant than any other parasite group.

This numeric imbalance has previously been reported for parasites of fishes, particularly at lower latitudes, as ectoparasite abundance appears to increase while endoparasite abundance remains relatively stagnant with declining latitude (Rohde and Heap 1998). The numeric dominance of monopisthocotylans may be promising for the use of these organisms as indicators of anthropogenic impact (Dzikowski et al. 2003; Kasembele et al. 2026).

Taxonomic resolution for these organisms may be high, as monopisthocotylans can typically be identified to the species level, unlike other taxa where larval stages lack fully developed morphological characters (see nematodes and trematodes in ‘Results’). Furthermore, the high number of monopisthocotylans may provide an ethical advantage as the number of host specimens that need to be sacrificed to infer statistically significant trends is reduced.

However, the imbalance between monopisthocotylans and other parasite taxa also creates a trade-off: the more time spent identifying numerous monopisthocotylans, the less time is available to screen enough hosts to detect statistically significant trends for other parasite groups.

Another issue concerns the treatment of species names in biodiversity analyses. New species may regularly be encountered when screening host specimens in understudied regions like West Africa (Jorge and Poulin 2018; e.g. Fig. 5). New species necessitate detailed morphological analysis to delimitate them from previously known closely related species through formal species descriptions. However, the practical need to publish ecological findings means researchers cannot always wait for the formal description of every new taxon, a process further slowed by a lack of specialists (Poulin and Presswell 2022) and the time-intensive nature of taxonomic work (Bouchet et al. 2023). In some studies, parasite specimens may not be identified to the species level at all (Poulin and Leung 2010; Poulin 2019; Hammoud et al. 2026). While this approach still allows for meaningful quantitative comparisons of community-wide trends when formal identification is not immediately possible, it may mask species-specific responses or render the biological interpretation of patterns challenging.

Even if specimens are identified to the species level, community ecological studies often deposit no photographic or physical vouchers in natural history collections (Turney et al. 2015), rendering these studies non-verifiable. Repeatability matters, as parasites can be particularly challenging to identify due to their often small body sizes and scarcity of known phenotypic characters to discriminate species reliably (Perkins et al. 2011; Thaenkham et al. 2022). The absence of digital or physical vouchers makes confirming records impossible, a situation that may result in incorrectly named records remaining indefinitely.

In the present study, we opted to provide a detailed justification of all species identifications verified by taxon-specific experts (part of the author list). Photographic evidence was submitted to a public repository (*iNaturalist*), and morphological voucher specimens were submitted to curated natural history collections (HU, MNHN Paris and Rabat). The approach detailed here may present a blueprint of how to survey parasites of fishes not only in West Africa, but also other underexplored regions.

### Concluding remarks

The relevance of the present survey is apparent for parasitological research in Senegal: we updated the list of species occurrences in the Bamboung MPA and the Saloum Delta as a whole. But how relevant are such species surveys in the wider context of parasite ecology?

We conclude that studies like ours establish a critical baseline for aquatic parasite biodiversity in historically understudied regions. Situated in the Tropical Atlantic yet adjacent to the Temperate Northern Atlantic, Senegal represents a biogeographic transition zone where marine species assemblages overlap with the Mediterranean and the Gulf of Guinea (Polidoro et al. 2017; Caballero Herrera et al. 2022). By documenting species that are globally distributed but locally unrecorded, this research partially eliminates an important geographical ‘blind spot’ and provides distribution data with global relevance.

Beyond documenting diversity, these surveys also establish a key baseline for community ecology. By integrating macroparasites – a group that rivals free-living taxa in terms of species richness and biomass (Dobson et al. 2008; Lambden and Johnson 2013; Preston et al. 2013) – we account for a frequently overlooked component of ecosystem complexity. As the inclusion of these organisms substantially restructures known food web topologies (Lafferty et al. 2006; Morton and Lafferty 2022), a more robust characterisation of parasite assemblages is essential for accurately quantifying how anthropogenic pressures influence coastal ecosystems.

## Acknowledgements

We thank Ansou Mané for technical support during field work in the Saloum Delta. Natascha Steffanie is thanked for supporting the molecular work. Fons Verheyde and Chilekwa Chisala from the Flanders Marine Institute (VLIZ) Library are thanked for providing access to part of the taxonomic literature. Rodney A. Bray and Geoffrey A. Boxshall (both NHM London) are thanked for their feedback concerning some of the species identifications. The *Direction des Aires Marines Communautaires Protegées* (DAMCP) of the Senegalese Ministry of the Environment and Sustainable Development (MEDD) is thanked for facilitating and authorising sampling in the Saloum Delta (0000494 MEDD/DAMCP). The Direction des Parcs Nationaux (DPN) of MEDD is thanked for authorising sample export under the Nagoya Protocol on Access and Benefit Sharing (001355 DPN/PF-APA).

## Data resources

The data supporting the findings of this study are openly available in public repositories. DNA sequence data generated during this study have been deposited in NCBI GenBank (see Table 5 for accession numbers). High-resolution specimen images and associated morphological data are archived on MorphoBank under Project ID XXXXXX (https://morphobank.org/XXXXXX), and images are also accessible on iNaturalist (project ID: 300535, URL: https://www.inaturalist.org/projects/300535). All R scripts used for statistical and computational analyses are available on GitHub (https://github.com/HU-Parasitology-and-OneHealth/Baseline-AquaticParasites-Mangroves-Saloum) and have been permanently archived on Zenodo (https://doi.org/XXXXXX).

## Supplementary Files

**Supplementary Table S1.** Polymerase chain reaction (PCR) primers, targeting barcoding regions, and thermal cycling profiles utilised in this study. For multi-primer assays, individual primer volumes, and specific stock conditions are annotated in the footnotes.

**Supplementary File S2.** Results of additional biodiversity analyses

**Supplementary File S3.** Data of invidiual host and parasite specimens collected in the Saloum Delta, Senegal.

