## Supplementary Table S1 for "A baseline for aquatic parasite diversity in West African mangroves: host-parasite records, DNA barcodes, and an experimental design for future use"

**Supplementary Table S1.** Polymerase chain reaction (PCR) primers, targeting barcoding regions, and thermal cycling profiles utilised in this study. For multi-primer assays, individual primer volumes, and specific stock conditions are annotated in the footnotes.

| Barcoding region | Forward primer(s) | Reverse primer(s) | Primer V [µl] | Intital denaturation | | Denaturation | | Annealing | | Extension | | Number of cycles | Final extension | |
|---|---|---|---|---|---|---|---|---|---|---|---|---|---|---|
| | | | | T [°C] | t | T [°C] | t | T [°C] | t | T [°C] | t | | T [°C] | t |
| 18S | WormA | WormB | 1.25 | 95 | 3 min | 95 | 20 s | 50 | 50 s | 72 | 3 min | 35 | 72 | 5 min |
| | Nem_18SF | Nem_18SR | 1.25 | 94 | 5 min | 94 | 30 s | 54 | 30 s | 72 | 60 s | 37 | 72 | 10 min |
| | Aca-SSUF | Aca-SSUR | 1.25 | 94 | 3 min | 94 | 60 s | 58 | 60 s | 72 | 60 s | 35 | 72 | 7 min |
| | 18SF | 18SR | 1.25 | 94 | 4 min | 94 | 30 s | 54 | 30 s | 72 | 60 s | 35 | 72 | 7 min |
| 18S + ITS1 | S1 | IR8 | 1.25 | 94 | 2 min | 94 | 60 s | 53 | 60 s | 72 | 90 s | 39 | 72 | 10 min |
| | S1 | Lig5.8R | 1.25 | 95 | 2 min | 95 | 60 s | 55 | 60 s | 72 | 90 s | 39 | 72 | 10 min |
| ITS1 + ITS2 | NC5 | NC2 | 1.25 | 94 | 5 min | 94 | 30 s | 54 | 30 s | 72 | 60 s | 37 | 72 | 10 min |
| ITS2 | 3S[1] | ITS2.2[1] | 1.00 | – | – | 95 | 3 min | 45 | 2 min | 72 | 90 s | 1 | – | – |
| | | | | – | – | 95 | 45 s | 50 | 45 s | 72 | 90 s | 4 | – | – |
| | | | | – | – | 95 | 20 s | 52 | 20 s | 72 | 90 s | 30 | 72 | 5 min |
| 28S | C1 | D2 | 1.25 | 94 | 2 min | 94 | 20 s | 56 | 30 s | 72 | 90 s | 39 | 72 | 10 min |
| | CichF28Sl | CichR28Sl | 1.25 | 94 | 2 min | 94 | 60 s | 53 | 60 s | 72 | 90 s | 39 | 72 | 10 min |
| | LSU5 | 1500R | 1.25 | 95 | 3 min | 95 | 50 s | 58 | 50 s | 72 | 80 s | 35 | 72 | 4 min |
| | NEM28SF | NEM28SR | 1.25 | 94 | 5 min | 94 | 30 s | 54 | 30 s | 72 | 60 s | 37 | 72 | 10 min |
| | 28SF | 28SR | 1.25 | 94 | 4 min | 94 | 30 s | 54 | 30 s | 72 | 60 s | 35 | 72 | 7 min |
| *cox*1 | ASmit1[2] | Cox1_Schisto_3<br>ASmit2[2] | 1.00 | 95 | 5 min | 95 | 60 s | 45 | 60 s | 72 | 60 s | 40 | 72 | 7 min |
| | | | 1.00 | 95 | 5 min | 95 | 60 s | 50 | 60 s | 72 | 60 s | 40 | 72 | 7 min |
| | Dig-cox1Fa | Dig-cox1R | 2.5 | 94 | 3 min | 94 | 30 s | 50 | 30 s | 72 | 30 s | 40 | 72 | 10 min |
| | LCO1490 | HCO2198 | 1.25 | 95 | 5 min | 95 | 30 s | 47 | 30 s | 72 | 60 s | 37 | 72 | 7 min |
| | NemF1_t1,<br>NemF2_t1,<br>NemF3_t1[1] | NemR1_t1,<br>NemR2_t1,<br>NemR3_t1[1] | 1.25[3] | 94 | 60 s | 94 | 40 s | 45 | 40 s | 72 | 60 s | 5 | – | – |
| | | | | – | – | 94 | 40 s | 51 | 40 s | 72 | 60 s | 35 | 72 | 5 min |

[1] PCR with varying conditions per cycle. [2] Nested PCR: product of 1st PCR was diluted 1:100 and used as template for second PCR. [3] Primer mix: 1.25 µl each of a 10-µM forward primer mixture and a 10-µM reverse primer mixture added.
