## Supplementary File S2 for "A baseline for aquatic parasite diversity in West African mangroves: host-parasite records, DNA barcodes, and an experimental design for future use"

**Supplementary File S2.** Results of additional biodiversity analyses

*Component communities: sampling unit–based rarefaction*

The results of this analysis (Fig. S2.1) mostly conformed to the individual-based rarefaction analysis (compare Fig. 11). However, the errors in the estimated biodiversity indices are markedly larger, which renders some differences non-significant. As observed in the individual-based analysis, the extrapolated species diversity of component communities of gill parasites for *C. guineensis* and *G. nigri* was significantly lower in the MPA than the OAA, but higher in the MPA for the communities infecting *S. melanotheron*. The inclusion of the gut parasite data rendered this difference for *C. guineensis* and *G. nigri* non-significant, but the parasite species diversity for *S. melanotheron* remained higher in the MPA than the OAA.

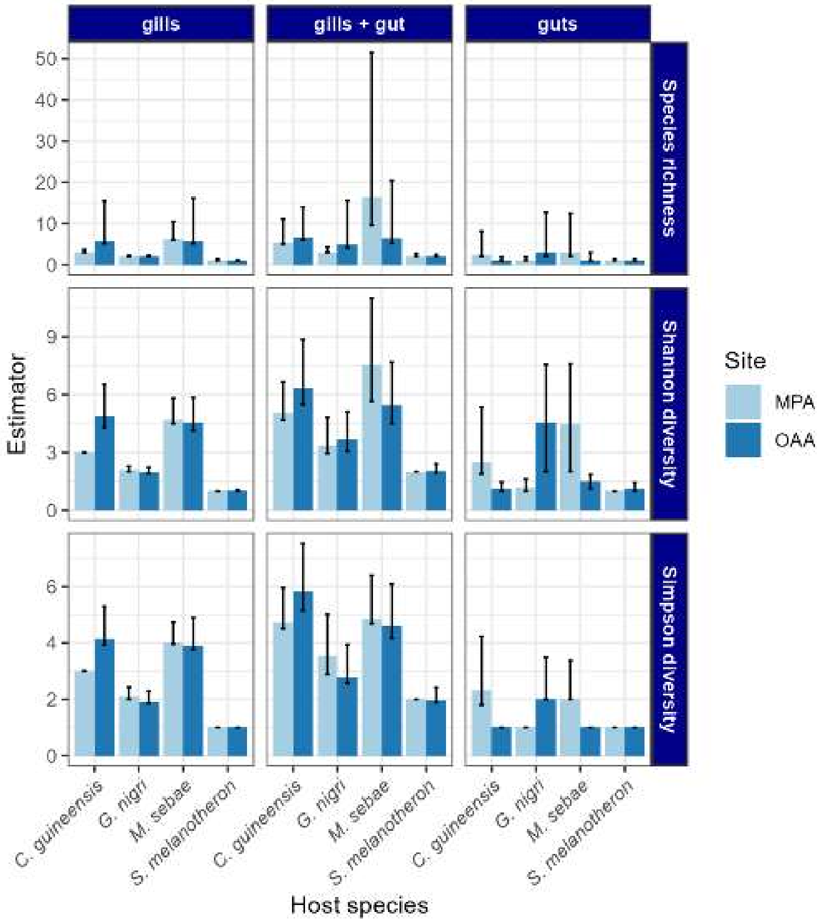


**Figure S2.1**. Estimated species diversity of component communities in the Bamboung Marine Protected Area (MPA) and the Bolong de Sangako Open-Access Area (OAA) based on rarefaction by sampling units and incidence data (presence/absence per host individual). Richness analyses were restricted to *C. guineensis*, *G. nigri*, *M. sebae*, and *S. melanotheron*, as all other host species were infected by only a single macroparasite species and therefore did not yield biologically meaningful richness estimates.

*Interaction diversity of species networks*

Similar to the results on the parasite biodiversity, the extrapolated Shannon and Gini-Simpson interaction diversity was estimated to be significantly lower in the MPA than in the OAA for the gill parasite communities (Fig. S2.2). Species richness estimates for all three datasets did not differ significantly between the MPA and OAA for all the gut parasites and the combined dataset (gut and gill parasites).

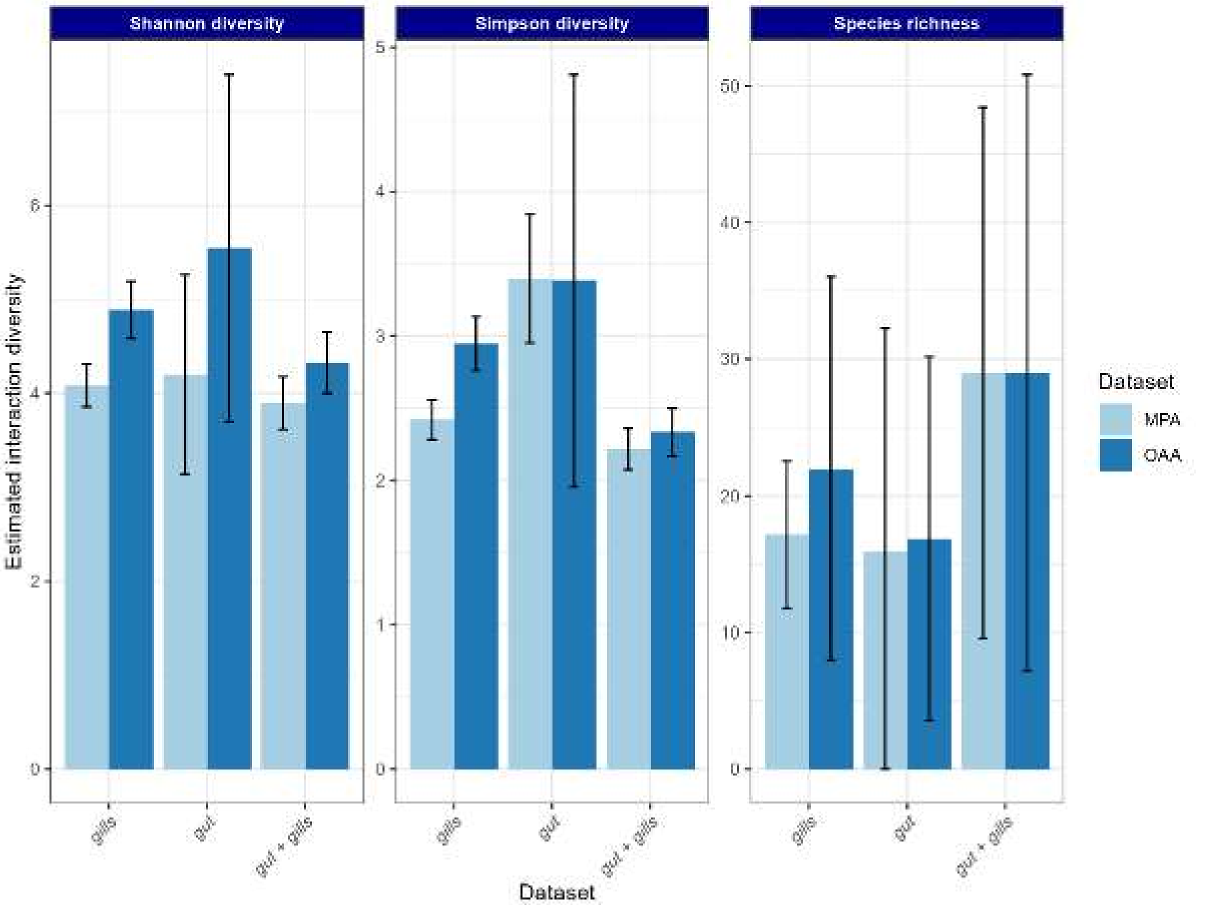


**Fig. S2.2.** Estimated interaction diversity of host-parasite networks in the Bamboung Marine Protected Area (MPA) and the Bolong de Sangako Open-Access Area (OAA) based on rarefaction of observed pairwise interactions of host and parasite individuals. Richness analyses were restricted to *C. guineensis*, *G. nigri*, *M. sebae*, and *S. melanotheron*, as all other host species were infected by only a single macroparasite species and therefore did not yield biologically meaningful richness estimates.
